# Chromosome evolution and the genetic basis of agronomically important traits in greater yam

**DOI:** 10.1101/2021.04.14.439117

**Authors:** Jessen V. Bredeson, Jessica B. Lyons, Ibukun O. Oniyinde, Nneka R. Okereke, Olufisayo Kolade, Ikenna Nnabue, Christian O. Nwadili, Eva Hřibová, Matthew Parker, Jeremiah Nwogha, Shengqiang Shu, Joseph Carlson, Robert Kariba, Samuel Muthemba, Katarzyna Knop, Geoffrey J. Barton, Anna V. Sherwood, Antonio Lopez-Montes, Robert Asiedu, Ramni Jamnadass, Alice Muchugi, David Goodstein, Chiedozie N. Egesi, Jonathan Featherston, Asrat Asfaw, Gordon G. Simpson, Jaroslav Doležel, Prasad S. Hendre, Allen Van Deynze, Pullikanti Lava Kumar, Jude E. Obidiegwu, Ranjana Bhattacharjee, Daniel S. Rokhsar

**Author notes:** Corresponding authors: Jude E. Obidiegwu < >, Ranjana Bhattacharjee < >, Daniel S. Rokhsar < >. These authors contributed equally: Jessen V. Bredeson, Jessica B. Lyons.

## Abstract

The nutrient-rich tubers of the greater yam *Dioscorea alata* L. provide food and income security for millions of people around the world. Despite its global importance, however, greater yam remains an “orphan crop.” Here we address this resource gap by presenting a highly-contiguous chromosome-scale genome assembly of greater yam combined with a dense genetic map derived from African breeding populations. The genome sequence reveals an ancient lineage-specific genome duplication, followed by extensive genome-wide reorganization. Using our new genomic tools we find quantitative trait loci for susceptibility to anthracnose, a damaging fungal pathogen of yam, and several tuber quality traits. Genomic analysis of breeding lines reveals both extensive inbreeding as well as regions of extensive heterozygosity that may represent interspecific introgression during domestication. These tools and insights will enable yam breeders to unlock the potential of this staple crop and take full advantage of its adaptability to varied environments.

## Introduction

Yams (genus *Dioscorea*) are an important source of food and income in tropical and subtropical regions of Africa, Asia, the Pacific, and Latin America, contributing more than 200 dietary calories per capita daily for around 300 million people^1^. Yam tubers are rich in carbohydrates, contain protein and vitamin C, and are storable for months after harvesting, so they are available year-round^2,3^. World annual production of yam in 2018 was estimated at 72.6 million tons (FAOSTAT 2020). Over 90% of global yam production comes from the ‘yam belt’ (Nigeria, Benin, Ghana, Togo, and Cote d’Ivoire) in West Africa, where yam’s importance is demonstrated by its vital role in traditional culture, rituals, and religion^3–5^. While yams are primarily dioecious, and hence obligate outcrossers, they are vegetatively propagated, allowing genotypes with desirable qualities (disease resistance, cooking quality, nutritional value) to be maintained over subsequent planting seasons.

Greater yam (*Dioscorea alata* L.), also called water yam, winged yam, or ube, among other names, is the species with the broadest global distribution^1^. *D. alata* is thought to have originated in Southeast Asia and/or Melanesia^2,6^. It was introduced to East Africa as many as 2,000 years ago and reached West Africa by the 1500s^2,7^. Several traits of *D. alata* make it particularly valuable for economic production and an excellent candidate for systematic improvement. It is adapted to tropical and temperate climates, has a relatively high tolerance to limited-water environments, and no other yam comes close for yield in terms of tuber weight. *D. alata* is easily propagated, its early vigor prevents weeds, and its tubers have high storability^8^. The tubers of *D. alata* possess high nutritional content relative to other *Dioscorea* spp^9,10^.

Over the last two decades, global yam production has doubled, but these increases have predominantly been achieved through the expansion of cultivated areas rather than increased productivity^1^ (FAOSTAT 2020). To meet the demands of an ever-growing population and tackle the threats that constrain yam production, the rapid development of improved yam varieties is urgently needed^11^. Conventional breeding for desired traits in greater yam is arduous, however, due to its long growth cycle and erratic flowering, and further complicated by the polyploidy common in this species^12–14^. Efforts are currently underway by breeders to develop greater yam varieties with improved yield, resistance to pests and diseases, and tuber quality consistent with organoleptic preferences such as taste, color, and texture^11^. A critical challenge for *D. alata* is its high susceptibility to the foliar disease anthracnose, caused by the fungal pathogen *Colletotrichum gloeosporioides* Penz. Anthracnose disease is characterized by leaf necrosis and shoot die-back, and can cause losses of over 80% of production^15–18^. Anthracnose disease affects *D. alata* more than other domesticated yams; moderate resistance to this disease is present, however, in *D. alata* landraces and breeder’s lines^19,20^.

High-quality genomic resources and tools can facilitate rapid breeding methods for greater yam improvement with huge potential to impact food and nutritional security, particularly in Africa. Here we describe a chromosome-scale reference genome for *D. alata* and a dense 10k marker consensus genetic linkage map from five populations involving seven distinct parental genotypes. Comparison of the *D. alata* reference genome with the recently sequenced genomes of the distantly related *D. rotundata*^*21*^ and *D. zingiberensis*^*22*^ reveals substantial conservation of chromosome structure between *D. alata* and *D. rotundata* but considerable rearrangement relative to more deeply divergent *D. zingiberensis* lineage. Analysis of the *D. alata* genome supports the existence of ancient polyploidy events shared across Dioscoreales, and reveals chromosome rearrangements after the most recent pan-Dioscoreales genome duplication. We use genomic and genetic resources to identify nine QTL for anthracnose resistance and tuber quality traits. Our dense multi-parental genetic map complements the maps previously used for QTL mapping for anthracnose resistance^23–25^ and sex determination^26^. These tools and resources will empower breeders to use modern genetic tools and methods to breed the crop more efficiently, thereby accelerating the release of improved varieties to farmers.

## Results and Discussion

### I. Genome sequence and structure

We generated a high-quality reference genome for *D. alata* by assembling whole-genome shotgun sequence data from PacBio single-molecule continuous long reads (234× coverage in reads with 15.1 kb N50 read length), with short-read sequencing for polishing and additional mate-pair linkage (see **Methods, Table 1, Supplementary Note 1, Supplementary Data 1**). High-throughput chromatin conformation contact (HiC) data and a composite meiotic linkage map (see below) were used to organize the contigs (N50 length 4.5 Mb) into n=20 chromosome-scale sequences, matching the observed karyotype, with each pair of homologous chromosomes represented by a single haplotype-mosaic sequence (**Supplementary Figs. 1** and **2**). The genome assembly spans a total of 479.5 Mb, consistent with estimates of 455 ± 39 Mb by flow cytometry^13^, and 477 Mb by k-mer-based analyses (**Table 1, Supplementary Note 1**). The chromosome-scale ‘version 2’ assembly is available via YamBase (ftp://yambase.org/genomes/Dioscorea_alata) and Phytozome (https://phytozome-next.jgi.doe.gov/info/Dalata_v2_1), replacing the early ‘version 1’ draft released in those databases in 2019.

**Table 1.**
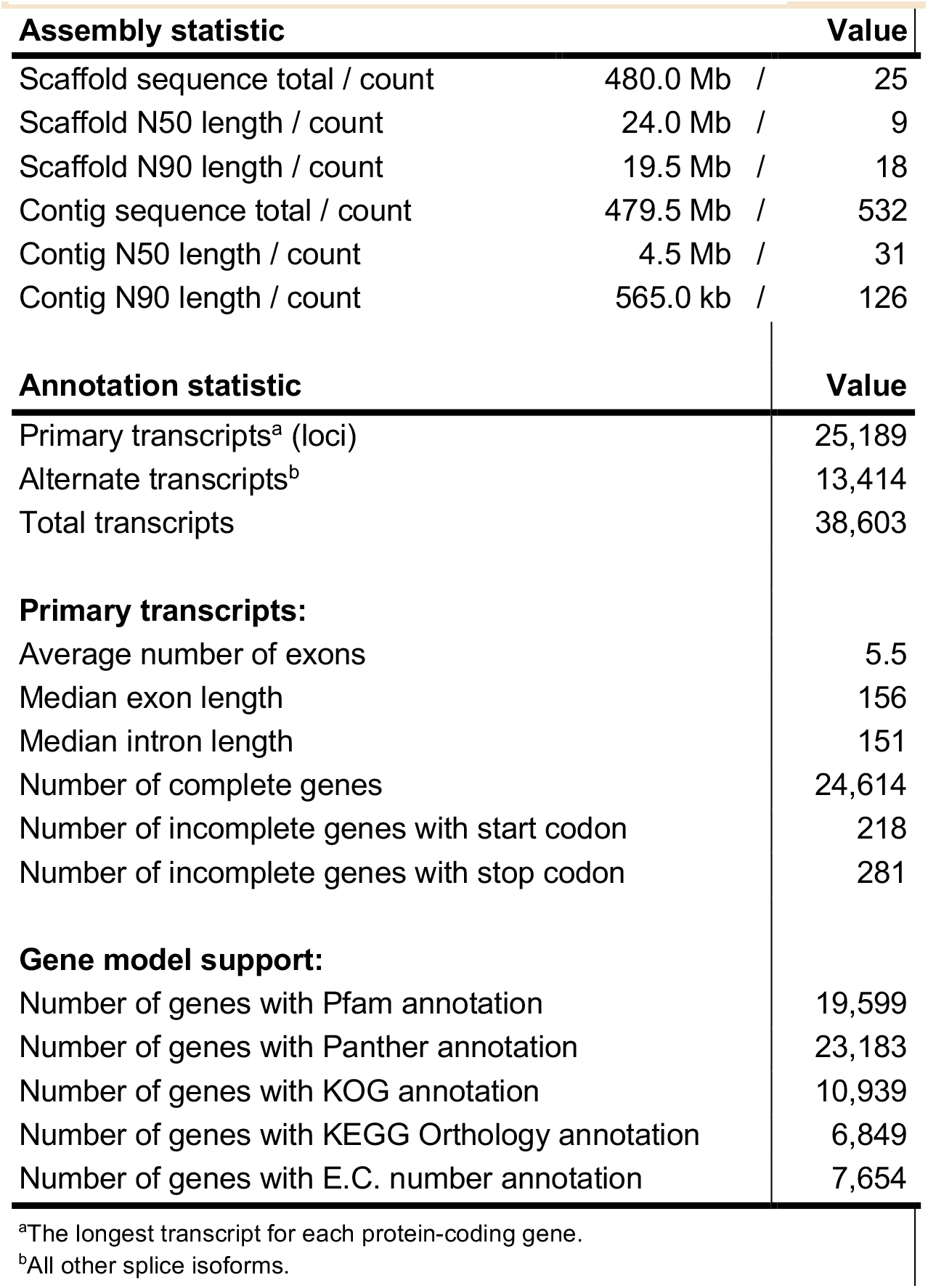
Assembly and Annotation Statistics.

The genomic reference genotype, TDa95/00328, is a breeding line from the Yam Breeding Unit of the International Institute of Tropical Agriculture (IITA), Ibadan, Nigeria. It is moderately resistant to anthracnose^23,27^ and has been used as a parent frequently in crossing programs. TDa95/00328 is diploid with 2n=2x=40, as confirmed by chromosome counting and DNA flow cytometry (**Supplementary Fig. 2**; Gatarira et al., in preparation) and genetically by segregation of AFLP^23^. The reference accession exhibits long runs of homozygosity due to inbreeding (**Supplementary Fig. 3**); outside of these segments we observe 7.9 heterozygous sites per kilobase.

To corroborate our genome assembly and provide tools for genetic analysis, we generated genetic linkage maps from nine mapping populations that involved seven distinct parents segregating for relevant phenotypic traits (**Table 2, Supplementary Table 1**; see also **Fig. 4**). These mapping populations were generated from biparental crosses performed at IITA, with 32– 317 progeny per cross. Genotyping was performed using sequence tags generated with DArTseq (Diversity Arrays Technology Pty), mapped to the genome assembly, and filtered (**Methods, Supplementary Note 2, Supplementary Data 2**), producing 13,584 biallelic markers that segregate in at least one of our mapping populations (**Supplementary Table 2**).

**Table 2.**
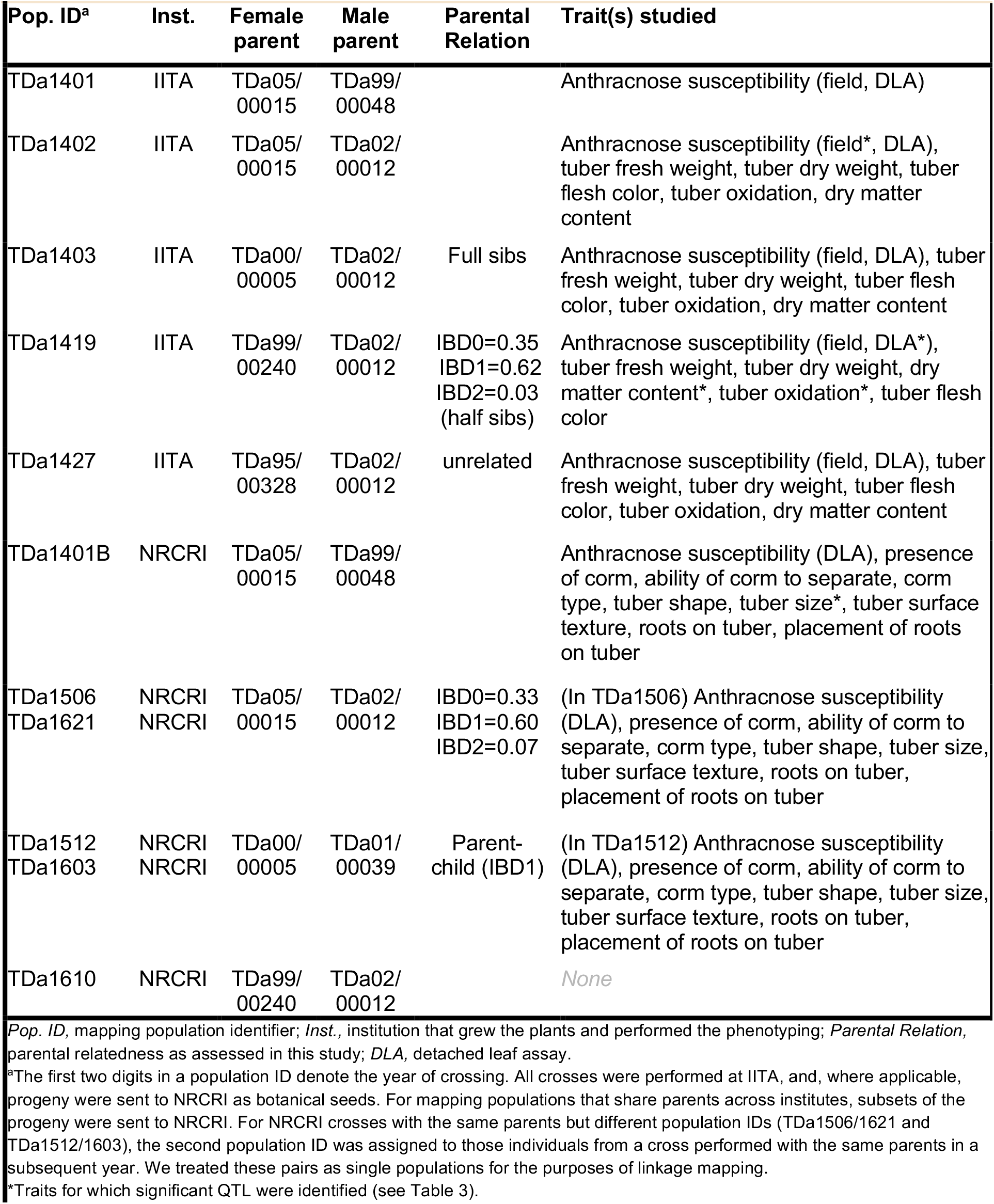
Mapping populations used in this study.

| Pop. ID<sup>a</sup> | Inst. | Female parent | Male parent | Parental Relation | Trait(s) studied |
| --- | --- | --- | --- | --- | --- |
| TDa1401 | IITA | TDa05/00015 | TDa99/00048 |  | Anthrachnose susceptibility (field, DLA) |
| TDa1402 | IITA | TDa05/00015 | TDa02/00012 |  | Anthrachnose susceptibility (field*, DLA), tuber fresh weight, tuber dry weight, tuber flesh color, tuber oxidation, dry matter content |
| TDa1403 | IITA | TDa00/00005 | TDa02/00012 | Full sibs | Anthrachnose susceptibility (field, DLA), tuber fresh weight, tuber dry weight, tuber flesh color, tuber oxidation, dry matter content |
| TDa1419 | IITA | TDa99/00240 | TDa02/00012 | IBD0=0.35<br>IBD1=0.62<br>IBD2=0.03 (half sibs) | Anthrachnose susceptibility (field, DLA*), tuber fresh weight, tuber dry weight, dry matter content*, tuber oxidation*, tuber flesh color |
| TDa1427 | IITA | TDa95/00328 | TDa02/00012 | unrelated | Anthrachnose susceptibility (field, DLA), tuber fresh weight, tuber dry weight, tuber flesh color, tuber oxidation, dry matter content |
| TDa1401B | NRCRI | TDa05/00015 | TDa99/00048 |  | Anthrachnose susceptibility (DLA), presence of corm, ability of corm to separate, corm type, tuber shape, tuber size*, tuber surface texture, roots on tuber, placement of roots on tuber |
| TDa1506<br>TDa1621 | NRCRI<br>NRCRI | TDa05/00015 | TDa02/00012 | IBD0=0.33<br>IBD1=0.60<br>IBD2=0.07 | (In TDa1506) Anthrachnose susceptibility (DLA), presence of corm, ability of corm to separate, corm type, tuber shape, tuber size, tuber surface texture, roots on tuber, placement of roots on tuber |
| TDa1512<br>TDa1603 | NRCRI<br>NRCRI | TDa00/00005 | TDa01/00039 | Parent-child (IBD1) | (In TDa1512) Anthrachnose susceptibility (DLA), presence of corm, ability of corm to separate, corm type, tuber shape, tuber size, tuber surface texture, roots on tuber, placement of roots on tuber |
| TDa1610 | NRCRI | TDa99/00240 | TDa02/00012 |  | None |

*Pop. ID*, mapping population identifier; *Inst.*, institution that grew the plants and performed the phenotyping; *Parental Relation*, parental relatedness as assessed in this study; *DLA*, detached leaf assay.
<sup>a</sup>The first two digits in a population ID denote the year of crossing. All crosses were performed at IITA, and, where applicable, progeny were sent to NRCRI as botanical seeds. For mapping populations that share parents across institutes, subsets of the progeny were sent to NRCRI. For NRCRI crosses with the same parents but different population IDs (TDa1506/1621 and TDa1512/1603), the second population ID was assigned to those individuals from a cross performed with the same parents in a subsequent year. We treated these pairs as single populations for the purposes of linkage mapping.
\*Traits for which significant QTL were identified (see Table 3).

The 20 linkage groups derived from individual maps corroborated the sequence-based genome assembly and were particularly useful for interpreting HiC linkage between telomeres and determining the correct intra-chromosomal orientations of the arms. These features were difficult to organize using HiC alone, due to strong Rabl conformations (**Fig. 1a**, and **Supplementary Figs. 1** and **4**) that led to contacts between the distal regions of chromosome arms (see below). The ten genetic maps were highly concordant (**Fig. 1b**; Kendall’s tau correlation coefficients = 0.9091–0.9626), and we combined them into a single composite linkage map using five maps that capture the genetic diversity of the seven distinct parents **(Supplementary Table 2, Supplementary Data 3**). The composite map spans 1,817.9 centimorgans, accounting for a total of 2,178 meioses (1,089 individuals), and includes 10,448 well-ordered (Kendall’s tau = 0.9989; **Supplementary Fig. 5**) markers (excluding markers genotyped in individual crosses that were discordant post-imputation and/or were not phaseable) (**Methods, Supplementary Note 2**). This is the highest resolution genetic linkage map for *D. alata* produced to date.

**Fig 1.**
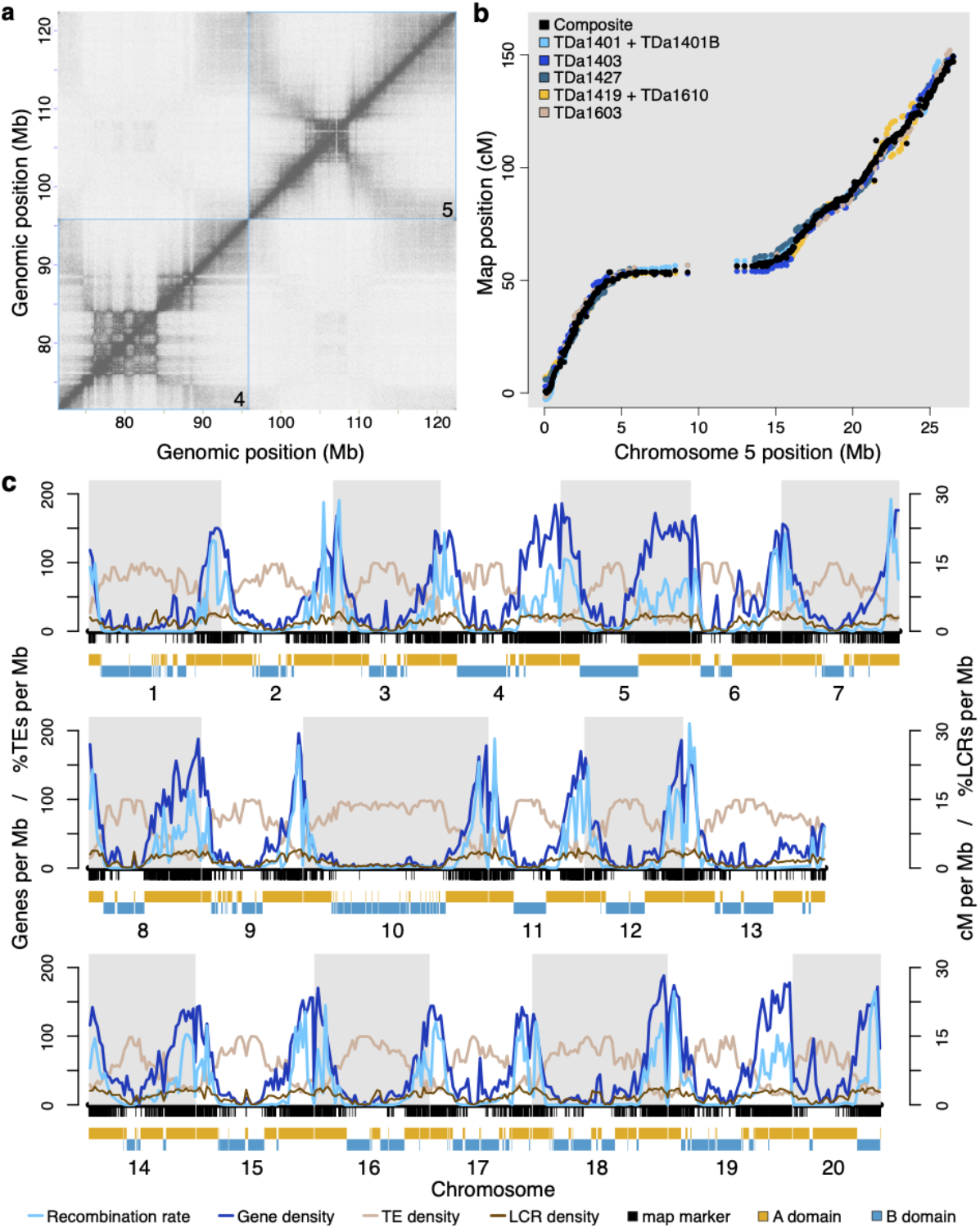
*D. alata* genome structure and recombination. **a** HiC contact matrix of TDa95/00328 chromosomes 4 and 5. Within chromosomes, the band of high contact density along the diagonal reflects the well-ordered underlying assembly. The “checkerboard” pattern observed between 75–85 Mb indicates chromatin domain A/B compartmentalization^28^ within chromosome 4. The ‘winged’ pattern observed within chromosomes, particularly chromosome 5, showing elevated contact densities between chromosome ends is typical of Rabl-structured chromosomes in the nucleus^29^. Chromosomes are outlined with cyan boxes. Each pixel represents the intersection between a pair of 50 kb loci along the chromosomes. The density of contacts between two loci is proportional to pixel color, with darker pixels representing more contacts and lighter representing fewer. **b** A composite genetic linkage map (black points), integrating five mapping populations (colored points, legend), is shown for chromosome 5. The maps exhibit highly-concordant marker orders (Kendall’s tau correlations between 0.9091 and 0.9626) and validate the large-scale correctness of the chromosome-scale assembly. The sigmoidal shape of the maps along the physical chromosome reflects suppressed recombination within the pericentromere. Individual component maps were scaled and shifted vertically to display their marker-order concordance. **c** The *D. alata* chromosome landscape is shown. Transposable elements (TEs; tan lines, left Y-axis) are enriched within the pericentromeres; while low-complexity repeat (LCR; brown, right Y-axis), protein-coding gene (dark blue line, left Y-axis), and meiotic recombination (cyan lines, right Y-axis) densities are elevated nearer the chromosome ends. Composite map markers are shown as black ticks under the X-axis, with A/B chromatin compartment structure drawn below (‘A’ compartment domains in gold and ‘B’ domains in dark cyan).

The *D. alata* reference genome encodes an estimated 25,189 protein-coding genes, based on an annotation that took advantage of both existing and new *D. alata* transcriptome resources as well as interspecific sequence homology (**Table 1, Methods, Supplementary Note 3**). With a benchmark set of embryophyte genes^30,31^, we estimate that the *D. alata* gene set is 97.8% complete, with 1.5% gene fragmentation. While BUSCO methodology suggests that only 0.7% of the genes are missing, this is an overestimate, since some of these nominally-missing genes are detected by more sensitive searches (**Supplementary Note 3**). Our new transcriptome datasets include short-read RNAseq as well as 626,000 long, single-molecule direct-RNA sequences from twelve TDa95/00328 tissues. The transcriptome data identified 13,414 alternative transcripts. The great majority of genes have functional assignments through Pfam (n=19,599) and Panther (n=23,183) (**Table 1**).

Within chromosomes, protein-coding gene and transposable element densities are strongly anticorrelated (Pearson’s *r* = −0.885), with gene loci concentrated in the highly-recombinogenic distal chromosome ends (Pearson’s *r* = +0.823) and transposable elements, particularly Ty3/metaviridae and Ty1/pseudoviridae LTRs and other unclassified repeats, are enriched in the recombination-poor pericentromeres (Pearson’s *r* = −0.718) (**Fig. 1d, Supplementary Fig. 6, Supplementary Table 3**). Homopolymers and simple-sequence repeats, however, were positively correlated with gene (Pearson’s *r* = +0.838) and recombination (Pearson’s *r* = +0.728) densities.

Analysis of chromatin conformation capture (HiC) data reveals the structure of interphase chromosomes in greater yam (**Methods, Supplementary Note 4**). We find that all chromosomes adopt a ‘Rabl’-in-bouquet conformation (**Supplementary Fig. 4**) in which each chromosome appears “folded” in the vicinity of the centromere, as (1) chromatin contacts are enriched among chromosome ends and (2) these chromosome ends are depleted of contacts with the pericentromeres (see also refs.^29,32^) Greater yam chromosomes also show alternating ‘A/B’ chromatin compartmentalization, as is demonstrated in several other plant species^33^. In greater yam, the gene-rich distal regions of each chromosome are generally spanned by open ‘A’ domains (between gene density and A/B domain status, Pearson’s *r* = +0.628), while the relatively gene-poor and transposon-rich pericentromeres are characterized by closed ‘B’ domains that are often punctuated by smaller ‘A’ domains (**Supplementary Fig. 7**).

### II. Comparative analysis and paleopolyploidy

Comparison of the greater yam genome and protein-coding annotation with those of white yam (*D. rotundata*, also known as Guinea yam), bitter yam (*D. dumetorum*), and peltate yam (*D. zingiberensis*) highlights the completeness of our sequence and annotation and the extensive sequence divergence across the genus. Among the *Dioscorea* species sequenced to date, the annotation of *D. alata* appears to be the most complete (**Supplementary Note 3**). For example, *D. alata* has the fewest missing conserved gene families in cross-species comparisons within Dioscoreaceae (53 in *D. alata* compared with 385 for *D. zingiberensis* and 595 for *D. rotundata*) and in cross-monocot comparisons (7 in *D. alata* compared with 99 in *D. zingiberensis* and 110 in *D. rotundata*) (**Supplementary Fig. 6**). These metrics combine genome assembly completeness and accuracy with exon-intron structure predictions based, in part, on transcriptome resources (see also **Supplementary Table 4**).

At the nucleotide level, *D. alata* coding sequences exhibit 97.4%, 93.6%, and 86.5% identity with *D. rotundata, D. dumetorum*, and *D. zingiberensis*, corresponding to median synonymous substitution rates (Ks) of 0.064, 0.163, and 0.389, respectively. These measures are consistent with *D. zingiberensis* being a deeply branching outgroup to the clade formed by *D. alata, D. rotundata*, and *D. dumetorum* (see also **Supplementary Table 5**), and highlights the ∼60 My old divergences within the genus *Dioscorea*. The medicinal plant *Trichopus zeylanicus* (common name ‘Arogyappacha’ in India, meaning “the green that gives strength”)^34^ is a more distantly-related member of the Dioscoreaceae family, with 77.9% identity and median Ks of 0.804.

The chromosomes of *D. alata* and *D. rotundata* are in 1:1 correspondence, and are highly collinear (**Fig. 2a, Supplementary Fig. 8a**). The few intra-chromosome differences observed could represent *bona fide* rearrangements between species or, possibly, imperfections in the *D. rotundata* v2 assembly^21^ that could have arisen from the reliance on linkage mapping to order and orient *D. rotundata* scaffolds, especially in recombination-poor pericentromeric regions of the genome. Under the assumption that *D. rotundata* chromosomes are in 1:1 correspondence with *D. alata* chromosomes, we can provisionally assign four large but unmapped *D. rotundata* scaffolds to chromosomes (**Fig. 2a**). We found one inter-chromosome difference (not present in the earlier *D. rotundata* assembly), which requires further study (**Supplementary Fig. 8a**).

**Fig 2.**
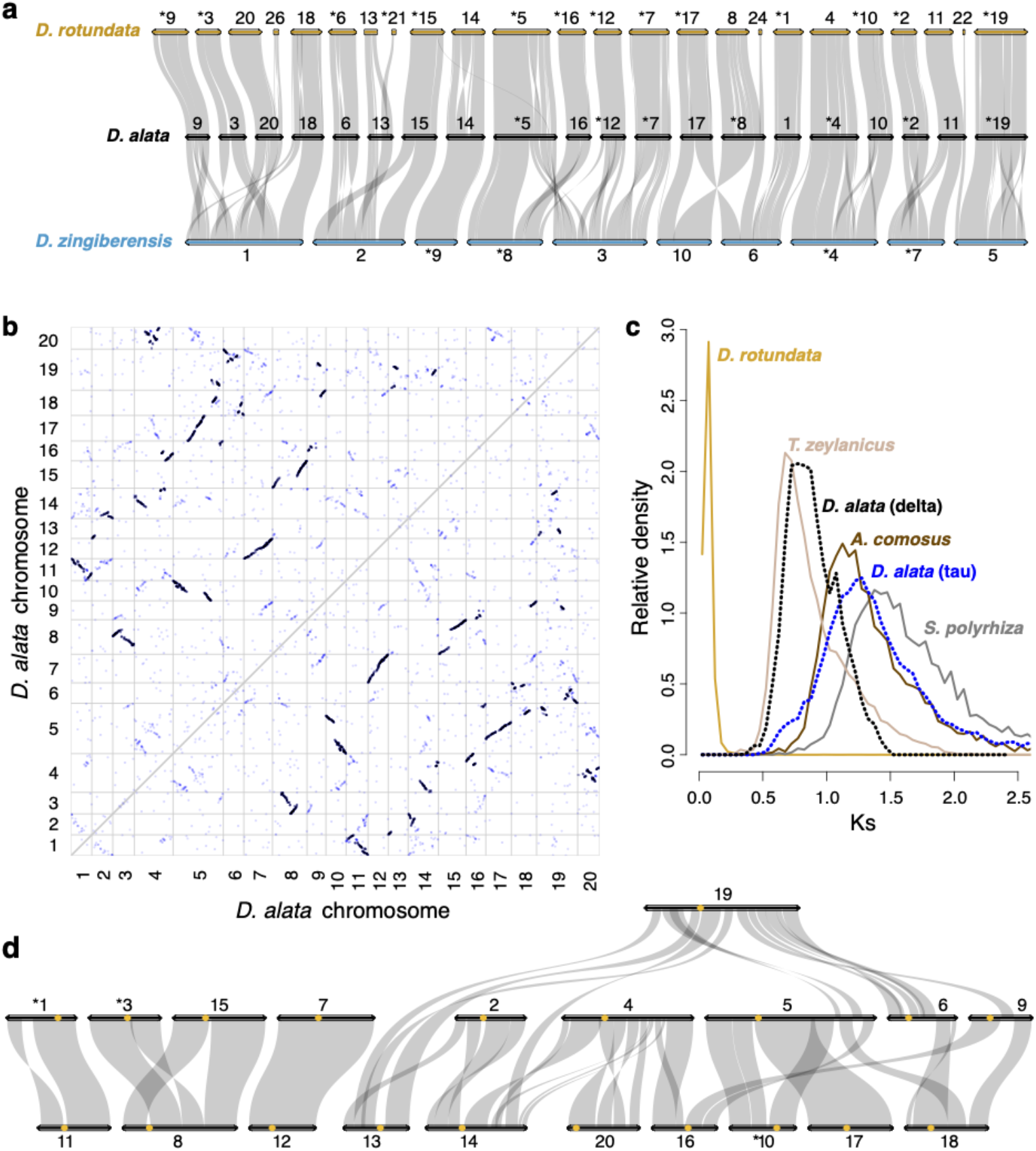
Dioscoreaceae chromosome evolution. **a** Ribbon diagram demonstrating conserved chromosomal synteny and large-scale segmental collinearity (semi-transparent grey ribbons) between *D. alata* (black horizontal bars), *D. rotundata* (gold), and *D. zingiberensis* (cyan) one-to-one orthologous gene pairs. Only *D. rotundata* sequences with five or more collinear genes are shown. To improve visual clarity, some chromosomes, marked with asterisks, were reverse complemented with respect to their assembled sequences. Chromosome sizes are proportional to the number of annotated genes. **b** Dot plot showing evidence of two whole-genome duplications exposed by TDa95/00328 intragenomic comparison. Each point represents a homoeologous gene pair and each white box (outlined in grey) represents the intersection of two chromosomes. Homoeology from the recent Dioscoreaceae ‘delta’ duplication is shown in black and the ancient, core monocot ‘tau’ duplication can be seen in blue. **c** The synonymous substitution rate (Ks) histograms for orthologous (solid lines) or homoeologous (dotted lines) gene pairs between *D. alata* and select species comparators are shown. The *D*.*alata*– *D*.*rotundata* ortholog density was rescaled by 0.25 to emphasize other comparisons. **d** Shared segmental homoeology (semi-transparent grey) between *D. alata* chromosomes (black horizontal bars) resulting from the delta duplication is depicted with a ribbon diagram, as in panel **a**, but with centromeres positions now included as gold circles.

While the draft *D. dumetorum* genome assembly is not organized into chromosomes, comparison with the *D. alata* reference shows that the two genomes are locally collinear on the scale of the *D. dumetorum* contigs, with only one discordance (**Supplementary Fig. 8b**). This observation suggests a provisional organization of the *D. dumetorum* genome into probable chromosomes. Notably, the distantly related *D. zingiberensis* has a haploid complement of n=10, compared with n=20 found in *D. alata, D. rotundata*, and *D. dumetorum*. We find that the *D. zingiberensis* chromosomes^22^ were formed from ancestral, *D. alata*-like chromosomes and/or chromosome arms by combinations of end-to-end and centric fusions and translocations (**Fig. 2a, Supplementary Fig. 8c**).

We found evidence for two ancient paleotetraploidies in the *D. alata* lineage. These duplications evidently preceded the origin of the genus, since *Dioscorea* genomes show one-to-one orthology (**Supplementary Fig. 8**). The most recent paleotetraploidy is apparent from extensive collinear paralogy in *D. alata* (**Fig. 2b**) and coincides with the genome duplication recently described in *D. zingiberensis*^*22,35*^. Following the common use of Greek letters to denote plant polyploidies, we designate this *Dioscorea* lineage duplication as ‘delta.’ The median sequence divergence between 1,578 delta paralogs in *D. alata* is Ks = 0.869 substitutions/site (**Fig. 2c**). While comparisons with the draft genome assembly of *T. zeylanicus* (Ks = 0.804 to *D. alata*) further suggest that the delta paleotetraploidy may have preceded the origin of the family Dioscoreaceae, the fragmentation of the *T. zeylanicus* assembly precludes a definitive assessment. The timing of the delta duplication (estimated to be 64 Mya^22^) is contemporaneous with the K/T boundary and a cluster of successful paleopolyploidies^36^.

Analysis of the *D. alata* genome reveals large-scale genomic reorganization after the delta duplication. *D. alata* chromosomes preserve long collinear paralogous segments arising from the delta paleotetraploidy event, and the genomic organization of these segments reveals large-scale rearrangements after whole-genome duplication (**Fig. 2d, Supplementary Data 4**). These include cases of one-to-one full chromosome paralogs, (chromosomes 1 and 11; 7 and 12) as well as examples of centric insertion (e.g., the paralog of chromosome 3 was inserted within the paralog of chromosome 15 to form chromosome 8; the paralog of chromosome 17 was inserted into the paralog of chromosome 10 to form most of the chromosome 5). Other large-scale rearrangements are evident, including apparent end-to-end ‘fusions’ (or more properly translocations^37^). Taken together, these paralogies provide further evidence for the delta duplication.

In addition to delta, the *D. alata* genome also displays relicts of a more ancient genome-wide duplication in the form of nearly collinear ancient paralogous segments with median Ks = 1.21 substitutions per site (**Fig. 2b**,**c**). We identify this duplication with the famed ‘tau’ duplication shared by other core monocots including grasses^38^, pineapple (*Ananas comosus*^*39*^), oil palm (*Elaeis guineensis*^*40*^), and asparagus (*Asparagus officinalis*^*41*^) but not duckweed *Spirodela polyrhiza*^*42*^. The clear 2:2 pattern of orthology between yam, pineapple, and oil palm (**Supplementary Fig. 8**) confirms that these three lineages have each experienced one lineage-specific whole-genome duplication (delta, sigma, and p, respectively) since they diverged from each other. This pattern implies that relicts of any earlier duplications observed in these species must represent shared events. Since Dioscoreales is one of the earliest branching core monocots (only Petrosaviales branches earlier), the discovery of tau in yam implies that this duplication likely preceded the divergence of the core monocot clade (**Supplementary Fig. 9**).

### III. QTL mapping

To demonstrate the utility of our dense linkage map and high-quality *D. alata* reference genome for advancing greater yam breeding, we searched for quantitative trait loci (QTL) for resistance to anthracnose disease and several tuber quality traits (dry matter, oxidation, tuber color, corm type, and other traits). Our mapping populations were generated in controlled crosses by yam breeders at IITA, Nigeria, using parents from the yam breeding program (**Table 2, Supplementary Table 1**). Phenotyping was performed in Nigeria at IITA Ibadan and NRCRI in Umudike (**Methods, Supplementary Note 5**). Leveraging the ability to clonally propagate individuals, we measured multiple traits over the years 2016–2019. Our QTL analyses exploited the imputed genotypes derived from our dense linkage maps. In total, we found eight distinct QTL: three for anthracnose resistance and five for tuber traits (**Fig. 3, Table 3, Supplementary Figs. 10–11**).

**Fig 3.**
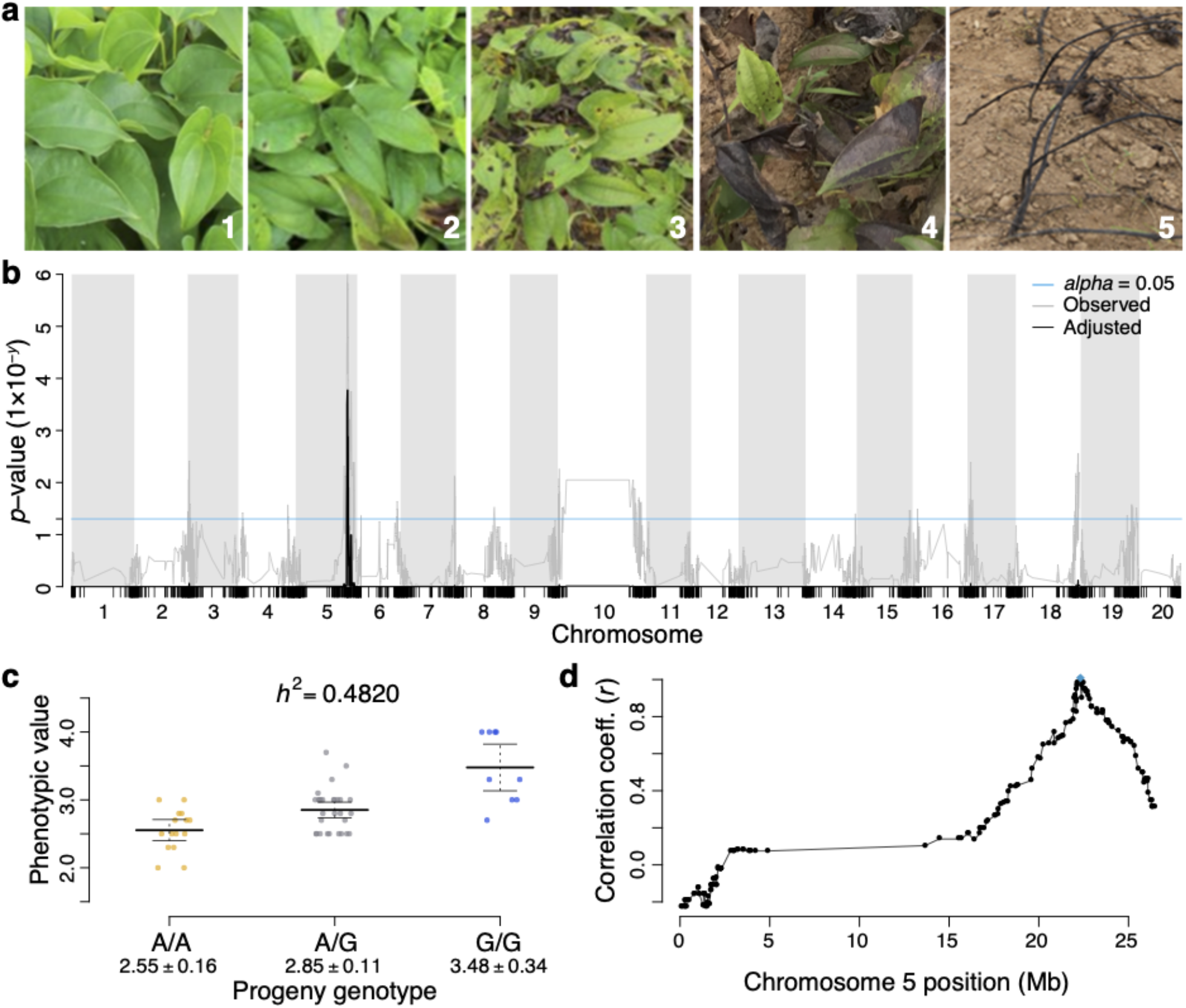
Quantitative trait locus for anthracnose resistance. **a** Exemplars of the yam anthracnose disease (YAD) field assessment severity rating (scored on a 1–5 scale) used at IITA in Ibadan, Nigeria. **b** QTL association scan for YAD resistance in the TDa1402 genetic population for year 2017. The family-wise max(*T*)-corrected association significance values for each genotyped locus is represented by the black line, and the uncorrected significance by the grey line. The minimum threshold for significance (*alpha* = 0.05) is represented as a cyan horizontal line. **c** An effect plot for the peak locus on chromosome 5 at 23.3 Mb, explaining a large proportion of the phenotypic variance (i.e., narrow-sense heritability, *h*^2^) observed. The alleles and genotypes are shown along the X-axis with their phenotypic values (horizontal lines mark the mean ± 95% confidence interval). **d** Plot showing the strength of linkage disequilibrium (LD) between the peak marker (cyan diamond) and other loci (black points) in chromosome 5. LD was calculated as Pearson’s correlation (*r*).

**Table 3.**
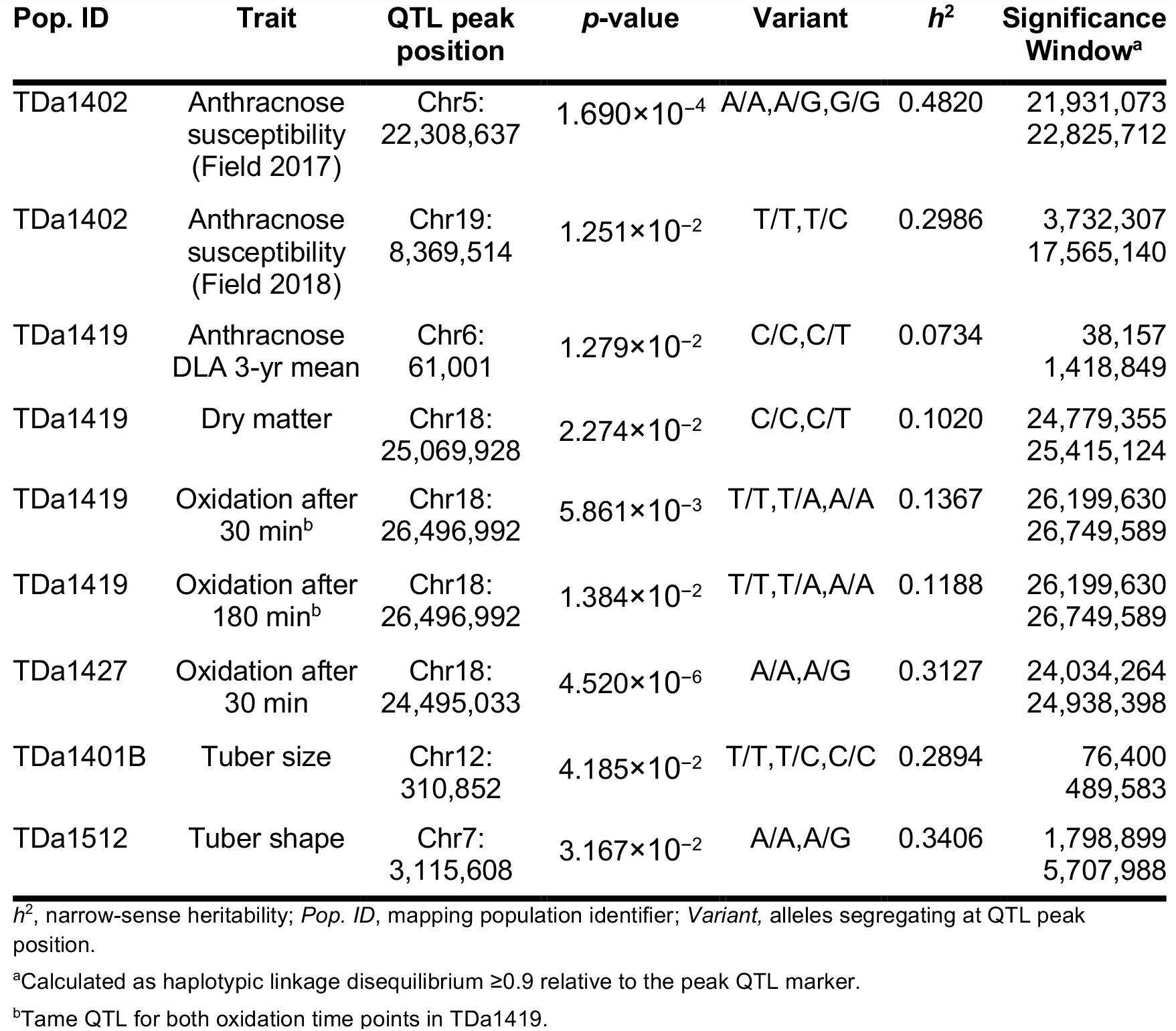
Significant QTL identified in this study.

#### Anthracnose resistance

Yam Anthracnose Disease (YAD), or yam dieback, is a major disease of yams caused by the fungus *Colletotrichum gloeosporioides*^*15,18*^. *D. alata* is particularly susceptible to YAD, although resistance has been shown to vary among *D. alata* genotypes^43^. We sought QTL for YAD resistance using field trials in five mapping populations and detached leaf assays in eight mapping populations (**Table 2, Methods, Supplementary Note 5**). While most of these populations did not show significant QTL, we found three significant anthracnose resistance QTL in two of them.

In field trials of the TDa1402 population, we found a major QTL on chromosome 5 (*p* = 1.7×10^−4^) that explains 48.2% of phenotypic variance in the 2017 data, with an additive effect (**Fig. 3a–c**), and a minor QTL on chromosome 19 (**Supplementary Fig. 10a–c)** that explains 29.9% of the variance in the 2018 data (*p* = 1.25×10^−2^). These QTL are candidates for use in marker-assisted breeding. However, since variation in levels of infestation, overall plant vigor, and timing and amount of rainfall influence disease severity in field trials, validation of these QTL is required.

In detached leaf assays of the TDa1419 population, performed under varying conditions over three years (**Methods**), we found a QTL of smaller effect (7.3% of phenotypic variance) on chromosome 6 (**Supplementary Fig. 10d–f**). While this QTL was significant (*p* = 1.279×10^−2^), it was found only using three-year averages, and the locus was not significantly associated with YAD in the data from individual years. Furthermore, anthracnose disease levels, as measured by detached leaf assay, were not significantly correlated across genotypes among years. These observations suggest that variation in YAD may be dominated by non-genetic factors.

#### Tuber quality traits

Post-harvest oxidation causes browning of yam tuber flesh and flavor changes that reduce crop value^44^. We found an additive-effect QTL for tuber oxidation after peeling at both 30 min (*p* = 5.861×10^−3^) and 180 min (*p* = 1.384×10^−2^) on chromosome 18 in the TDa1419 population (**Supplementary Fig. 11a–f**). The QTL explained 13.67% and 11.88% of the phenotypic variance at 30 and 180 minutes after peeling, respectively. In the TDa1427 population, a closely linked QTL (*p* = 4.520×10^−6^) located 2 Mb upstream on the same chromosome explained 31.3% of the phenotypic variance in oxidation after 30 minutes (**Supplementary Fig. 11g–i**). Although enzymatic browning in yam remains poorly understood, polyphenol oxidase and peroxidase are active during browning of *D. alata* and *D. rotundata*^*45*^, and inhibition of this activity has been shown to reduce browning in Chinese yam (*D. polystachya*)^46^. We find a cluster of three peroxidases on chromosome 18 at 26.23–26.36 Mb, within ∼200 kb of the oxidation QTL at 26.50 Mb in TDa1419 and within 2 Mb of the oxidation QTL in TDa1427, raising the possibility that oxidation is affected by genetic variation in peroxidase activity.

Dry matter (principally starch) content is an important measure of yam yield^47^. We found a single, minor QTL (explaining 10.2% of the phenotypic variance for dry matter) on chromosome 18 (**Supplementary Fig. 11j–l**) in population TDa1419 at position Chr18:25,069,928 (*p* = 2.274×10^−2^). The genotypes observed at this QTL are segregating in the population in a pseudo-testcross configuration. Lastly, we identified two QTL for tuber size (*p* = 4.185×10^−2^) and shape (*p* = 3.167×10^−2^) in populations TDa1401B and TDa1512, respectively, accounting for 28.9% and 34.1% of their phenotypic variances (**Supplementary Fig. 11k–p**).

Several recent studies have identified QTL for anthracnose resistance or tuber quality traits in *D. alata*. Two significant anthracnose QTL were identified using EST-SSRs^25^, and three using GBS-SNPs^48^. Three loci associated with dry matter content and two associated with oxidative browning were identified via a genome-wide association study (GWAS)^49^. We identified six distinct QTL for these traits, and none of them co-localize with the QTL from these earlier studies. This suggests that genotype-by-environment interactions, population structure and size, and/or QTL inference methods may affect which loci are identified in *D. alata* QTL studies. For example, our analysis of the TDa1402 population grown in two different fields in 2017 and 2018 identified anthracnose QTL on different chromosomes.

### IV. Genetic variation within greater yam

To enable future genetic analyses we developed a catalog of over 3.05 million biallelic single-nucleotide variants (SNVs) in *D. alata*, based on whole-genome shotgun resequencing (**Supplementary Note 6, Supplementary Data 1**) of breeding lines representing the seven parents of our biparental mapping populations and an additional breeding line (TDa95-310). Of the 3.05 million biallelic SNVs, in our collection, 1.89 million could be confidently genotyped across all samples. Included within the larger set are 305.5k coding SNVs (251.5k in the reduced set) with predicted effect, 127.1k of which introduce non-synonymous amino acid changes.

We used these dense SNVs to determine the relationships among the eight breeding lines (**Supplementary Tables 1** and **6, Fig. 4a**) by estimating the fractions of their genomes they shared as identical by descent (IBD). We identified six parent-child relationships (i.e., one haplotype shared (IBD1) across the entire genome; relatedness coefficients ∼0.50) and five second-degree relationships (i.e., coefficients of ∼0.25). All second-degree relations showed unusually high values of IBD1, and both first- and second-degree relations shared substantial IBD2, suggesting a history of recent inbreeding. The relationships inferred are consistent with available pedigree records (**Supplementary Table 1**), with the addition of several previously unrecorded grandparent-grandchild relationships. Although the use of highly related parents in breeding programs limits the diversity of alleles available for selection, we note that as a practical matter yam crosses are limited to genotypes that flower appropriately, consistently and profusely.

**Fig 4.**
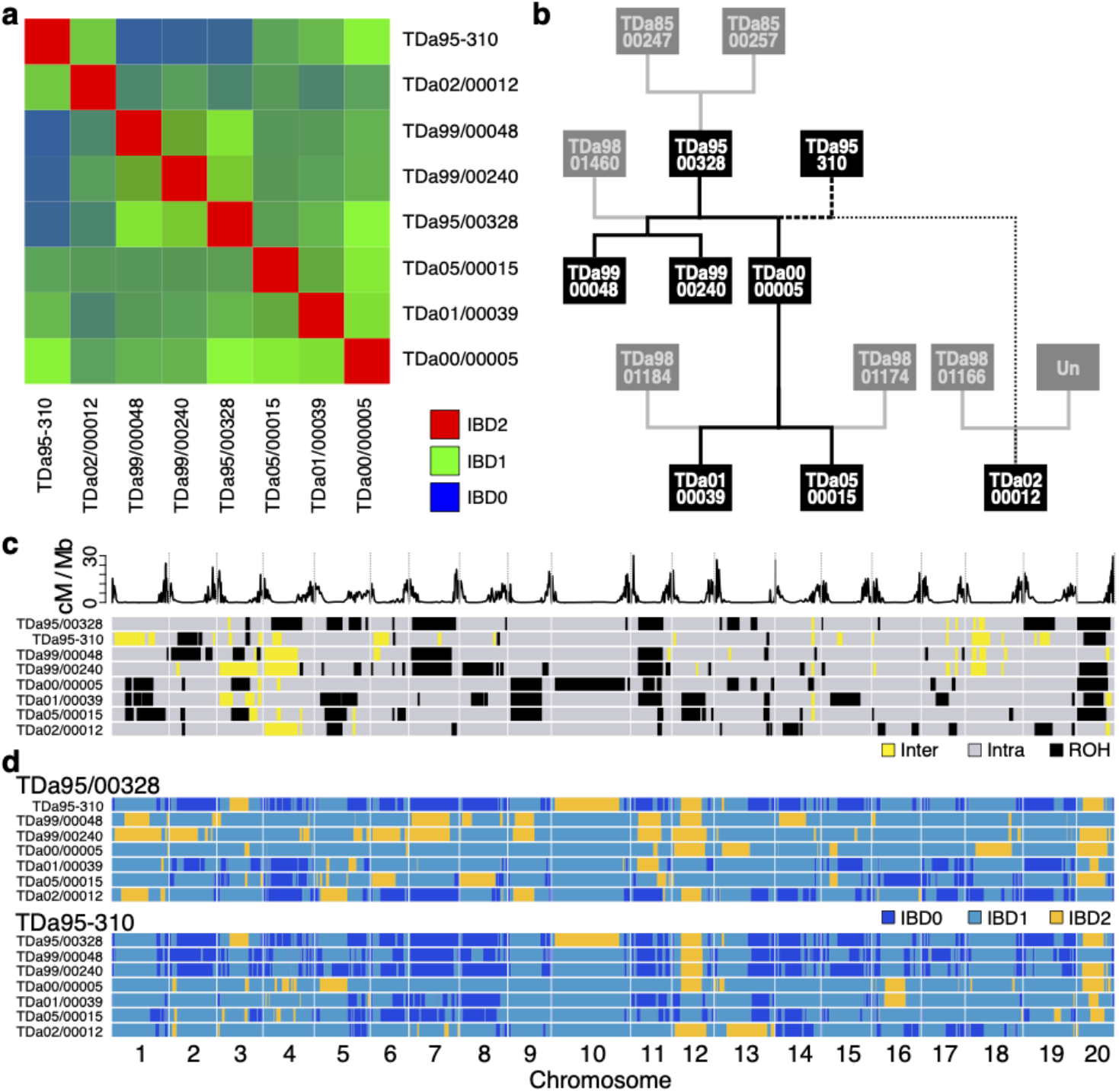
Relationships between eight deeply sequenced *D. alata* breeding lines. **a** Red, green, and blue color-channel matrix of identity-by-descent (IBD) relatedness between all pairs of individuals (see also **Supplementary Table 6**). The red channel represents the degree of genome identity (IBD2), green pairs share one haplotype (IBD1), and pairs in blue share few or no haplotypes (IBD0). **b** Pedigree of relationships. Sequenced individuals are represented in black boxes with white text, while individuals not sequenced are in grey. Relationships known via IITA records are drawn with solid lines (see **Supplementary Table 1**). Relationships that could be confirmed using direct sequence comparison are highlighted with solid black lines, and those that could not be are in grey. Inferred cryptic relationships are indicated with dashed lines (first-degree as bold dashed lines and second-degree relations as thin dotted lines). Unexpectedly, TDa95-310 was revealed to be a parent of TDa00/00005, and a grandparent of TDa02/00012. **c** Regions of heterozygosity, autozygosity, and possible introgression. Within a background of intraspecific genetic variation (grey), large homozygous blocks (runs of homozygosity [ROH], black) appear common in the resequenced individuals, indicating autozygosity from historical inbreeding. In addition, large blocks of exceptionally-high heterozygosity (yellow) can also be observed, indicating possible introgressions (interspecific variation) in one or more of the unsampled founders of the pedigree. Recombination rate along each chromosome is shown in the track above. **d** Haplotype sharing between TDa95/00328 and all other resequenced individuals, and TDa95-310 and all others. Regions of the genome where an individual shares two haplotypes (i.e, they are IBD2) with TDa95/00328 (or TDa95-310) are highlighted in orange. Regions where only one haplotype is shared (i.e., IBD1) are colored in cyan. Regions where both individuals compared do not share any haplotypes (i.e, they are IBD0) are colored dark blue.

Unexpectedly, our identity-by-descent analysis shows that TDa95-310 shares a parent-child relationship to TDa00/00005 and a grandparent-grandchild relationship to TDa01/00039 and TDa05/00015. This finding implies that TDa95-310 and the individual TDa98/00150, which appears in the corresponding position in pedigrees, are clones, or that TDa98/00150 is not a parent of TDa00/00005. TDa95-310 is a landrace from Cote d’Ivoire that is likely derived from an accession known as ‘Brazo-Fuerte’ (‘strong arm’) introduced from Latin America. It is susceptible to anthracnose and has been used as parent material for crossing^50,51^. Since we find that TDa95-310 is a grandparent of TDa02/00012, based on the reported pedigree (**Fig. 4b**), TDa95-310 must also be a parent of either (a) TDa98/01166 or (b) the unknown pollen parent of TDa02/00012. Additional genotyping will resolve this mystery and prevent accidental inbreeding using TDa95-310.

We find extended runs of homozygosity among our eight sequenced lines, as expected based on their high degree of relatedness (**Fig. 4c**). Long blocks of homozygosity generally stretch across pericentromeric regions, consistent with the low recombination rates in these regions (**Figs. 1** and **4**). Although our sampling is not random, the extensive homozygosity (and identity across genotypes) suggests that there may have been selection for the haplotype on chromosome 20 that appears in a homozygous state in six of our eight breeding lines, as well as some other common haplotypes seen in **Fig. 4d**. The reduced genetic variation present in these breeding lines suggests a strong need for the introduction of additional diversity in yam breeding programs at IITA and other national institutes.

Conversely, we find that multiple genomes contain several long runs of unusually high heterozygosity (**Fig. 4c, Supplementary Fig. 3**). While the typical rate of single nucleotide heterozygosity across 100 kb blocks is ∼7–10 SNVs per kb (excluding runs of homozygosity), these highly heterozygous runs have more than 17.5 SNVs/kb (**Supplementary Fig. 3c**,**d**,**f–g**). In cassava and citrus, blocks of high heterozygosity exceeding 1% variation have been demonstrated to be due to interspecific introgression^52,53^. The co-cultivation of related yam species (**Supplementary Fig. 12, Supplementary Table 7**) by growers and breeders suggests that these blocks (some of which are found overlapping low-recombination-rate pericentromeric regions, e.g., on chromosome 4) are the result of past interspecific introgression. Since the Pacific yam *D. nummularia* is the only other yam species shown to be interfertile with *D. alata*^*20*^, we speculate that it is the source of introgression into greater yam breeding lines, possibly before introduction to Africa. The retention of these hybrid sequences in this germplasm suggests that they may confer some possible adaptive advantage, as has been hypothesized in cassava (*Manihot esculenta* Crantz)^52^. Wolfe et al.^54^ showed that *Manihot glaziovii* Muell. Arg. segments introgressed into and maintained as heterozygous in the cassava genome are associated with preferred traits. In the future, comparison of these highly heterozygous regions with sequences from related *Dioscorea* spp. should reveal the source of these interspecific contributions to the greater yam germplasm.

### V. Concluding Remarks

The near complete and contiguous chromosome-scale assembly of greater yam reported here, along with the associated genetic and genomic resources, opens new avenues for improving this important staple crop. We demonstrated the utility of these resources by finding eight QTL for anthracnose disease resistance and tuber quality traits, which will facilitate marker-assisted breeding in this crop. A major hurdle for breeders is the difficulty of making a successful cross in *D. alata* due to lack of flowering, limited seed set, and differences in flowering time. Genome-enabled methods such as marker-assisted selection, GWAS and genomic selection will allow breeders to make the most out of each cross, and use fewer resources to maintain genotypes that are less likely to be useful. By analyzing the diversity of popular breeding lines, we found that they are highly related and, in some cases, have long runs of homozygosity that reduce the genetic diversity available for selection but may represent genomic regions fixed for desirable traits. Analysis of a broader sampling of African greater yam germplasm will prove valuable to avoiding inbreeding depression associated with inbreeding elite lines^55^. Conversely, we found regions of presumptive interspecific hybridization, pointing to the potential value of broader crosses that may enable the transfer of valuable traits from other yam species while minimizing linkage drag with genome-assisted selection. Similarly, the genome sequence also enables the application of gene editing to directly alter genotypes in a targeted manner, preserving genetic backgrounds that confer cohorts of desirable traits. Although this is the first genome for *D. alata*, its small genome and the advent of rapid long-read technologies open the door to rapidly assemble additional accessions to discover and leverage structural variants for breeding. Such variants have been shown to control important traits such as plant development^56^.

Greater yam has a high potential for increased yield and broader cultivation, with advantages compared with other root-tuber-banana crops due to its superior nutritious content and low glycemic index^57,58^. Greater yam’s ability to grow in tropical and sub-temperate regions around the world suggests that it is highly adaptable to environment, and that there may be adaptive traits (and associated alleles) that could be exploited in different global contexts. It establishes itself vigorously, is higher yielding than other domesticated yam species, and is highly tolerant to marginal, poor soil and drought conditions, and thus likely nutrient use efficient^8^. These traits will be valuable assets in a changing climate. Greater yam is also highly tolerant of the most significant yam virus, yam mosaic virus^19^. By leveraging QTL and genome-wide-association for disease resistance and tuber quality, as well as marker-aided breeding strategies and genome editing, yam breeders are poised to rapidly generate disease resistant, high-performing, farmer-/consumer-preferred, climate-resilient varieties of *D. alata*.

## Methods

### Reference accession

The breeding line TDa95/00328, from the International Institute of Tropical Agriculture (IITA) yam breeding collection, was chosen as the *D. alata* reference genome accession because it is moderately resistant to anthracnose (a fungal disease caused by *Colletotrichum gloeosporioides*, and was confirmed to be diploid by marker segregation analysis)^23,27^. Chromosome number (2n=40) and ploidy were further confirmed through chromosome counting and DNA flow cytometry (**Supplementary Note 1, Supplementary Fig. 2;** Gatarira et al., in preparation).

### Genome sequencing

High molecular weight DNA for Pacific Biosciences (PacBio, Menlo Park, USA) Single-Molecule Real-Time (SMRT) continuous long-read (CLR) sequencing was isolated as described in **Supplementary Note 1**. PacBio library preparation and sequencing were performed at the University of California, Davis Genome and Biomedical Sciences Facility. Three libraries were constructed as per manufacturer protocol, with fragments smaller than 7 kb, 15 kb, and 20 kb, respectively, excluded using Blue Pippin. In total, one RSII and 20 Sequel SMRT cells of CLR data were generated for a combined 235× sequence depth. Half of the 112.4 Gb of generated bases were sequenced in reads 14.5 kb or longer.

For HiC chromatin conformation capture, suspensions of intact nuclei from *D. alata* (TDa95/00328) were prepared from young leaves and apical parts of the stem according to ref.^59^ at the Institute of Experimental Botany, Olomouc, Czech Republic, with modifications as described in **Supplementary Note 1**. These nuclei were sent to Dovetail Genomics for HiC library preparation as described previously^60^. The libraries were sequenced on an Illumina HiSeq 4000 to produce 358.5 million 151 bp paired-end reads.

For genome polishing, a 625 bp insert-size Illumina TruSeq library was made and sequenced on a HiSeq 2500 at UC Berkeley’s Vincent J. Coates Genomics Sequencing Lab (VCGSL), yielding 131 million 251 bp paired reads (137× depth). For contig linking, three Nextera mate-pair libraries (insert sizes ∼2.5 kb, 6 kb, and 9 kb) were prepared and sequenced as 151 bp paired-end reads on a HiSeq 4000 at the UC Davis Genome and Biomedical Sciences Facility. More details are described in **Supplementary Note 1**.

A listing of all TDa95/00328 sequencing data, and corresponding NCBI SRA accession numbers, may be found in **Supplementary Data 1**.

### Genome assembly

We assembled the *D. alata* genome with Canu^61^ v1.7-221-gb5bffcf from the longest 110× of PacBio CLR reads (50.228 Gb in reads 19.8 kb or longer). Contigs were filtered down to a single mosaic haplotype in JuiceBox^62,63^ v1.8.9, taking into account median contig depth (**Supplementary Fig. 13**), sequence similarity, and HiC contacts. Non-redundant contigs were scaffolded into chromosomes using SSPACE^64^ v3 and 3D-DNA^65^ commit 2796c3b. Misassemblies were corrected using genetic maps and JuiceBox HiC visualization. The assembly was polished twice with Arrow^66^ v2.2.2 (SMRT Link v6.0.0.47841) followed by two rounds of Illumina-based polishing with FreeBayes^67^ v1.1.0-54-g49413aa and custom scripts (**Supplementary Note 1)**.

### DArTseq genotyping

DNA was isolated at IITA and NRCRI from their respective mapping populations and parents using modified CTAB methods (**Supplementary Note 2**). DNA samples were genotyped by Integrated Genotyping Service and Support (IGSS, BecA-ILRI hub, Nairobi, Kenya) or DArT (Canberra, Australia) using the ‘high-density’ DArTseq reduced-representation method. DArTseq genotype data may be found in **Supplementary Data 2**. Lists of sequence data used for DArTseq genotyping, and corresponding NCBI SRA accession numbers, are provided in **Supplementary Data 1**.

### Genetic linkage mapping

DArTseq genotyping datasets were mapped to the v2 genome sequence, then filtered for a minimum 90% genotyping completeness and F_1_ Mendelian segregation via χ^2^ goodness-of-fit tests (*p*-value ≥ 1×10^−2^) on allele and genotype frequencies using VCFtools and custom scripts (‘MapTK’ in https://bitbucket.org/rokhsar-lab/gbs-analysis). Half-sibs, off-types, and sample errors were detected as in ref.^68^ and removed. Parental genotypes from one dataset were substituted when a sample by the same name was found to be inconsistent in another. Genotypes were phased and imputed using AlphaFamImpute^69^ v0.1 and parent-averaged linkage maps (**Supplementary Data 3**) constructed in JoinMap^70,71^ v4.1 with the maximum-likelihood mapping function for cross-pollinated populations, which were then integrated into a composite map using LPmerge^72^ v1.7. Further detail regarding genetic linkage mapping can be found in **Supplementary Note 2**.

### RNA sequencing

RNA was extracted at ICRAF from 12 tissues from a single TDa95/00328 plant grown onsite in Nairobi, Kenya. Tissues included leaf petiole, roots, various stages of leaves (initial sprouting leaf, leaf bud, young leaf, semi-matured leaf, matured leaf, fifth leaf), bark, stem, first internode, and middle vine as described in **Supplementary Note 3**. RNA samples were pooled for sequencing by two technologies.

Illumina RNAseq libraries were prepared using the TruSeq stranded mRNA preparation kit (Illumina cat# 20020594) and sequenced at the Agricultural Research Council Biotechnology Platform (ARC-BTP) in Pretoria, South Africa on an Illumina HiSeq 2500 as 125 bp paired ends.

Oxford Nanopore Technologies (ONT) Direct RNA Sequencing (Nanopore DRS) and data processing were performed at the University of Dundee, Dundee, UK. The Nanopore DRS library was prepared using the SQK-RNA001 kit (ONT) as previously described^73^ using 5 µg of total RNA as input for library preparation, and sequenced on R9.4 SpotON Flow Cells (ONT) using a 48 h runtime. Nanopore DRS reads were base-called using Guppy v2.3.1 (ONT), then corrected using proovread v2.14.1 without sampling^74^. Pinfish (ONT; v0.1.0) transcript assembly used corrected reads aligned to the v2 assembly with Minimap2^75^ (v2.8; -x splice). More details on Nanopore transcriptome sequencing are in **Supplementary Note 3**.

### Protein-coding gene annotation

Transcript assemblies (TAs) were constructed with PERTRAN (Shengqiang Shu, unpublished) from 107M pairs of Illumina RNA-seq reads, combining our data with those of ref.^76^ (SRA: SRR1518381 and SRR1518382) and ref.^77^ (SRA: SRR3938623) along with 44k 454 ESTs from ref.^50^ (SRA: SAMN00169815, SAMN00169801, SAMN00169798). A merged set of 86,399 Tas were constructed by PASA^78^ from the above RNA-seq TAs along with 53k assemblies from corrected Nanopore DRS reads, and 18 full-length cDNAs collected from NCBI.

Gene loci were determined by TA alignments and/or EXONERATE^79^ peptide alignments from *Arabidopsis thaliana*^*80*^ TAIR10, *Glycine max*^*81*^ Wm82.a4.v1, *Sorghum bicolor*^*82*^ v3.1.1, *Oryza sativa*^*83*^ v7.0, *Setaria viridis*^*84*^ v2.1, *Amborella trichopoda*^*85*^ v1.0, *Zostera marina*^*86*^ v2.2, *Musa acuminata*^*87*^ v1, *Ananas comosus*^*39*^ v3, and *Vitis vinifera*^*88*^ v2.1 proteomes obtained from Phytozome v13 (https://phytozome-next.jgi.doe.gov) and Swiss-Prot proteins (2018, release 11^89^). Gene models were predicted using FGENESH+^90^, FGENESH_EST, EXONERATE, PASA assembly-derived ORFs, and AUGUSTUS via BRAKER1^91^. After selecting the best-scoring (see **Supplementary Note 3**) predictions at each locus, UTRs and alternative transcripts were added with PASA. The Annotation completeness of this and other Dioscoreaceae species (**Supplementary Table 4**) were measured using BUSCO^30^ v3.0.2-11-g1554283 with the Embryophyta OrthoDB^31^ v10 database.

### Genomic repeat annotation

Repeat annotation was performed twice (see **Supplementary Note 3**) with RepeatMasker^92^. The initial round annotated *de novo* repeats inferred from the preliminary v1 assembly by RepeatModeler^93^ v1.0.11, combined with *Dioscorea* repeats deposited in RepBase^94^. The second round used a repeat library inferred by RepeatModeler v2.0.1 (-LTRstruct) from the more complete v2 assembly.

### Comparisons with other monocot genomes

Orthologous genes were clustered across the available assembled Dioscoreaceae species (*D. alata, D. rotundata, D. dumetorum, D. zingiberensis*, and *T. zeylanicus*) with OrthoFinder^95^ v2.4.1. This procedure produced 5,454 clusters of genes in strict 1:1:1:1 correspondence among the *Dioscorea* species of which 99.9% (n=5451), 90.5% (n=4937), and 99.1% (n=5404) were localized to chromosome-scale scaffolds in *D. alata, D. rotundata*, and *D. zingiberensis*, respectively. We also used OrthoFinder to compare a broader set of monocots (*D. alata, D*.*rotundata, D*.*dumetorum, D*.*zingiberensis, T. zeylanicus, Xerophyta viscosa, Apostasia shenzhenica, Dendrobium catenatum, Asparagus officinalis, Elaeis guineensis, Phoenix dactylifera, Musa acuminata, Oriza sativa, Zea mays, Ananas comosus, Spirodela polyrhiza, Zostera marina, Arabidopsis thaliana*, and *Amborella trichopoda*), which are presented in **Supplementary Fig. 6** using the ClusterVenn^96^ online tool. See **Supplementary Note 3** for more detail.

### Chromosome landscape, Rabl chromatin structure, and centromere estimates

The A/B compartment structure (**Supplementary Fig. 7**) for each chromosome was inferred at 100 kb resolution with Knight-Ruiz (KR)-balanced MapQ30 intra-chromosomal HiC count matrices using a custom script (‘call-compartments’ at https://bitbucket.org/bredeson/artisanal). Centromeric positions were estimated with JuiceBox following the principles described in ref.^97^. Rabl chromatin structure (**Supplementary Note 4**) was visualized in R using ‘prcomp’ (‘chr-structure.R’ script in https://github.com/bredeson/Dioscorea-alata-genomics) on KR-balanced MapQ30 inter-chromosomal HiC count matrices, with chromosome 2 as the reference comparator. Pearson’s correlations (*r*) between gene count, low-complexity and transposable element repeat densities, recombination rate, and A/B compartment domain status were computed using 500 kb non-overlapping windows with BEDtools^98^ v2.28.0 and R^99^ v3.5.3 (**Supplementary Note 4**).

### Synteny and comparative genomics

We used BLASTP^120,121^ v2.10.0 to execute protein searches between *D. alata* and each comparator species. All translated isoforms were included and hits were first filtered for best-scoring alignment per locus pair. Mutually-best hit (MBH) pairs were selected using the C-score ranking method. MBHs within one megabase of the query locus were disallowed in intragenomic comparisons to avoid artifactual alignments between recent, tandemly-duplicated sequences. Coding sequences for each MBH pair were re-aligned as codons with DIALIGN-TX^122^ v1.0.2, unaligned codons were removed, and synonymous substitution (Ks) rates were calculated with the ‘kaks’ function from the SeqinR^123^ v3.6-1 R package. Only MBH pairs with Ks values between 0.0 and 9.0 were retained. All ribbon diagrams were generated with the jcvi.graphics.karyotype module in MCscan^124^ v1.0.14-0-g58b7710b.

MBHs were clustered for conserved collinearity using a minimum spanning tree algorithm (‘cluster-collinear-bedpe’ script in https://bitbucket.org/bredeson/artisanal), requiring a minimum of five MBH pairs per cluster and allowing a maximum extension radius of 20 genes. These ‘lenient’ MBH clusters (provided in **Supplementary Data 4**) were visualized with dotplots and the m:n (e.g., 1:1, 1:2, etc.) patterns of paralogy determined per species comparison. MBHs were then re-clustered to select the best m:n clusters, requiring empirically-determined minimum and maximum Ks thresholds, and a maximum overlap tolerance of 50% between clusters (this was increased to 75% for the *D. alata*-*D. rotundata* comparison to accommodate small internal inversions).

Refined Ks distributions for the ‘delta’ and ‘tau’ duplications were obtained via *D. alata* intragenomic comparison: A maximum ΔKs threshold of 0.15 between edges in a cluster was applied on the ‘constrained’ dataset to obtain the refined delta homoeologs. Tau duplication homoeologs were extracted from the lenient dataset by first subtracting the constrained homoeolog dataset, then filtering for pairs with empirically-determined minimum and maximum Ks thresholds and a ΔKs ≤ 0.20 restriction.

### Phenotyping

#### Planting scheme at IITA

Phenotyping of five mapping populations was performed at IITA from 2016–2019. In 2016, mapping populations were planted in single pots and grown in the screenhouse for seed tuber multiplication and screening of anthracnose disease in a controlled environment. In 2017, individual mini-tubers of each mapping population were pre-planted in pots to ensure germination, and one-month-old seedlings were transplanted in the field using a ridge-and-furrow system. Land preparation, weeding, staking and harvesting were carried out following standard field operating protocol for yam as reported by Asfaw^100^. In 2018 and 2019, harvested tubers were cut into mini-setts of 100 g each, treated with pesticide to prevent rotting, and planted in the field as above. More detail on the planting scheme used at IITA may be found in **Supplementary Note 5**.

### Phenotyping of mapping populations for anthracnose disease

#### Assessment of anthracnose disease at IITA

*Field assessment*: In 2017 and 2018, each plant in the five IITA populations (TDa1401, TDa1402, 1403, 1419 and 1427) was visually scored in the field for yam anthracnose disease (YAD) severity at 3 MAP (months after planting) and 6 MAP using a 1–5 scale as follows: Score 1 = No symptoms, Score 2 = 1–25%, Score 3 = 25–50%, Score 4 = 50–75%, Score 5 = >75%. *Detached leaf assay* (DLA): DLA analysis was performed at IITA in 2016 on plants grown in the screenhouse, and in 2017 and 2018 on plants grown in the field, following a modified protocol of Green et al.^101^ and Nwadili *et al*.^*102*^.

#### Anthracnose disease assessment at NRCRI

At the National Root Crops Research Institute (NRCRI, Umudike, Nigeria), site-specific *C. gloeosporioides* isolates were collected and evaluated, as described in **Supplementary Note 5**. The most virulent isolate was used for anthracnose severity evaluation of NRCRI *D*.*alata* mapping populations using DLA^102^.

More detailed descriptions of phenotyping for anthracnose disease may be found in **Supplementary Note 5**.

### Phenotyping of mapping populations for post-harvest tuber traits

#### Phenotypic characterization of tuber dry matter content at IITA

After harvest, healthy yam tubers were sampled in each replication for dry matter determination. The tubers of each genotype were cleaned with water to remove soil particles. Thereafter, the tubers were peeled and grated for easy oven drying; 100 g of freshly grated tuber flesh sample was weighed, put into a Kraft paper bag, and dried at 105 °C for 16 h. After drying, the weight of each sample was recorded and the dry matter content was determined using the following formula:

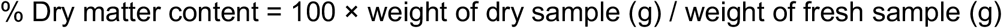

#### Phenotypic characterization of tuber flesh color and oxidation/oxidative browning at IITA

After harvest, one well-developed and mature representative tuber was sampled in each replication. The sampled tuber was peeled, cut, and chipped with a hand chipper to get small thickness size pieces. A chromameter (CR-410, Konica Minolta, Japan) was used to read the total color of sampled pieces placed on a petri dish immediately and exposure to air at 0 min, 30 min, and 180 min. The lightness (L*), red/green coordinate (a*), and yellow/blue coordinate (b*) parameters were recorded for each chromameter reading for the determination of the total color difference. A reference white porcelain tile was used to calibrate the chromameter before each determination^103^.

Tuber whiteness was calculated with the formula (http://docs-hoffmann.de/cielab03022003.pdf):

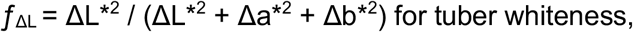

where ΔL* = difference in lightness and darkness ([+] = lighter, [−] = darker, Δa* = difference in red and green ([+] = redder, [−] = greener) and Δb* = difference in yellow and blue ([+] = yellower, [−] = bluer).

Tuber flesh oxidation was estimated from the total variation from the difference in the final and initial color reading as:

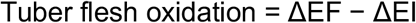

where ΔEF = color reader value at the final time (30 minutes) and ΔEI = Initial color reader value at 0 minutes.

#### Post-harvest tuber evaluation at NRCRI

Of the three populations evaluated at NRCRI, 172 progeny survived. As soon as the yam tubers were harvested, eight traits were assessed using the descriptors from Asfaw^100^: presence or absence of corm (CORM), the ability of corm to separate (CORSEP), type of corm (CORTYP), tuber shape (TBRS), tuber size (TBRSZ), tuber surface texture (TBRST), roots on tuber (RTBS) and position of roots on tuber (PRTBS).

| Trait | Scales |
| --- | --- |
| CORM | 0 = Absent; 1 = Present |
| CORSEP | 0 = No; 1 = Yes |
| CORTYP | 1 = Regular; 2 = Transversally elongated; 3 = Branched |
| TBRS | 1 = Spherical/round; 2 = Oval; 3 = Cylindrical; 5 = Irregular |
| TBRSZ | 1 = Small (less than 15 cm length),<br>2 = Medium (between 15 and 25 cm in length),<br>3 = Big (more than 25 cm in length) |
| TBRST | 1 = Smooth; 2 = Rough |
| RTBS | 0 = No roots; 2 = Few; 3 = Many. |
| PRTBS | 1 = Lower; 2 = Middle; 3 = Upper; 4 = Entire tuber. |

**Supplementary Data 5** contains phenotyping data from each of the mapping populations.

### QTL analysis

QTL association analyses integrated linkage maps, imputed genotype data, and phenotype data into Binary PED files using PLINK^104,105^ v1.90b6.16. Only progeny samples with both genotype and phenotype data were retained per trait. Some traits were initially scored using a discrete 0– 2 system, which PLINK assumes are missing/case/control phenotypes; these traits values were transformed out of the 0–2 range before analysis (e.g., by adding an offset of 1 or 2 to all values, depending on initial data range). An independent QTL association analysis was performed for each trait, and per-locus association *p*-values were adjusted for multiple-testing by max(*T*) correction^104,106^ with 1×10^6^ phenotype label-swap permutations. A locus was considered significant if the max(*T*)-adjusted *p*-value exceeded α= 0.05.

For each identified QTL, an effect plot was generated to determine the dominance pattern and estimate narrow-sense heritability (*h*^2^) at the peak marker. Effect plots and *h*^2^ were calculated as described in ref.^107^ (pg. 122) using a custom R^99^ script. The interval around each QTL peak was determined by extending the region upstream and downstream until LD depreciated to 0.9. The effect status (i.e., dominance) for chr19 and chr6 anthracnose QTL could not be determined because the alleles of interest at these loci are segregating in pseudo-testcross configurations.

### WGS Illumina sequencing

DNA samples from the breeding lines listed in **Supplementary Table 1** were isolated at IITA (**Supplementary Note 6**). TruSeq Illumina libraries were constructed and sequenced at the VCGSL. Inferred insert sizes ranged from 247–876 bp. These libraries were sequenced on HiSeq 2500 or HiSeq 4000 with read lengths ranging from 150–251 bp, yielding combined sample depths of 19 to 230×. **Supplementary Data 1** lists all Illumina sequence data from our breeding lines, including external data, and accompanying summary statistics.

### WGS variant calling

Single-nucleotide variants (SNVs) were called from the whole-genome resequencing datasets listed in **Supplementary Data 1**. Briefly, Illumina reads were screened for TruSeq adapters with fastq-mcf (ea-utils^108^ tool suite) v1.04.807-18-gbd148d4, then aligned with BWA-MEM^109^ v0.7.17-11-g20d0a13 to a TDa95/00328 v2 genome database containing *D. alata* plastid and mitochondrial sequences and a *Pseudomonas synxantha* genome (see **Supplementary Note 6**) as bait for contaminant reads. BAM files were processed with SAMtools^110^ v1.9-93-g0ca96a4 to fix mate information, mark duplicates, sort, merge, and filter for properly-paired reads. Initial SNVs and indels were called with the Genome Analysis ToolKit (GATK; v3.8-1-0-gf15c1c3ef) HaplotypeCaller and GenotypeGVCFs tools^111^. False-positive variant and genotype calls were filtered using sample-specific minimum- and maximum-depth constraints, allele-balance binomial test (two-tailed α= 0.001) thresholds, read depth mask, and annotated repeat masks. See **Supplementary Note 6** for a more complete description. Only biallelic SNVs were used in downstream analyses.

### WGS population analyses

Using 1.89 million SNVs with 75% or more of individuals genotyped, pairwise genome-wide relatedness estimates were obtained with VCFtools^112^ v0.1.16-16-g954e607. The resulting relatedness network and origination year encoded in each sample’s identifier were used to verify IITA pedigrees. Segmental (5000 SNV windows, 1000 SNV step) identity-by-descent (IBD) between sample pairs and the intrinsic heterozygosity or autozygosity of each sample were estimated with custom scripts (‘IBD’ and ‘snvrate’ from https://bitbucket.org/rokhsar-lab/wgs-analysis). A 100 kb sliding window (10 kb step) was called autozygous if the rate of intrinsic heterozygosity was less than 2×10^−4^. This threshold was empirically determined (**Supplementary Fig. 3, Supplementary Note 6**).

### Mitochondrial and plastid sequence assemblies and phylogenetics

Mitochondrial and plastid DNA sequences were assembled using the *de novo* and comparative methods (**Supplementary Note 7**). The IboSweet3 *D. dumetorum* plastid was extracted from the Siajeu et al.^113^ assembly. Our Dioscoreaceae DNA phylogeny was built from plastid long single-copy regions using MAFFT^114,115^ FFT-NS-i v7.427 (--6merpair --maxiterate 1000), Gblocks v0.91b, and PhyML^116^ v3.3.20190909 (--leave_duplicates --freerates -a e -d nt -b 1000 -f m -o tlr -t e -v e). The monocot plastid phylogeny was constructed using OrthoFinder^95,117,118^ v2.4.1 (MAFFT v7.427 alignment and IQ-TREE^119^ v2.0.3 phylogenetic reconstruction). All trees were visualized with FigTree v1.4.4 (https://github.com/rambaut/figtree).

## Supporting information

Supplementary Data 1: Sequence data

Supplementary Data 2: Mapping population DArTseq genotyping reports

Supplementary Data 3a: Genetic linkage maps

Supplementary Data 3b: Genetic linkage maps

Supplementary Data 4: Delta duplication homoeologous segments

Supplementary Data 5a: Phenotyping data YAD field

Supplementary Data 5b: Phenotyping data YAD DLA

Supplementary Data 5c: Phenotyping data tuber traits

## Data availability

The genome sequence, annotation (including both repeat annotations), and SNP data are available on Phytozome (https://phytozome-next.jgi.doe.gov/info/Dalata_v2_1) and YamBase (https://yambase.org/organism/Dioscorea_alata/genome). The genome assembly and sequence data generated for this work are deposited at NCBI under BioProject PRJNA666450.

## Code availability

Custom scripts used for analyzing resequencing, HiC, linkage mapping, and QTL data are available at [https://github.com/bredeson/Dioscorea-alata-genomics].

## Acknowledgements

We thank Oanh Nguyen for troubleshooting and advice for DNA isolation and PacBio sequencing, Emily Kumimoto for mate pair libraries, and Lutz Froenicke for management, at the University of California, Davis Genome and Biomedical Sciences Facility, Davis, CA. Andrzej Kilian (Diversity Arrays Technology); Clay Sneller, Jackline Chepkoech, Mercy Chepngetich, and IGSS/SEQART staff at BecA-ILRI Hub, for facilitating DArTseq genotyping. We thank the staff of Bioscience Center, Yam Breeding Unit, Pathology/Virology Unit, and Farm Office at IITA, Ibadan, Nigeria for all their support in laboratory and field activities. Special thanks to Kwabena Darkwa and Agre Paterne, IITA, Ibadan Nigeria for their support in phenotyping population TDa1401. Boas Pucker provided the single-haploid assembly of *D. dumetorum*. Christopher Saski and Mary Duke provided WGS data of TDa95/00328 and TDa95-310. We thank Ismail Rabbi for early discussions in proposal development, and he and Gezahegn Girma for providing *D. alata* DNA of specific breeding lines.

This work is based on a project supported by the National Science Foundation BREAD program, Award No. 1543967 to DSR, RB, and JEO.

We wish to acknowledge subsidy from the Integrated Genotyping Service and Support platform, a collaborative project between the International Livestock Research Institute (ILRI) and the Bill and Melinda Gates Foundation.

DNA extractions for PacBio sequencing, and RNA extractions, were carried out at ICRAF with partial support from the African Orphan Crops Consortium.

RNAseq was funded by the Illumina Greater Good Initiative.

Nanopore DRS work was supported by The University of Dundee Global Challenges Research Fund to GGS & GJB, Biotechnology and Biological Sciences Research Council (BB/M004155/1) to GGS & GJB, and H2020 Marie Skłodowska-Curie Actions (799300) to KK.

Sequencing performed at the Vincent J. Coates Genomics Sequencing Laboratory, UC Berkeley, was partially supported by NIH S10 OD018174 Instrumentation Grant.

DSR was supported by Chan Zuckerberg BioHub, internal funds at the Okinawa Institute of Science and Technology, and the Marthella Foskett-Brown Chair in Biological Science at UC Berkeley.

## Author Contributions

Conceived, designed, and led study: DSR, RB, JEO, JVB, JBL

Genome assembly and chromatin structure, chromosome landscape, comparative genomics, chromosome evolution, population genetic, and phylogenetic analyses: JVB (lead), DSR

Genome sequencing planning and coordination: DSR, JVB, AVD, JBL Genetic mapping: JVB (lead), JBL

QTL analysis: JVB

Overall project management: JBL

Mapping population development: ALM, AA

Mapping population management/propagation: RB, IOO, AA, JN, IN Development of and info on breeding lines: RA, ALM

Phenotyping of mapping populations: IOO, OK, AA, PLK, NRO, CON, IN, JN

Preparation of cell nuclei for HiC analysis; karyotype and chromosome counting: JD (lead), EH

Nanopore DRS sequencing and analysis: MP, KK, AVS, GJB, GGS (lead)

DNA isolation for reference genome, sequencing of breeding lines, and genotyping: IOO, NRO, JN, RK, SM, PSH

RNA isolation: RK, SM, PSH (lead) Provision of RNAseq data: JF

Wrote manuscript: DSR, JVB, JBL, JEO, OK, NRO, CN, RB, EH with input from AVD, GGS, JD

Annotation and database management: DG (lead), SS, JC

Other project planning/site-specific supervision: IOO, CNE, RJ, AM

## Ethics Declarations

### Competing Interests

DSR is a member of the Scientific Advisory Board of, and a minor shareholder in, Dovetail Genomics LLC, which provides as a service the high-throughput chromatin conformation capture (Hi-C) technology used in this study.

## Supplementary Information

(Provided in a separate document)

Supplementary Fig. 1: Genome-wide HiC contact matrix of TDa95/00328 chromosomes.

Supplementary Fig. 2: TDa95/00328 chromosome count.

Supplementary Fig. 3: Histograms summarizing the rates of haplotypic heterozygosity and homozygosity calculated in 100 kb sliding windows, with 10 kb step.

Supplementary Fig. 4: Principal Component Analysis (PCA) for Rabl conformation.

Supplementary Fig. 5: Chromosome-scale assembly and composite linkage map comparison.

Supplementary Fig. 6: Protein-coding gene orthology table.

Supplementary Fig. 7: A/B compartment structure.

Supplementary Fig. 8: Evidence of conserved synteny and paleotetraploidy.

Supplementary Fig. 9: Phylogenetic distribution of monocot plastid genome sequences.

Supplementary Fig. 10: Anthracnose quantitative trait locus analyses.

Supplementary Fig. 11: Tuber trait QTL scans.

Supplementary Fig. 12: Phylogenetic distribution of Dioscoreaceae plastid genome sequences.

Supplementary Fig. 13: Evaluating assembled contig ploidy.

Supplementary Fig. 14: Minimum and maximum allele-balance filters.

Supplementary Table 1: Traits of parents of mapping populations.

Supplementary Table 2: Component and composite linkage maps.

Supplementary Table 3: *D. alata* repeat class count and nucleotide abundances.

Supplementary Table 4: Dioscoreaceae assembly and annotation BUSCO comparisons.

Supplementary Table 5: Pairwise synonymous substitution rate and identity matrix.

Supplementary Table 6: Relatedness among eight sequenced breeding lines.

Supplementary Table 7: Species and accessions used in this work.

Supplementary Table 8: Parameters used for mapping in JoinMap.

Supplementary Note 1: Chromosome counting, genome sequencing, and assembly

Supplementary Note 2: Genetic linkage mapping

Supplementary Note 3: RNA sequencing and genome annotation

Supplementary Note 4: Chromosome landscape, Rabl chromatin structure, and centromere estimation

Supplementary Note 5: Phenotyping

Supplementary Note 6: Whole-genome ancestry reconstruction and population genetic analysis

Supplementary Note 7: Mitochondrial and plastid sequence assemblies

## Supplementary Data Files

**Supplementary Data 1: Sequence data. (Excel file)**

The first tab of this file lists WGS data generated for this work, associated sequencing statistics, and SRA deposition numbers. Data used from external sources for *D. alata* breeding lines TDa95-310^51^ and TDa95/00328^51^ (TGAC); and *D. dumetorum*^*113*^, are also listed in the first tab. Subsequent tabs list parents and progeny from mapping populations, including GenBank biosample and SRA deposition numbers.

**Supplementary Data 2: Mapping population DArTseq genotyping reports. (Excel file)**

DArTseq genotype reports for individual and combined populations. All reports are in “single-row” format, with the exception of TDa1402, which is in “two-row” format.

**Supplementary Data 3a–b: Genetic linkage maps. (two Excel files)**

Files of genetic linkage maps for the individual and combined populations. Supplementary Data 3a consolidates maps containing all markers and includes the composite linkage map. Supplementary Data 3b consolidates the linkage maps with non-redundant marker sets used for the QTL analyses.

**Supplementary Data 4: Delta duplication homoeologous segments. (Excel file)**

File listing the homoeologous collinear segments between chromosomes arising from the delta duplication.

**Supplementary Data 5a–c: Phenotyping data. (three Excel files)**

Phenotype data for mapping populations. One file for each of yam anthracnose disease (YAD) severity field assay, YAD detached leaf assay (DLA), and tuber traits, respectively.

## Notes

### Competing Interest Statement

Author Daniel Rokhsar is a member of the Scientific Advisory Board of, and a minor shareholder in, Dovetail Genomics LLC, which provides as a service the high-throughput chromatin conformation capture (Hi-C) technology used in this study.

