## Supplementary Information for "Chromosome evolution and the genetic basis of agronomically important traits in greater yam"

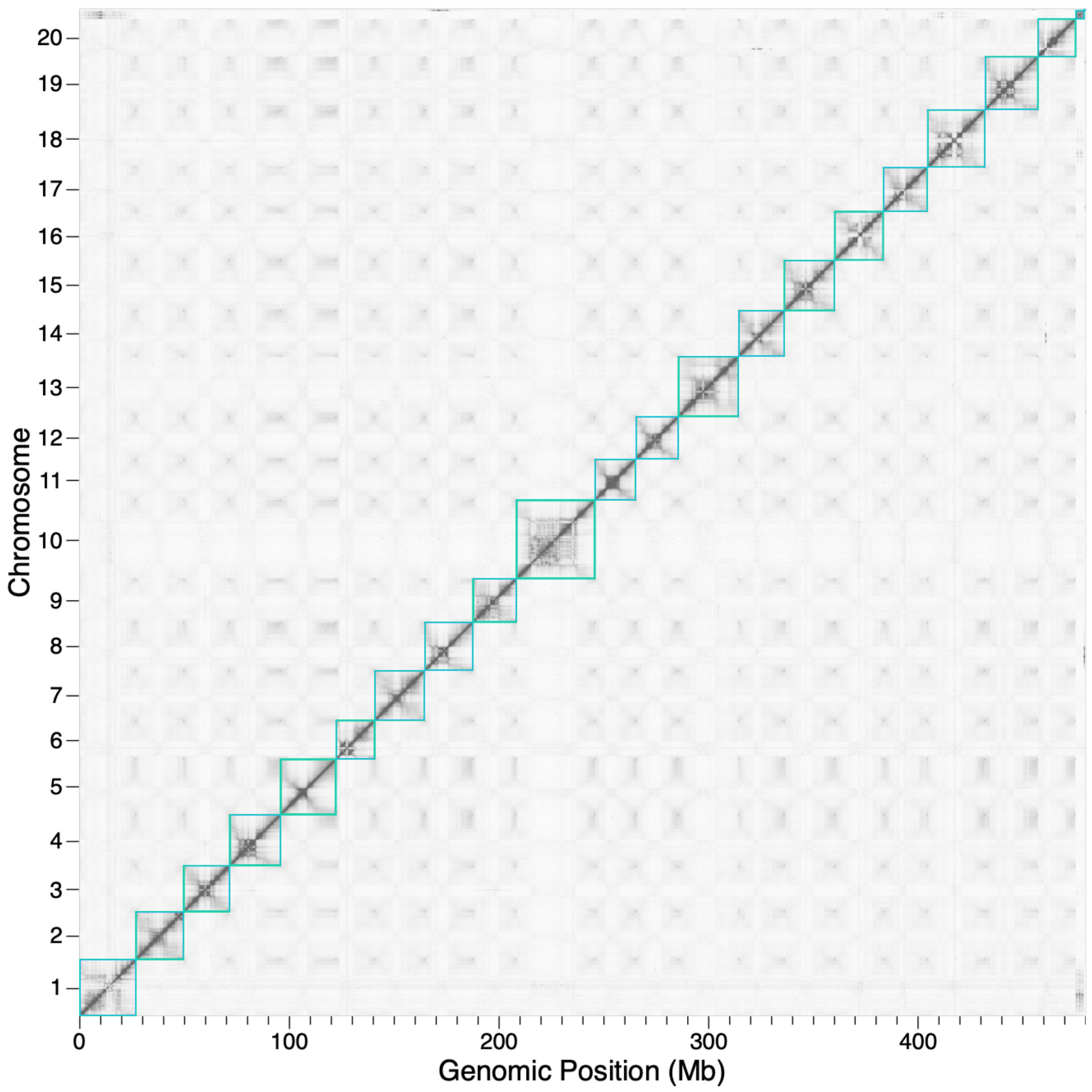


**Supplementary Fig. 1 Genome-wide HiC contact matrix of TDa95/00328 chromosomes.** Within chromosomes, the high contact density along the diagonal reflects the well-ordered underlying assembly. The apparent 'winged' patterns within chromosomes, and the elevated contact densities between chromosome ends, is typical of Rabl-structured chromosomes in the nucleus. Chromosomes are outlined with cyan boxes. Each pixel represents the intersection between a pair of 250 kb loci along the chromosomes. The density of contacts between two loci is proportional to pixel color, with darker pixels representing more contacts and lighter representing fewer. Chromosome numbers are listed along the Y-axis, and cumulative genomic position (in Mb) along the X-axis.

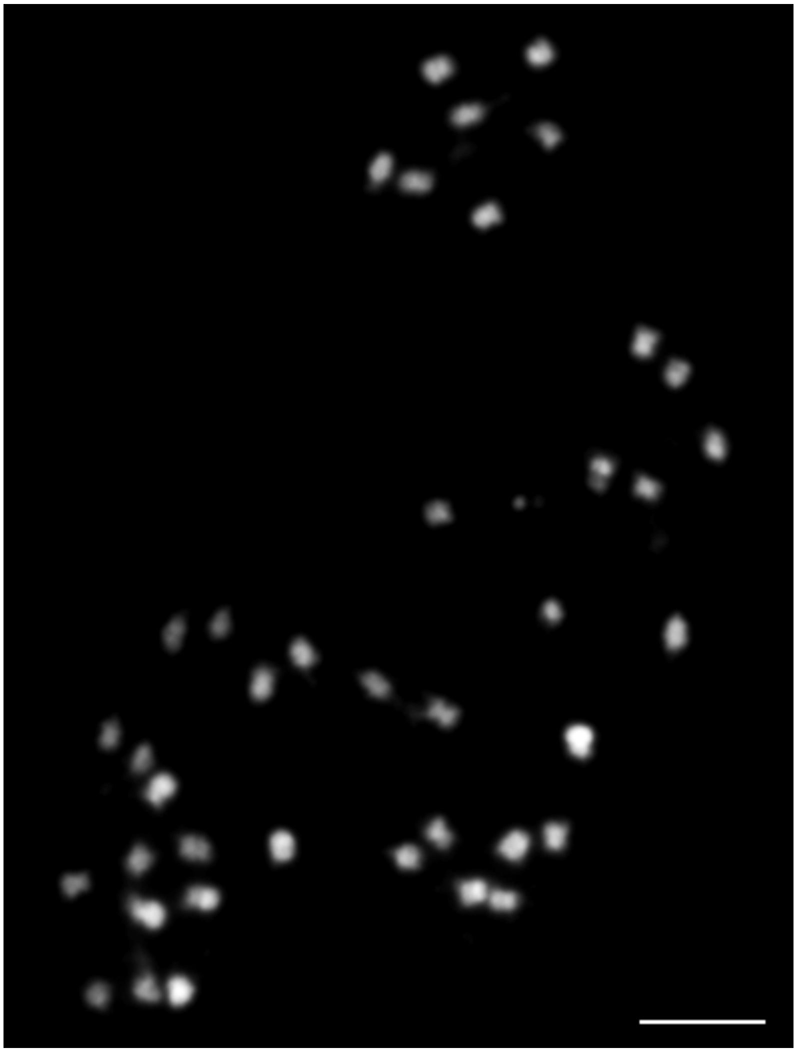

**Supplementary Fig. 2 TDa95/00328 chromosome count.** Mitotic metaphase chromosomes of *D. alata* TDa95/00328 (2n=40). Scale bar corresponds to 5 μm.

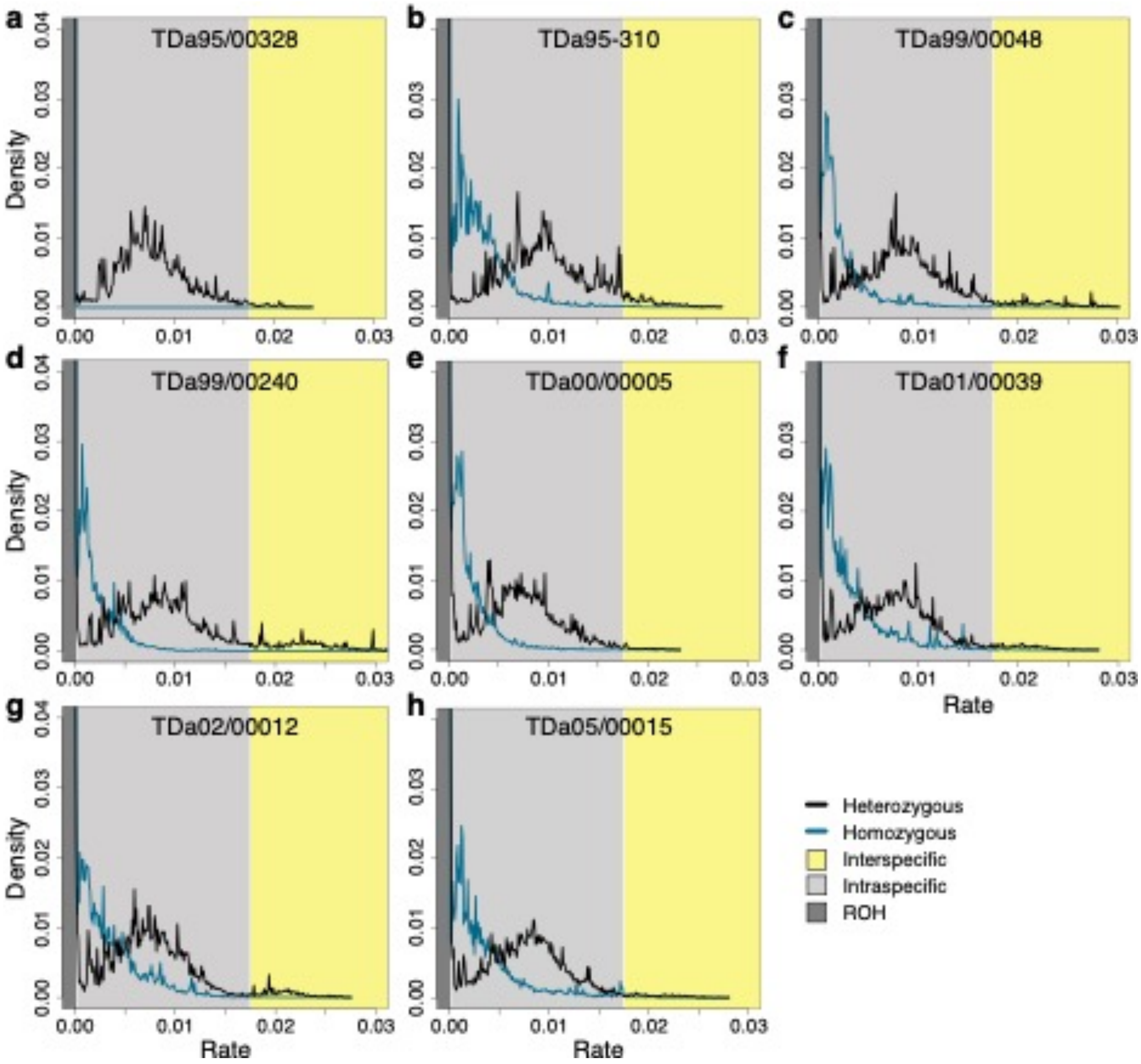


**Supplementary Fig. 3 Histograms summarizing the rates of haplotypic heterozygosity and homozygosity calculated in 100 kb sliding windows, with 10 kb step.** Broadly, three heterozygosity regimes were defined: Runs of homozygosity (ROH, black field), requiring a near-zero (<0.0002) rate of heterozygous SNVs; intraspecific variation (heterozygosity < 0.0175, grey field); and possible interspecific variation (heterozygosity ≥ 0.0175, yellow field). Histograms for each of the eight breeding lines are shown: **a** TDa95/00328, **b** TDa95-310, **c** TDa99/00048, **d** TDa99/00240, **e** TDa00/00005, **f** TDa01/00039, **g** TDa02/00012, and **h** TDa05/00015.

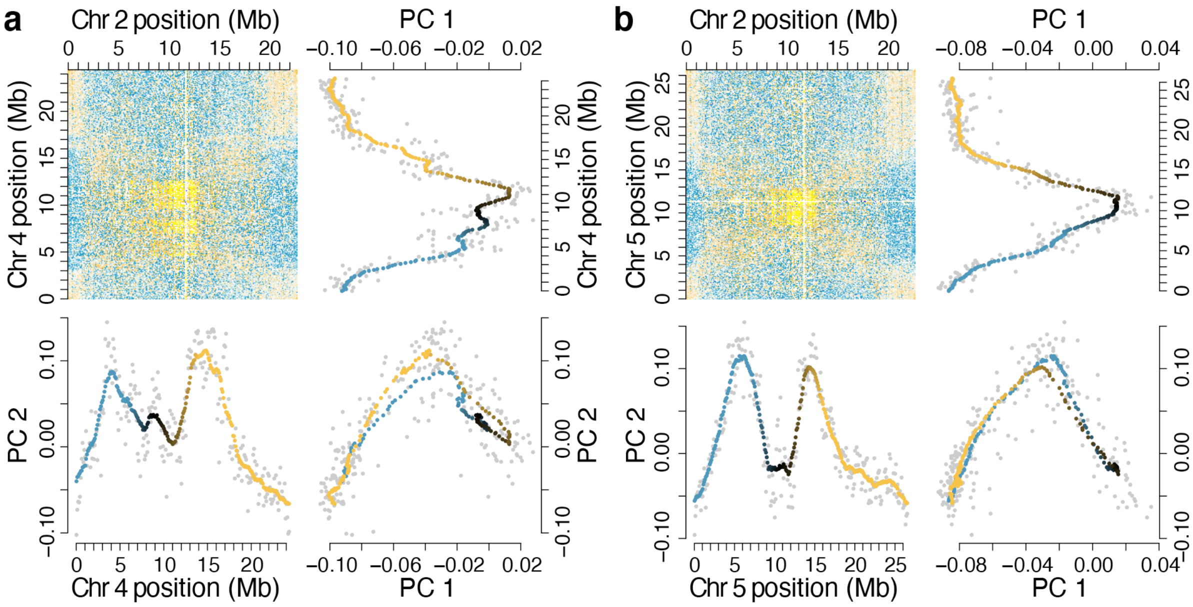


**Supplementary Fig. 4 Principal Component Analysis (PCA) for Rabl conformation.** The first principal component of variation (PC1) traverses the centromere-telomere axis, while the second (PC2) traverses the arm-arm axis (i.e., enriched contacts between positions of chromosome arms). PCA of chromosome 2 contacts with **a** chromosome 4 and **b** chromosome 5. For each panel, the HiC contact enrichment matrix (yellow, enrichment; cyan, depletion; white, parity or no data) is plotted in the upper-left quadrant, PC1 loadings plotted against chromosome position in the upper-right quadrant, PC2 loadings plotted along the chromosome in the lower-left quadrant, and PC1 plotted against PC2 in the lower-right. The smoothed loadings for each PC are colored along a gradient from cyan, to black, to gold for the p-arm, centromere, and q-arm, respectively. The original PC loadings are plotted as grey points.

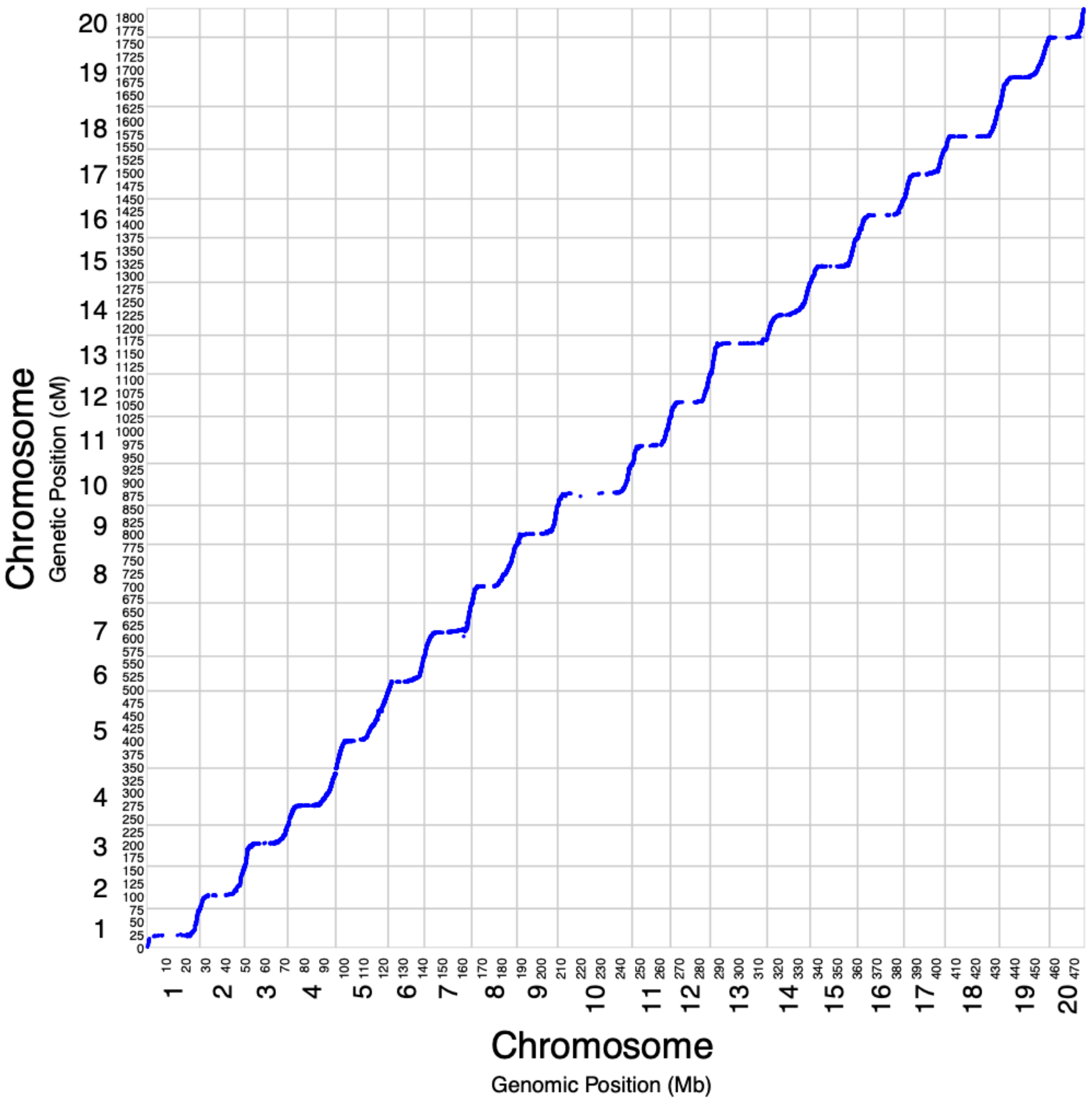


**Supplementary Fig. 5 Chromosome-scale assembly and composite linkage map comparison.** Dot plot demonstrating the large-scale agreement between the 20 assembled chromosome-scale scaffolds and the high-resolution composite linkage map. Each blue point represents one of the 10,448 markers in the composite map plotted by its genomic position (X-axis) and genetic position (Y-axis). The relatively smooth sigmoidal lines formed by the points and general lack of point scatter illustrate the high-degree of structural correctness down to ~8.3 kb resolution. Chromosome numbers (large text) and cumulative coordinates (small text) are listed on both axes.

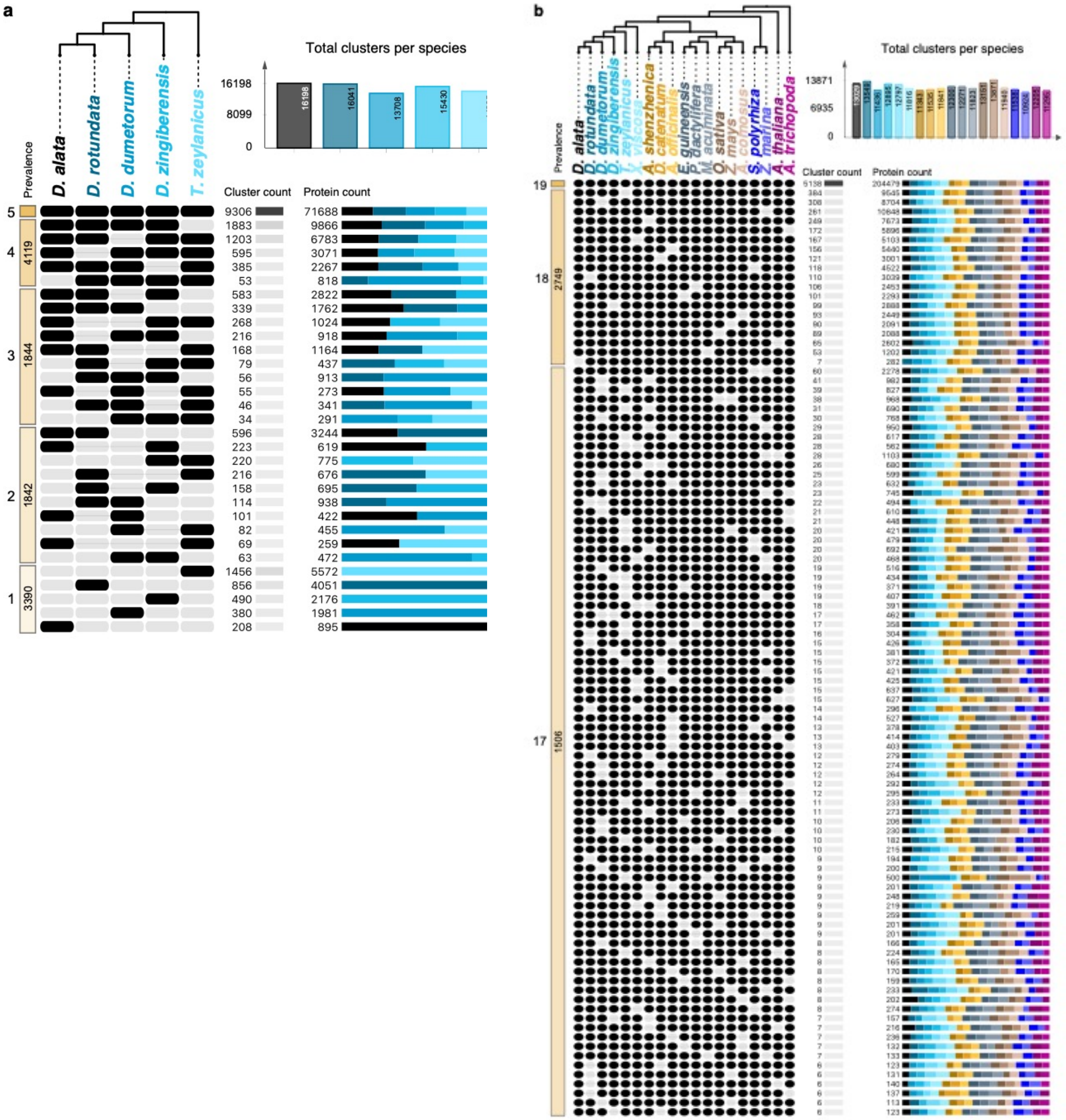


**Supplementary Fig. 6 Protein-coding gene orthology table. a** For the Dioscoreaceae genomes shown, 19,934 total orthologous clusters were inferred with OrthoFinder and visualized with ClusterVenn. Species-level ortholog membership presence (black ovals) or absence (light grey ovals) is plotted, sorted by cluster prevalence and count in descending order. Cluster and protein counts are featured to the right. For example, the second row indicates there are 2,426 orthologous clusters (composed of 9,877 total genes) present in all Dioscorea species but absent in *Trichopus zeylanicus*. Horizontal bar plots illustrate the relative frequency of genes by species per cluster prevalence category. Bars are colored to match their species labels. Also shown are a histogram of total clusters found in each species. **b** Orthology membership is summarized for thirteen monocot species and two outgroups, Arabidopsis thaliana and *Amborella trichopoda*, as in panel a. A cladogram of species relationships is also provided above. In total, 456,577 proteins were grouped into 25,425 orthologous gene clusters.

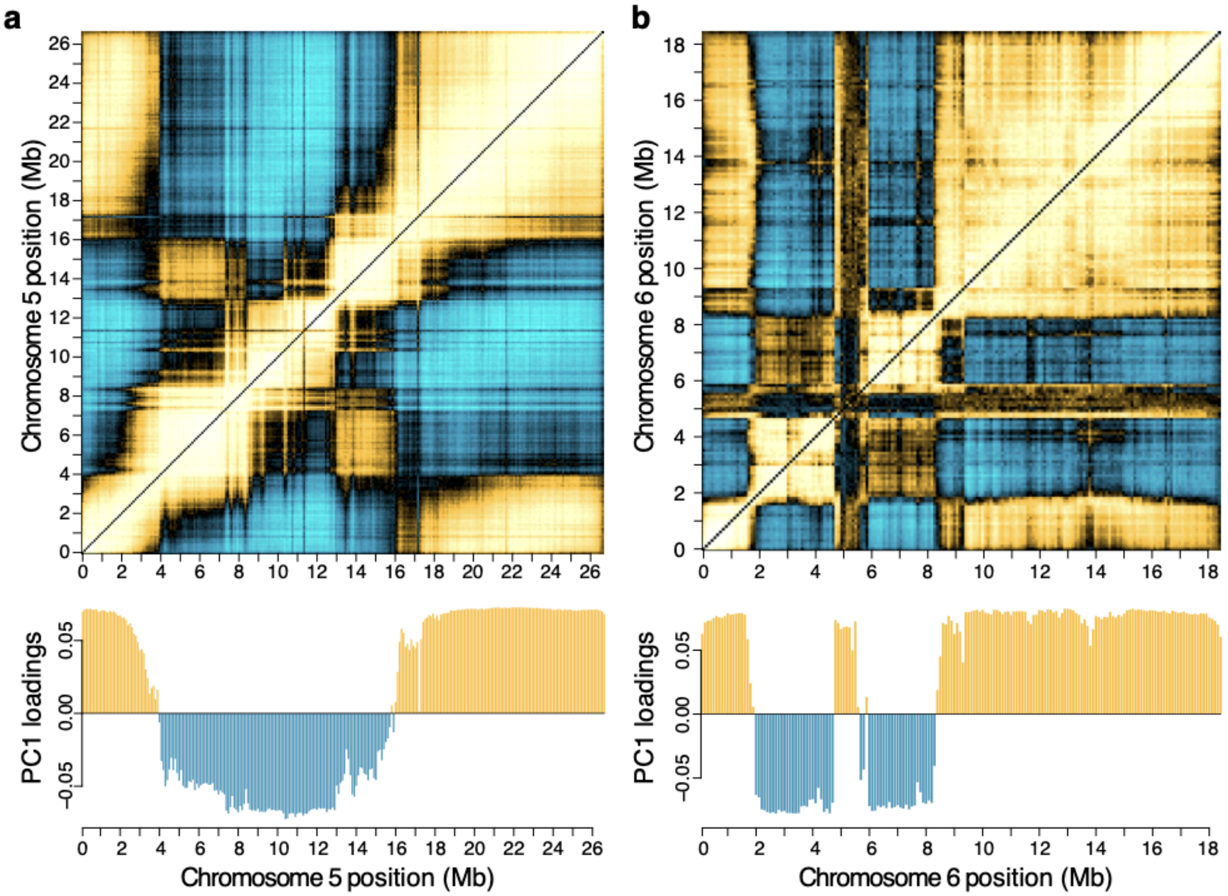


**Supplementary Fig. 7 A/B compartment structure.** Intra-chromosomal HiC contact correlation matrices and the A/B compartment structure extracted from each. Colored pixels in the matrices reflect Pearson's correlation coefficient values between a pair of 100 kb non-overlapping loci, where positively-correlated loci are represented in yellow, negatively-correlated loci in cyan, and weakly-correlated/uncorrelated loci in black. The A/B compartment structure for each chromosome was inferred from the first principal component (PC1) loadings plotted against chromosome position. To illustrate, **a** chromosome 5 and **b** chromosome 6 are shown.

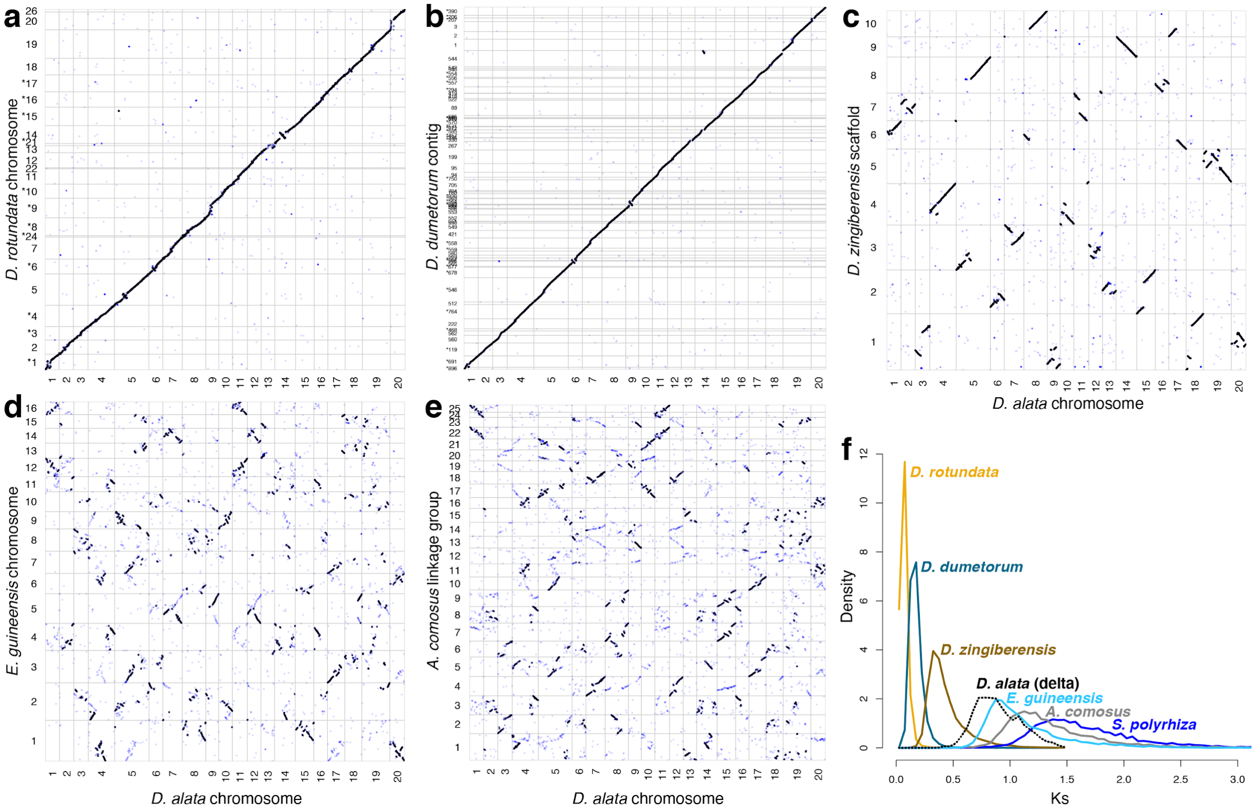


**Supplementary Fig. 8 Evidence of conserved synteny and paleotetraploidy.** Conserved synteny between *Dioscorea alata* and **a** *D. rotundata,* **b** *D. dumetorum*, **c** *D. zingiberensis*, **d** *Elaeis guineensis*, and **e** *Ananas comosus*. Points represent X-Y pairs of genes plotted by their respective genome-wide ordinal indices. Dense linear clusters of genes correspond to segments of chromosomes with shared ancestry. Boundaries between chromosomes are represented by horizontal and vertical grey lines. For clarity, only chromosome-scale scaffolds and contigs larger than 1 Mb are shown. Sequences reverse complemented for plotting are marked with asterisks. Rates of synonymous substitution (Ks) calculated between homoeologous *D. alata* (n=1,578) gene pairs and between *D. alata* orthologs in *D. rotundata* (n=14,889), *D. dumetorum* (n=13,667), *E.guineensis* (n=7,296), and *A. comosus* (n=6,405).

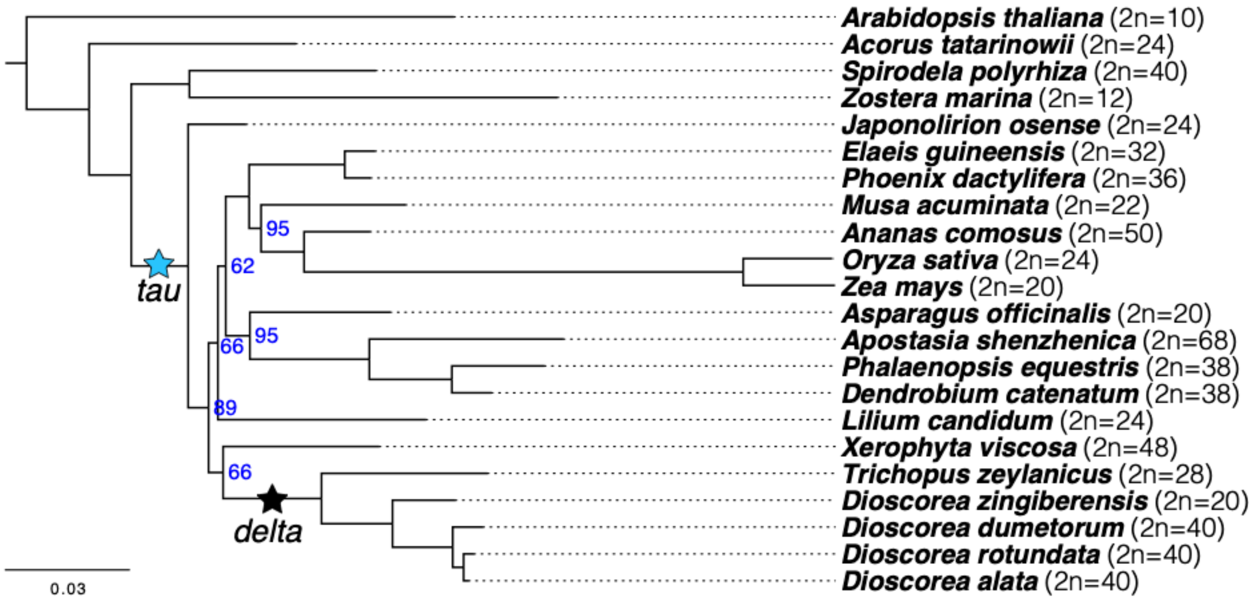


**Supplementary Fig. 9 Phylogenetic distribution of monocot plastid genome sequences.** Plastid protein phylogeny of Liliopsid species with *Arabidopsis thaliana* as an outgroup and rooted by *Amborella trichopoda* (not shown). Selected whole-genome duplication events are depicted with stars. Names for each species are shown in bold text and their diploid chromosome number annotated in parentheses. Bootstrap support (blue) is shown in percentages for nodes with values less than 100% (n=1000).

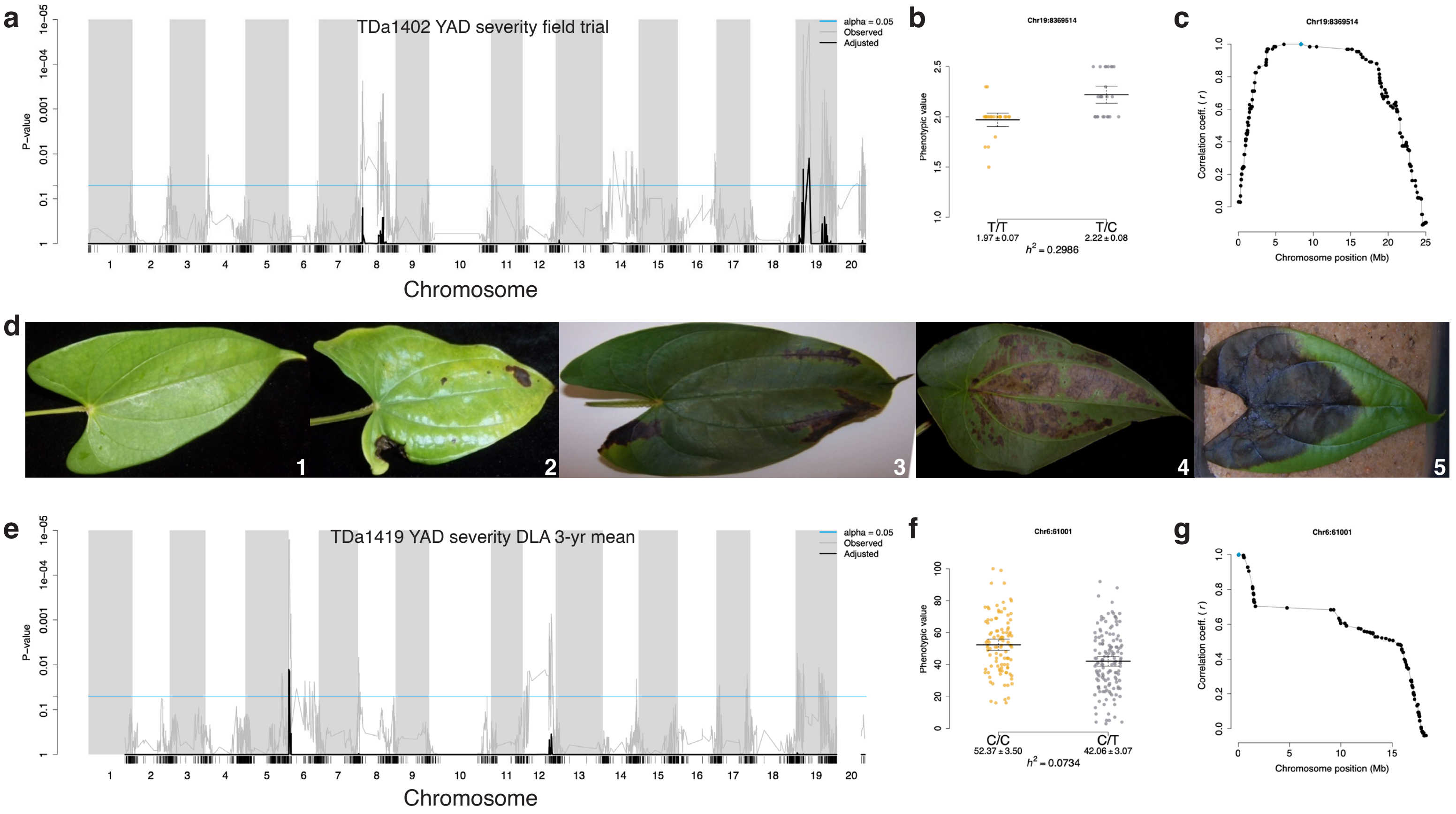


**Supplementary Fig. 10 Anthracnose quantitative trait locus analyses.** Association analyses for yam anthracnose disease (YAD) resistance were conducted for some crosses with both field trial and detached-leaf assay (DLA) susceptibility phenotype data. Plots **a**–**c** show QTL association mapping results for the TDa1402 population YAD field trial for year 2018. **a** Genome-wide QTL association scan. The family-wise max($T$)-corrected association significance values for each genotyped locus is represented with black lines, and the uncorrected significance in grey lines. The minimum threshold for significance (*alpha* = 0.05) is represented as a cyan horizontal line. **b** An effect plot for the peak locus on chromosome 19 (at 8.4 Mb) with the estimated phenotypic variance (i.e., narrow-sense heritability, $h^2$). The alleles and genotypes are shown along the X-axis with their phenotypic values (mean ± 95% confidence interval). **c** Plot showing the strength of linkage disequilibrium (LD) between the peak marker (cyan diamond) and other loci (black points) in chromosome 19. LD was calculated as Pearson's correlation ($r$). **d** Example images showing the DLA YAD severity scale (1–5), calculated from the relative infected leaf area. Plots **e**–**g** show significant QTL association results for the putative QTL identified on chromosome 6 at 61 kb with the TDa1419 population YAD DLA three-year (2016–2018) mean data.

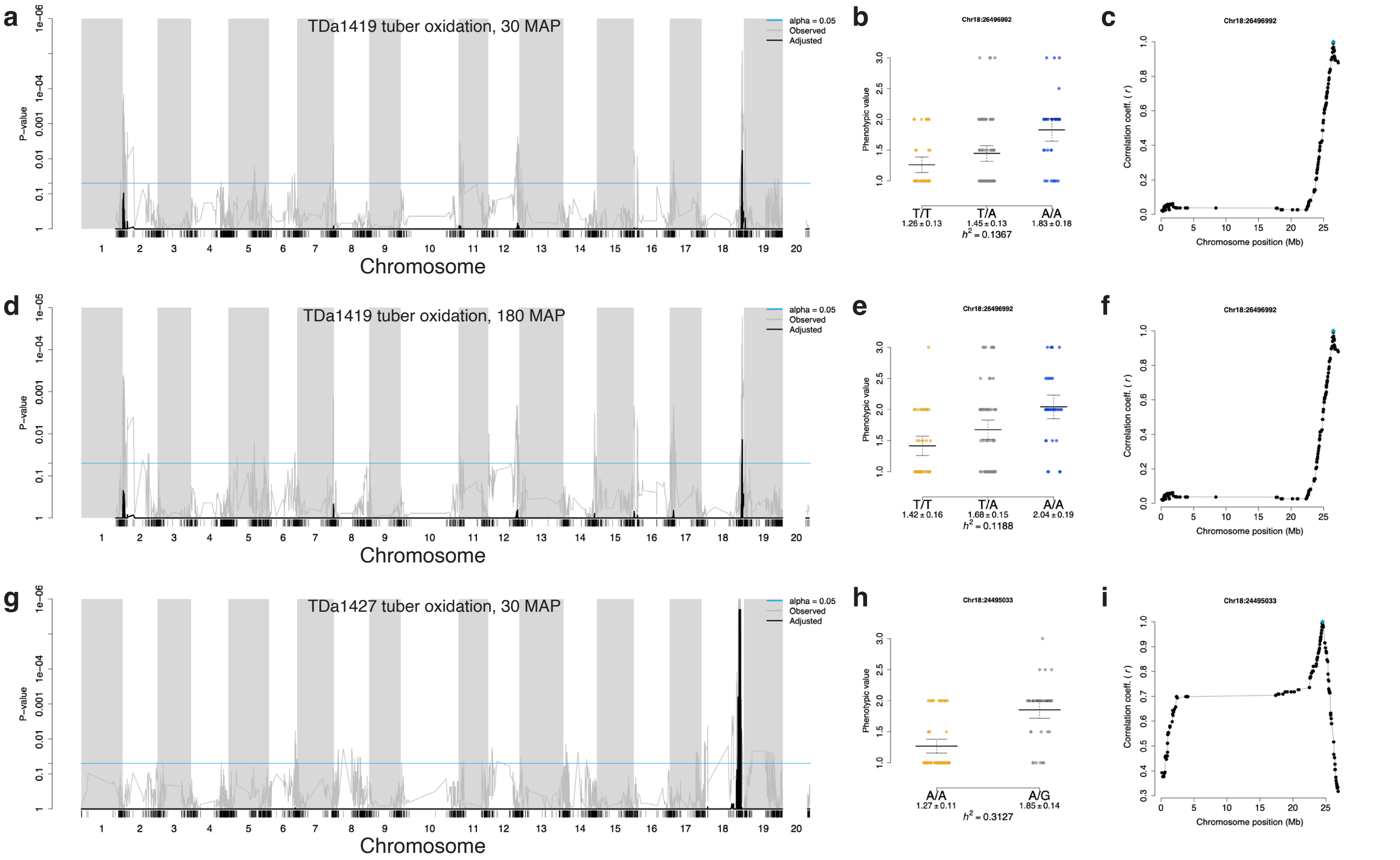
a
TDa1419 tuber oxidation, 30 MAP
alpha = 0.05
Observed
Adjusted
P−value
Chromosome
b
Chr18:26496992
Phenotypic value
T/T
1.26 ± 0.13
T/A
1.45 ± 0.13
A/A
1.83 ± 0.18
$h^2$ = 0.1367
c
Chr18:26496992
Correlation coeff. ( r )
Chromosome position (Mb)
d
TDa1419 tuber oxidation, 180 MAP
alpha = 0.05
Observed
Adjusted
P−value
Chromosome
e
Chr18:26496992
Phenotypic value
T/T
1.42 ± 0.16
T/A
1.68 ± 0.15
A/A
2.04 ± 0.19
$h^2$ = 0.1188
f
Chr18:26496992
Correlation coeff. ( r )
Chromosome position (Mb)
g
TDa1427 tuber oxidation, 30 MAP
alpha = 0.05
Observed
Adjusted
P−value
Chromosome
h
Chr18:24495033
Phenotypic value
A/A
1.27 ± 0.11
A/G
1.85 ± 0.14
$h^2$ = 0.3127
i
Chr18:24495033
Correlation coeff. ( r )
Chromosome position (Mb)

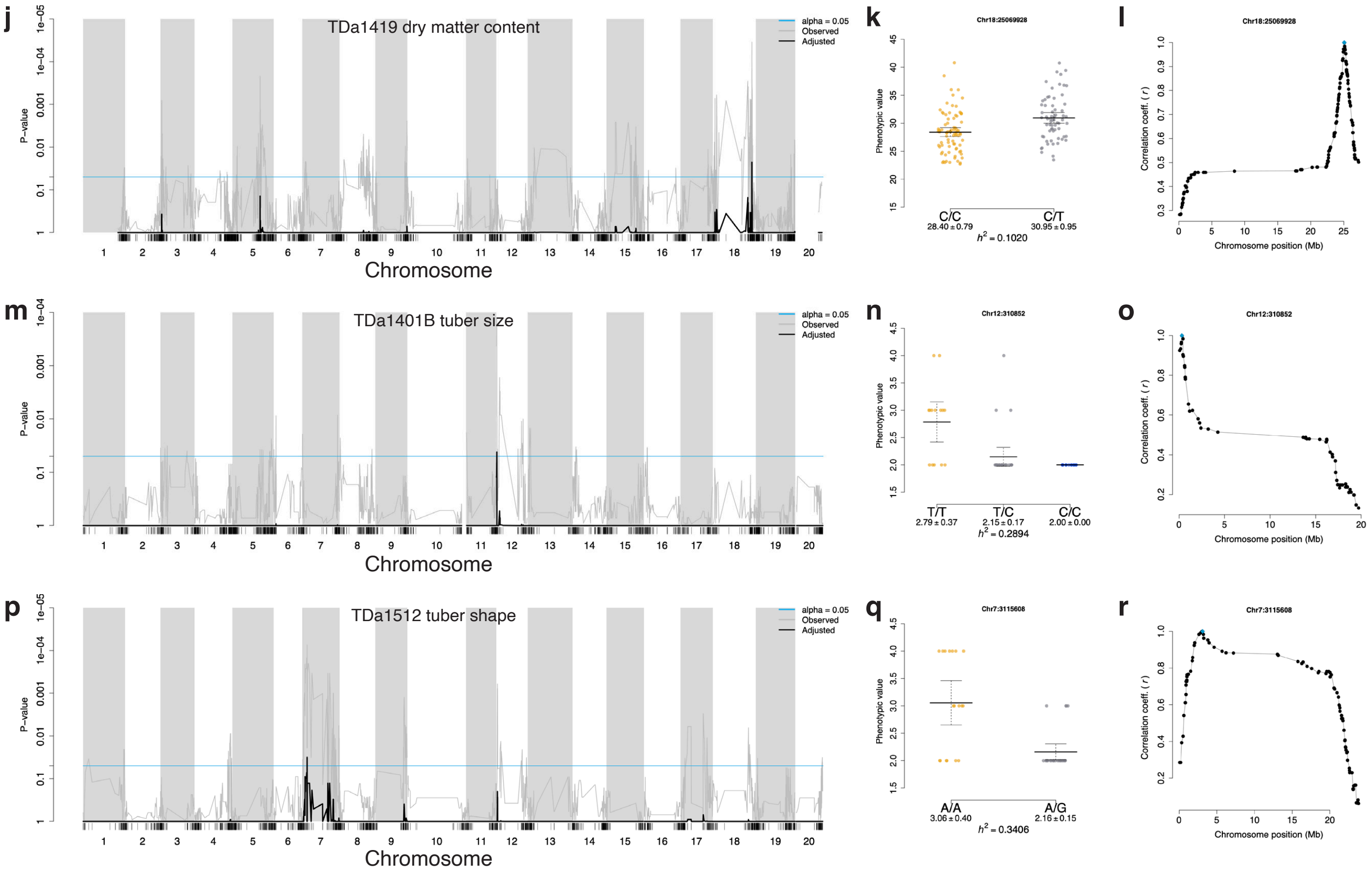


**Supplementary Fig. 11 Tuber trait QTL scans. a** Genome-wide QTL association scan. The family-wise max(*T*)-corrected association significance values for each genotyped locus is represented with black lines, and the uncorrected significance in grey lines. The minimum threshold for significance (*alpha* = 0.05) is represented as a cyan horizontal line. **b** An effect plot for the peak locus with the estimated phenotypic variance (i.e., narrow-sense heritability, $h^2$). The alleles and genotypes are shown along the X-axis with their phenotypic values (mean ± 95% confidence interval). **c** Plot showing the strength of linkage disequilibrium (LD) between the peak marker (cyan diamond) and other loci (black points) in the chromosome. LD was calculated as Pearson's correlation (*r*). **a–c** TDa1419, tuber oxidation 30 mins after peeling (MAP); **d–f** TDa1419, tuber oxidation 180 MAP; **g–i** TDa1427, tuber oxidation 30 MAP; **j–l** TDa1419, dry matter content; **m–o** TDa1401B population, tuber size; and **p–r** TDa1512 population, tuber shape.

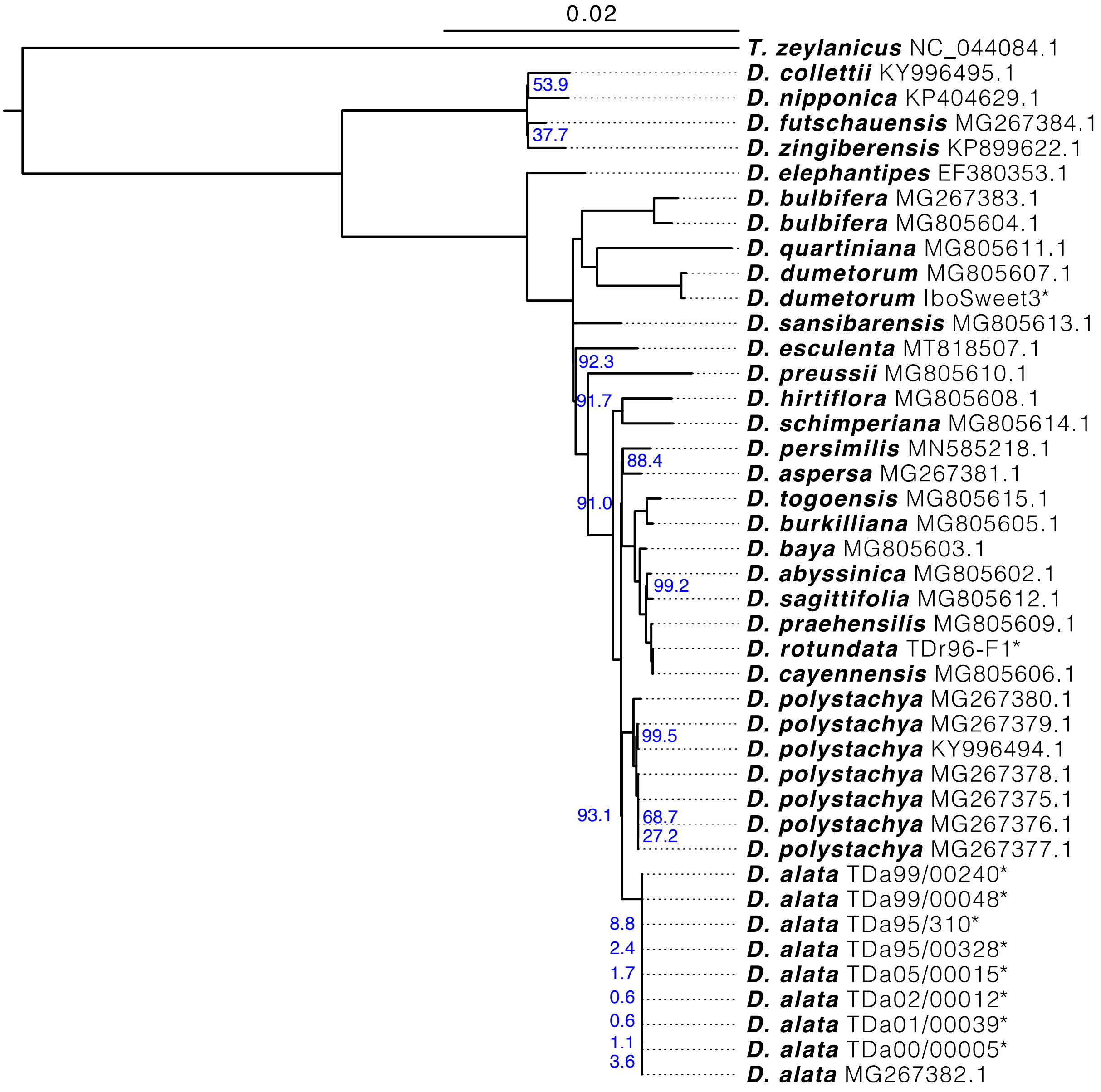


**Supplementary Fig. 12 Phylogenetic distribution of Dioscoreaceae plastid genome sequences.** Maximum likelihood phylogeny of 42 *Dioscorea* plastid genome sequences (26 distinct species) and a related *Trichopus zeylanicus*. Species names for each sample are shown in bold, and sample name or NCBI accession number in light text. Sequences assembled in this study have been asterisked. The phylogeny was rooted with *Xerophyta viscosa* (not shown) and 1000 bootstrap support iterations were performed. Bootstrap support (blue) is shown in percentages for nodes with values less than 100%. IboSweet3 and TDr96-F1 were assembled using sequence data from Siadjeu et al.[1] and Tamiru et al.[2], respectively.

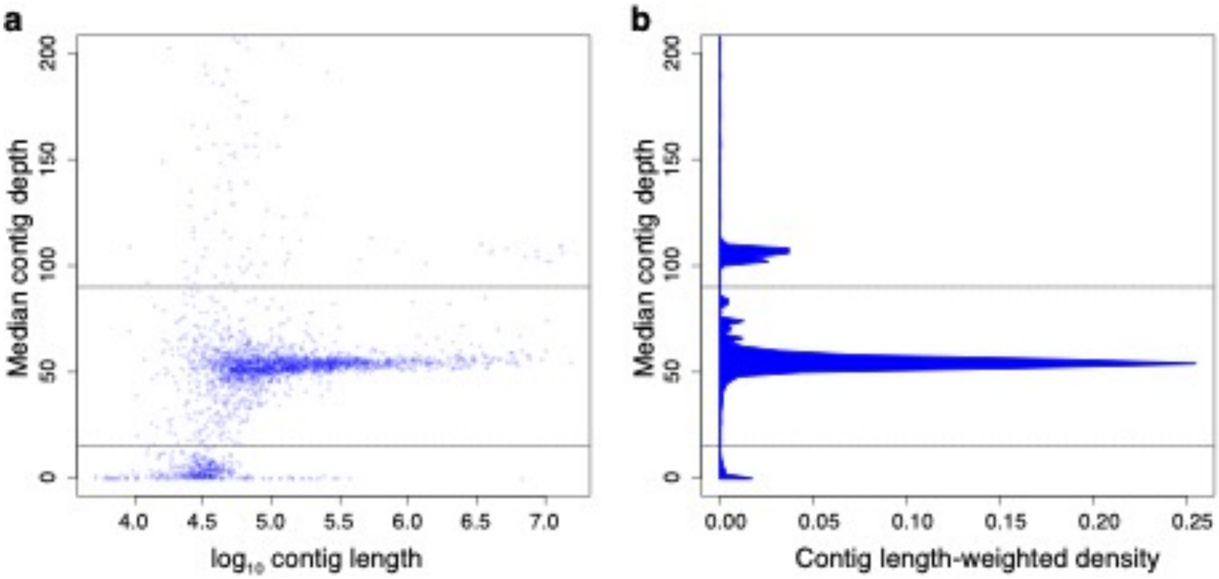


**Supplementary Fig. 13 Evaluating assembled contig ploidy. a** Median depth (Y-axis) for each contig (blue points) is plotted against the $\log_{10}$ contig length (X-axis), showing three depth strata (demarcated by horizontal dotted lines). **b** Median contig depth (Y-axis) is plotted against the density of sequences weighted by their nucleotide lengths (X-axis), illustrating that the preponderance of assembled genome was resolved into single-haplotype contigs, rather than collapsed. Putatively artifactual sequences are captured in the lower stratum, contigs at 1× haploid copies (or 53× depth of coverage) in the middle stratum, and collapsed diploid and repeat contigs, at 2× or greater haploid copies (106× depth), in the upper stratum.

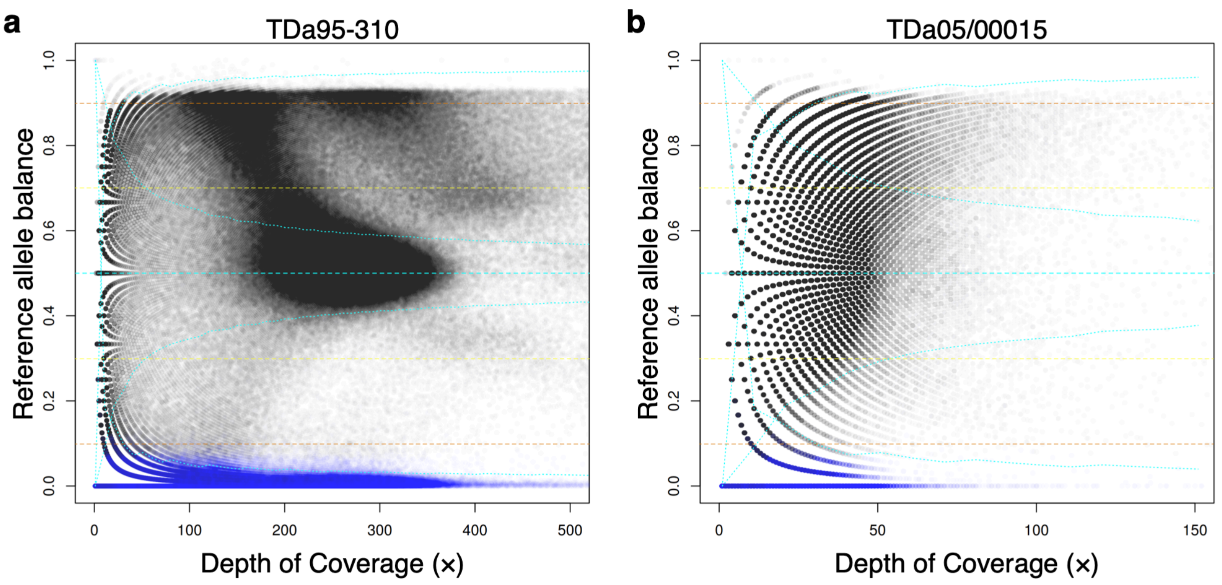


**Supplementary Fig. 14 Minimum and maximum allele-balance filters.** Thresholds imposed on heterozygous (black points) and homozygous (blue points) genotypes to mitigate the effects of repetitive sequences, artifacts, and genotyping errors. Depth-independent thresholds (yellow dashed lines) bounded heterozygous genotypes between 0.3 and 0.7, and additional thresholds (convergent cyan dotted lines) were applied using depth-dependent binomial tests ($p_0$=0.5, $p$-value > 0.001). Homozygous genotypes were similarly bounded by depth-independent thresholds at 0.1 and 0.9 (orange dashed lines), with binomial-test-based depth-dependent thresholds (divergent cyan dotted lines; $p_0$=0.01, $p$-value > 0.001). Examples of **a** TDa95-310 at high depth of coverage (×) and **b** TDa05/00015 at lower depth.

**Supplementary Table 1 Traits of parents of mapping populations.**

| ID | Seed parent[a] | Pollen parent[a] | Traits |
|---|---|---|---|
| TDa95/00328 | TDa85/00247 | TDa85/00257 | Breeding line used for genome assembly. *C. gloeosporioides* strain-specific resistance to anthracnose[3,4]. Profuse flowering and high fruit setting[5]. Purple fleshed, slight oxidation, poor tuber shape. |
| TDa00/00005 | TDa95/00328 | TDa98/00150 | Anthracnose resistant breeding line. Flowers earlier than other *D. alata*. Poor cooking quality, profuse flowering, brown-fleshed, high oxidation, high dry matter. |
| TDa02/00012 | TDa98/01166 | Unknown[b] | Breeding line with high yield and high shoot dry weight, fast tuber bulking, late senescence, creamy or white tuber flesh[6]. Anthracnose resistant. Slight oxidation, high dry matter content. |
| TDa99/00240 | TDa95/00328 | TDa98/01460 | Breeding line with short tuber dormancy[6]. Susceptible to anthracnose. Cream fleshed, no oxidation, medium dry matter content. |
| TDa01/00039 | TDa00/00005 | TDa98/01184 | Breeding line susceptible to anthracnose. Extra early flowering, good cooking quality. |
| TDa05/00015 | TDa00/00005 | TDa98/01174 | Breeding line with creamy white tuber flesh, little or no oxidation, high dry matter content, high yield, and intermediate maturity. |
| TDa99/00048 | TDa95/00328 | TDa98/01460 | Anthracnose-tolerant breeding line. Slight oxidation, high dry matter content, intermediate maturity, cream-fleshed. |

[a]The seed and pollen parents listed here are as recorded in the IITA breeding records. These are not all consistent with what we found from sequencing data, see Fig. 4 and Supplementary Table 6. The parents of the reference genotype TDa95/00328 have previously been reported as TDa92-2 and TDa85/00257[3,4]; however, this was probably incorrect, as the parents we show here reflect the IITA yam breeding records and those in YamBase.
[b]Open pollination

**Supplementary Table 2 Component and composite linkage maps.**

| Source Inst.[a] | Pop. ID | Num. Genotyped Progeny | Num. Mapping Progeny | Num. Total Markers | Num. Unique Markers | Num. QTL Markers | Total Map Length | Tau Corr. |
|---|---|---|---|---|---|---|---|---|
| IITA | TDa1401 | 317 | 294 | 5,191 | 2,182 | 2,749 | 1,645.0 | 0.9567 |
| IITA | TDa1402 | 71 | 65 | 7,505 | 1,185 | 2,181 | 1,766.2 | 0.9378 |
| IITA | *TDa1403 | 123 | 107 | 5,186 | 1,385 | 2,169 | 1,687.2 | 0.9224 |
| IITA | TDa1419 | 288 | 265 | 4,552 | 1,987 | 2,577 | 1,634.5 | 0.9565 |
| IITA | *TDa1427 | 164 | 150 | 5,797 | 1,771 | 2,643 | 1,771.3 | 0.9438 |
| NRCRI | TDa1401B | 149 | 132 | 3,447 | 1,267 | 1,707 | 1,655.1 | 0.9510 |
| NRCRI | TDa1506 | 51 | 33 | 3,440 | 522 | 1,002 | 1,682.9 | 0.9210 |
| NRCRI | TDa1512 | 127 | 117 | 4,004 | 1,433 | 1,644 | 1,641.8 | 0.9617 |
| NRCRI | TDa1603<br>TDa1610 | 32 | 25 | 2,323 | 420 | 802 | 1,626.5 | 0.9020 |
| NRCRI | TDa1621 | 84 | 69 | 3,872 | 899 | 1,494 | 1,743.1 | 0.9533 |
| IITA<br>NRCRI | *TDa1401<br>TDa1401B | 466 | 444 | 4,808 | 2,580 | – | 1,729.5 | 0.9454 |
| NRCRI<br>NRCRI | *TDa1419<br>TDa1610 | 320 | 291 | 4,384 | 2,028 | – | 1,619.1 | 0.9626 |
| NRCRI<br>NRCRI | *TDa1506<br>TDa1621 | 135 | 130 | 3,138 | 1,413 | – | 1,784.8 | 0.9091 |
| **Composite** | | **–** | **–** | **10,448** | **4,307** | **–** | **1,817.9** | **0.9989** |

*Source Inst.,* Institution that grew the plants and isolated DNA; *Pop. ID,* Mapping population identifier; *Num. Genotyped progeny,* The total number of putative progeny genotyped; *Num. Mapping Progeny,* Final number of progeny used for mapping, after filtering for relatedness and for identity in JoinMap; *Num. Total Markers,* Total number of markers passing segregation and post-imputation filters; *Num. Unique Markers,* Number of markers with genetically unique positions, regardless of parental genotype configuration; *Num. QTL Markers,* Number of markers with genetically unique positions and parental genotype configurations; *Tau Corr.,* Kendall's Tau Correlation coefficient.
[a]Table rows with two populations listed describe maps made using progeny from the same parental pairs, re-genotyped in a single, combined DArTseq report.
*Maps included in the composite linkage map.

**Supplementary Table 3 *D. alata* repeat class count and nucleotide abundances.**

| Class | Count | Bases |
|---|---|---|
| Unknown | 313,492 | 126,559,846 |
| LTR - Ty3/metaviridae[a] | 67,188 | 74,297,546 |
| LTR - Ty1/pseudoviridae[a] | 49,431 | 44,889,004 |
| LTR - Unknown | 42,799 | 30,661,425 |
| DNA - MULE-MuDR | 13,454 | 11,523,320 |
| LINE - L1 | 15,802 | 11,074,913 |
| Simple repeat | 180,650 | 7,313,907 |
| DNA - CMC-EnSpm | 5,971 | 3,353,258 |
| DNA - PIF-Harbinger | 4,818 | 2,403,923 |
| LTR - Caulimovirus | 3,034 | 2,087,172 |
| RC - Helitron | 3,403 | 1,737,013 |
| DNA - hAT-Ac | 2,498 | 1,494,735 |
| Low complexity | 28,053 | 1,401,595 |
| DNA - hAT-Tip100 | 1,252 | 792,122 |
| DNA - hAT-Tag1 | 1,245 | 616,245 |
| LINE - RTE-BovB | 910 | 259,153 |
| Unknown - Helitron-2 | 188 | 116,229 |
| DNA - TcMar-Pogo | 244 | 70,588 |
| LINE - I-Jockey | 413 | 64,769 |
| LTR | 49 | 41,014 |
| **Total** | **734,894** | **320,757,777** |

[a]Nomenclature based on e.g. Neumann *et al.*[7]

**Supplementary Table 4 Dioscoreaceae assembly and annotation BUSCO comparisons.**

| Species | Assembly | | | Annotation | | |
|---|---|---|---|---|---|---|
| | C | F | M | C | F | M |
| *D. alata* v2 | 96.6 (96.4) | 1.4 (1.5) | 2.0 (2.1) | 97.8 | 1.5 | 0.7 |
| *D. rotundata* v2 | 94.0 (86.5) | 2.2 (2.3) | 3.8 (11.2) | 83.0 | 9.5 | 7.5 |
| *D. dumetorum* | 96.5 | 0.7 | 2.8 | 88.3 | 5.2 | 6.5 |
| *D. zingiberensis* | 96.8 (96.2) | 0.9 (1.1) | 2.3 (2.7) | 93.1 | 2.8 | 4.1 |
| *T. zeylanicus* | 97.8 | 0.9 | 1.3 | 62.4 | 16.1 | 21.5 |

*C*, Complete; *F*, Fragmented; and *M*, Missing. Percentages of (n=1,375) BUSCOs are shown. Values for chromosome-scale scaffolds only are shown in parentheses.

**Supplementary Table 5 Pairwise synonymous substitution rate and identity matrix.**

| Species | *D. alata* | *D. rotundata* | *D. dumetorum* | *D. zingiberensis* | *T. zeylanicus* | *X. viscosa* |
|---|---|---|---|---|---|---|
| ***D. alata*** | – | 0.0638 | 0.1632 | 0.3891 | 0.8040 | 1.3721 |
| ***D. rotundata*** | 0.9739 | – | 0.1649 | 0.3846 | 0.7934 | 1.3727 |
| ***D. dumetorum*** | 0.9359 | 0.9360 | – | 0.3926 | 0.8116 | 1.3776 |
| ***D. zingiberensis*** | 0.8651 | 0.8685 | 0.8667 | – | 0.7365 | 1.3297 |
| ***T. zeylanicus*** | 0.7792 | 0.7819 | 0.7778 | 0.7925 | – | 1.3542 |
| ***X. viscosa*** | 0.7114 | 0.7140 | 0.7118 | 0.7185 | 0.7112 | – |

*D*, *Dioscorea*; *T*, *Trichopus*; *X*, *Xerophyta;* above diagonal, median synonymous substitution (Ks) rate; below diagonal, median gap-excluded nucleotide identity.

**Supplementary Table 6 Relatedness among eight sequenced breeding lines.**

| ID 1 | ID 2 | IBD0 | IBD1 | IBD2 | $2\hat{\pi}$ | $2\phi$ | Pedigree Explanation |
|---|---|---|---|---|---|---|---|
| TDa95/00328 | TDa99/00240 | 0.000 | 0.789 | 0.211 | 0.605 | 0.521 | **PO**: TDa95/00328 is parent of TDa99/00240; IBD2 implies TDa98/01460 is (distantly) related to TDa95/00328 |
| TDa99/00048 | TDa99/00240 | 0.092 | 0.649 | 0.259 | 0.584 | 0.559 | **FS**: Both share PO relationships with TDa95/00328 and TDa98/01460 |
| TDa00/00005 | TDa01/00039 | 0.000 | 0.880 | 0.120 | 0.560 | 0.489 | **PO**: TDa00/00005 is parent of TDa01/00039 |
| TDa95/00328 | TDa99/00048 | 0.000 | 0.897 | 0.103 | 0.551 | 0.462 | **PO**: TDa95/00328 is parent of TDa99/00048 |
| TDa00/00005 | TDa05/00015 | 0.000 | 0.923 | 0.077 | 0.539 | 0.471 | **PO**: TDa00/00005 of parent of TDa05/00015 |
| TDa95-310 | TDa00/00005 | 0.000 | 0.946 | 0.054 | 0.527 | 0.523 | **PO**: TDa95-310 is inferred parent of TDa00/00005 |
| TDa95/00328 | TDa00/00005 | 0.000 | 0.968 | 0.032 | 0.516 | 0.430 | **PO**: TDa95/00328 is parent of TDa00/00005 |
| TDa01/00039 | TDa05/00015 | 0.184 | 0.667 | 0.149 | 0.482 | 0.397 | **HSFC**: Half-sibs, but TDa01/00039 has a distant secondary relationship to shared parent TDa00/00005. |
| TDa95-310 | TDa02/00012 | 0.168 | 0.788 | 0.043 | 0.437 | 0.343 | **GG**: Inferred relationship; cannot be PO relationship with IBD0 |
| TDa95/00328 | TDa05/00015 | 0.294 | 0.618 | 0.088 | 0.397 | 0.253 | **GG**: TDa95/00328 is grandparent of TDa05/00015 |
| TDa95/00328 | TDa01/00039 | 0.258 | 0.716 | 0.026 | 0.384 | 0.154 | **GG**: TDa95/00328 is grandparent of TDa01/00039 |
| TDa99/00240 | TDa05/00015 | 0.318 | 0.597 | 0.085 | 0.383 | 0.230 | **HAV**: TDa95/00328 is parent of TDa99/00240 and grandparent of TDa05/00015; HAV because TDa95-310 is not a parent to TDa99/00048 nor TDa99/000240 |
| TDa99/00240 | TDa00/00005 | 0.264 | 0.706 | 0.030 | 0.383 | 0.221 | **HS**: Half sibs, with shared parent TDa95/00328 |
| TDa95-310 | TDa01/00039 | 0.256 | 0.730 | 0.014 | 0.379 | 0.286 | **GG**: TDa95-310 is a parent to TDa00/00005; cannot be PO with significant IBD0 |
| TDa02/00012 | TDa05/00015 | 0.326 | 0.605 | 0.073 | 0.374 | 0.203 | **HAV**: TDa95-310 is a parent of TDa00/00005 and grandparent to TDa02/00012 |

| | | | | | | | |
|---|---|---|---|---|---|---|---|
| TDa99/ 00048 | TDa01/ 00039 | 0.325 | 0.605 | 0.071 | 0.373 | 0.233 | **HAV**: TDa95/00328 is parent of TDa99/00048 and grandparent of TDa01/00039 |
| TDa99/ 00048 | TDa00/ 00005 | 0.277 | 0.703 | 0.020 | 0.372 | 0.233 | **HS**: Half sibs, with shared parent TDa95/00328 |
| TDa99/ 00048 | TDa05/ 00015 | 0.331 | 0.595 | 0.074 | 0.371 | 0.218 | **HAV**: TDa95/00328 is parent of TDa99/00048 and grandparent of TDa05/00015 |
| TDa99/ 00240 | TDa01/ 00039 | 0.330 | 0.641 | 0.029 | 0.349 | 0.135 | **HAV**: TDa95/00328 is parent of TDa99/00240 and grandparent to TDa01/00039 |
| TDa99/ 00240 | TDa02/ 00012 | 0.349 | 0.622 | 0.029 | 0.340 | 0.192 | **HAV**: Shares distant relation via TDa95-310 mate, not TDa95-310 |
| TDa00/ 00005 | TDa02/ 00012 | 0.344 | 0.638 | 0.018 | 0.337 | 0.147 | **HAV**: TDa95-310 is parent of TDa00/00005 and grandparent to TDa02/00012 |
| TDa95- 310 | TDa05/ 00015 | 0.338 | 0.649 | 0.013 | 0.337 | 0.236 | **GG**: TDa95-310 is grandparent of TDa05/00015 (via TDa00/00005) |
| TDa01/ 00039 | TDa02/ 00012 | 0.424 | 0.521 | 0.054 | 0.315 | 0.156 | **HFC**: TDa95-310 is parent of TDa00/00005 and grandparent of TDa01/00039 and TDa02/00012 |
| TDa99/ 00048 | TDa02/ 00012 | 0.421 | 0.533 | 0.046 | 0.313 | 0.214 | **UN**: Unrelated in-law |
| TDa95/ 00328 | TDa02/ 00012 | 0.433 | 0.526 | 0.040 | 0.303 | 0.005 | **HS/GG/AV**: TDa95/00328 and TDa95-310 share distant relation |
| TDa95/ 00328 | TDa95- 310 | 0.590 | 0.400 | 0.010 | 0.210 | −0.007 | **UN**: Unrelated in-law |
| TDa95- 310 | TDa99/ 00240 | 0.606 | 0.391 | 0.003 | 0.199 | −0.020 | **UN**: Unrelated in-law |
| TDa95- 310 | TDa99/ 00048 | 0.622 | 0.376 | 0.002 | 0.190 | 0.026 | **UN**: Unrelated in-law |

*IBD*, identity by descent; *IBD0*, no shared haplotypes; *IBD1*, one shared haplotype; *IBD2*, two shared haplotypes; $2\hat{\pi}$, relatedness coefficient calculated as IBD2 + 2·IBD1; $2\phi$, relatedness coefficient calculated as 2·KING-robust[8] relatedness; *PO*, parent-offspring; *FS*, full-sibling; *HSFC*, half-sibling + first-cousin (non-shared parents are siblings); *HS*, half-sibling (one shared parent); *GG*, grandparent-grandchild; *AV*, avuncular; *HAV*, half-avuncular; *HFC*, Half first cousin (sharing only one grandparent); *UN*, unrelated.

**Supplementary Table 7 Species and accessions used in this work.**

| Species name | Plastid Assemblies | Main genome Assemblies | Notes |
|---|---|---|---|
| *Dioscorea abyssinica* | MG805602.1 | – | – |
| *Dioscorea alata* | MG267382.1 | – | – |
| *Dioscorea alata* | TBD* | TBD*[,a,b] | TDa95/00328 (This study) |
| *Dioscorea aspersa* | MG267381.1 | – | – |
| *Dioscorea baya* | MG805603.1 | – | – |
| *Dioscorea bulbifera* | MG267383.1 | – | – |
| *Dioscorea bulbifera* | MG805604.1 | – | – |
| *Dioscorea burkilliana* | MG805605.1 | – | Plastid partial |
| *Dioscorea cayennensis* | MG805606.1 | – | – |
| *Dioscorea collettii* | KY996495.1 | – | – |
| *Dioscorea dumetorum* | MG805607.1 | – | – |
| *Dioscorea dumetorum* | TBD* | GCA_902712375.1 | IboSweet3[1] |
| *Dioscorea elephantipes* | EF380353.1 | – | – |
| *Dioscorea esculenta* | MT818507.1 | – | – |
| *Dioscorea futschauensis* | MG267384.1 | – | – |
| *Dioscorea hirtiflora* | MG805608.1 | – | – |
| *Dioscorea nipponica* | KP404629.1 | – | Unverified |
| *Dioscorea persimilis* | MN585218.1 | – | – |
| *Dioscorea polystachya* | MG267380.1 | – | Isolate JSSY |
| *Dioscorea polystachya* | MG267378.1 | – | Isolate FJHG |
| *Dioscorea polystachya* | MG267377.1 | – | Isolate BJXS |
| *Dioscorea polystachya* | MG267376.1 | – | Isolate SDTS |
| *Dioscorea polystachya* | MG267375.1 | – | Isolate HBFL2 |
| *Dioscorea polystachya* | MG267379.1 | – | Isolate HNYT |
| *Dioscorea polystachya* | KY996494.1 | – | – |
| *Dioscorea praehensilis* | MG805609.1 | – | – |
| *Dioscorea preussii* | MG805610.1 | – | – |
| *Dioscorea quartiniana* | MG805611.1 | – | – |
| *Dioscorea rotundata* | TBD* | GCA_009730915.1[a] | TDr96-F1[2,9] |
| *Dioscorea sagittifolia* | MG805612.1 | – | – |
| *Dioscorea sansibarensis* | MG805613.1 | – | – |
| *Dioscorea schimperiana* | MG805614.1 | – | – |
| *Dioscorea togoensis* | MG805615.1 | – | – |
| *Dioscorea villosa* | KY085893.1 | – | – |
| *Dioscorea zingiberensis* | KP899622.1 | GCA_014060945.1[a] | – |
| *Trichopus zeylanicus* ssp. *travancoricus* | MK674169.1 | GCA_005019695.1 | 2n=28; ~850 Mb genome[10] |
| *Apostasia shenzhenica* | MG772639.1 | GCA_002786265.1[b] | Plastid scrambled |
| *Asparagus officinalis* | KY364194.1 | GCF_001876935.1[a,b] | – |
| *Dendrobium catenatum* | KX507360.1 | GCF_001605985.2[b] | – |

| | | | |
|---|---|---|---|
| *Phalaenopsis equestris* | JF719062.1 | GCF_001263595.1[b] | – |
| *Musa acuminata* | HF677508.1 | GCF_000313855.2[a,b] | – |
| *Elaeis guineensis* | JF274081.1 | GCF_000442705.1[a,b] | – |
| *Phoenix dactylifera* | GU811709.2 | GCF_000413155.1[b] | – |
| *Ananas comosus* | NC_026220.1 | GCF_001540865.1[a,b] | – |
| *Oryza sativa* | KM103369.1 | GCF_001433935.1[a,b] | – |
| *Zea mays* | NC_001666.2 | GCF_000005005.2[a,b] | – |
| *Xerophyta viscosa* | MK279914.1 | GCA_002076135.1 | . |
| *Lilium candidum* | NC_042399.1 | – | No nuclear genome |
| *Japonolirion osense* | NC_036154.1 | – | No nuclear genome |
| *Acorus tatarinowii* | NC_045294.1 | – | No nuclear genome |
| *Zostera marina* | MF370229.1 | GCA_001185155.1[b] | – |
| *Spirodela polyrhiza* | MN419335.1 | GCA_001981405.1[a] | JGIv1.2 annotation mapped to GCA_001981405.1 |
| *Arabidopsis thaliana* | NC_000932.1 | GCF_000001735.4[a,b] | – |
| *Amborella trichopoda* | NC_005086.1 | GCF_000471905.2[b] | – |

[a]Chromosome-scale assembly
[b]Protein-coding gene annotation deposited in NCBI.
*Sequences new to this study.

**Supplementary Table 8 Parameters used for mapping in JoinMap.**

| Source Inst. | Pop. ID | Min. LOD | SA | GS |
|---|---|---|---|---|
| IITA | TDa1401 | 6 | 7,500/75,000(Chrs 17, 20)<br>5,000/50,000(Chr 4)<br>3,000/30,000(all others) | 50,000/5,000(Chrs 17, 20)<br>10,000/1,000(all others) |
| IITA | TDa1402 | 7 | 7,500/75,000(Chr 4)<br>5,000/50,000(Chrs 3, 13, 14, 17)<br>3,000/30,000(all others) | 50,000/5,000(Chr 4)<br>25,000/2,500(Chr 3)<br>10,000/1,000(all others) |
| IITA | TDa1403 | 8[a] | 7,500/75,000(Chr 4)<br>3,000/30,000(all others) | 50,000/5,000(Chr 4)<br>10,000/1,000(all others) |
| IITA | TDa1419 | 6 | 7,500/75,000(Chr 6, 7)<br>3,000/30,000(all others) | 50,000/5,000(Chrs 6, 7)<br>10,000/1,000(all others) |
| IITA | TDa1427 | 6 | 5,000/50,000(Chr 8, 9)<br>3,000/30,000(all others) | 10,000/1,000 |
| NRCRI | TDa1401B | 7 | 7,500/75,000(Chr 2, 17, 20)<br>3,000/30,000(all others) | 10,000/1,000 |
| NRCRI | TDa1506 | N/A[b] | 7,500/75,000(Chrs 6, 7)<br>3,000/30,000(all others) | 50,000/5,000(Chrs 6, 7)<br>10,000/1,000(all others) |
| NRCRI | TDa1512<br>TDa1603 | 7[a] | 10,000/100,000(Chrs 6, 13, 20)<br>7,500/75,000(all others) | 75,000/7,500(Chrs 6, 13, 20)<br>50,000/5,000(all others) |
| NRCRI | TDa1610 | N/A[b] | 7,500/75,000(Chrs 2, 4, 7, 8, 13, 16)<br>3,000/30,000(all others) | 50,000/5,000(Chrs 2, 4, 7, 8, 13, 16)<br>10,000/1,000(all others) |
| NRCRI | TDa1621 | 7 | 7,500/75,000(Chrs 6, 8, 19)<br>3,000/30,000(all others) | 10,000/1,000 |
| IITA<br>NRCRI | TDa1401<br>TDa1401B | 8 | 1,000/10,000 | 10,000/1,000 |
| NRCRI<br>NRCRI | TDa1419<br>TDa1610 | 9 | 10,000/100,000(Chr 6)<br>1,000/10,000(all others) | 65,000/6,500(Chr 6)<br>10,000/1,000(all others) |
| NRCRI<br>NRCRI | TDa1506<br>TDa1621 | 7 | 7,500/75,000(Chrs 4, 6, 16)<br>1,000/10,000(all others) | 50,000/5,000(Chrs 4, 6, 16)<br>10,000/1000(all others) |

*Source Inst.,* Institution that grew the plants and isolated DNA; *Pop. ID,* Mapping population identifier; *Min. LOD.*, Minimum log of odds threshold for evidence of linkage (marker grouping); *SA*, Simulated annealing parameters (map order optimization): Chain length/stop after # chains without improvement; *GS*, Gibbs sampling parameters (MC-EM multipoint ML estimation of rec. frequencies): Length of burn-in chain/chain length per EM cycle
[a]Applies to chromosomes 2–14, 16–20
[b]The genomic sequence was used to group markers into linkage groups.

### Supplementary Methods

#### Supplementary Note 1: Chromosome counting, genome sequencing, and assembly

##### Chromosome preparations

Mitotic metaphase spreads of *D. alata* (line TDa 95/00328) were prepared using dropping technique according to ref.[11], with minor modifications. Protoplast suspension was prepared from seven actively growing roots. Meristematic regions of root tips were incubated in mixture of 2% pectinase and 2% cellulase (in 75 mM KCl and 7.5 mM EDTA, pH 4) for 90 min at 37 °C, filtered through 150 µm nylon mesh and pelleted by centrifugation. Pellets were washed three times in 75 mM KCl and 7.5 mM EDTA solution followed by washing in 70% ethanol. 5 µl of protoplast suspension was dropped onto microscopic slide, followed by adding 14 µl of fixative solution (3:1) to induce protoplast bursting. Finally, slides were air dried and mounted in Vectashield mounting solution with DAPI (Vector Laboratories, Ltd, Peterborough, UK). Chromosome spreads were examined with Axio Imager.Z2 microscope (Cale Zeiss, Oberkochen, Germany) equipped with Cool Cube 1 camera (Metasystems, Altlussheim, Germany) and analyzed with ISIS v5.4.7. software (Metasystems). Final **Supplementary Fig. 2** was edited in GIMP v2.8 (GNU Image Manipulation Program, https://www.gimp.org/).

##### Genome sequencing

**Isolation of high molecular weight DNA for PacBio sequencing**
TDa95/00328 plants sourced from IITA Ibadan, Nigeria were grown at World Agroforestry (CIFOR-ICRAF) in Nairobi, Kenya. Four fresh leaves weighing around 1 g were ground using mortar and pestle in liquid nitrogen. To the ground powder, 25 mL of preheated (65 °C) high salt CTAB buffer (100 mM Tris-Cl, pH 8.0; 20 mM EDTA, pH 8.0; 3 M NaCl; with freshly added 3% polyvinylpyrrolidone (PVP), 2-mercaptoethanol, and 3% CTAB) was added and kept in 65 °C water bath for 45 min with intermittent gentle shaking. The heated solution was cooled in an ice bath, 1/2 vol of 5 M NaCl was added followed by the addition of 1/10 vol of chilled isopropanol, and the solution was gently mixed. It was then centrifuged at 3,500 *g* in a refrigerated macrocentrifuge for 20 min at 4 °C. The clear supernatant was separated and purified by mixing with an equal volume of dichloromethane and centrifuged as before. It was further purified using an equal volume of chloroform: isoamyl alcohol (24:1) and separated using the centrifuge as described. The clear supernatant was separated, an equal volume of isopropanol added, and stored at 4 °C (or −20 °C if needed) for 1–3 h for DNA precipitation. The DNA was pelleted by centrifugation as before, washed with a sufficient amount of 70% ethanol, and dried at room temperature. The dried pellet was dissolved in a minimum volume (preferably less than 500 µL) of TE (10 mM Tris-Cl, pH 8.0; 1 mM EDTA, pH 8.0). The dissolved DNA was treated with 10 µL of RNase (10 mg/mL) at room temperature for 1 h, gently mixed with an equal volume of chloroform:isoamyl alcohol (24:1) and centrifuged in a refrigerated microfuge at 13,000 *g* for 15 min at 4 °C. The supernatant was separated, two volumes of chilled absolute ethanol were

added, and stored in −20 °C for 1–2 h for DNA precipitation. The precipitated DNA was pelleted by centrifugation in the refrigerated microfuge as before. The pellet was washed with 70% ethanol, dried at ambient temperature, and dissolved in a minimum volume of TE, preferably less than 300–400 µL. The dissolved DNA was quality checked on 0.8% agarose gel for DNA integrity, optical density ratios were checked using Nanodrop, and DNA was quantified using fluorometric method using Qubit. This DNA was used for Pacific Biosciences (PacBio, Menlo Park, USA) Single-Molecule Real-Time (SMRT) continuous long-read (CLR) sequencing.

**Generation of mate pair data**

Fresh leaf samples from TDa95/00328 were collected from the field at IITA and kept on ice. 100 mg of leaf sample was placed in a 2.0 mL Eppendorf tube, and ground with liquid nitrogen. To remove secondary metabolites, it was washed 2–3 times by adding 1000 µl HEPES buffer [0.1 M HEPES, PVP, L-ascorbic acid, 2-mercaptoethanol, and sterile distilled water], mixing thoroughly, centrifuging at 13,000 rpm for 2 min, and decanting the supernatant. Next, 800 µl of freshly prepared CTAB buffer [1M Tris HCl pH 8, 0.5 EDTA pH 8, 5M NaCl pH 8, 1% mercaptoethanol, 3% CTAB] was added and mixed well to ensure homogenization, and samples were incubated for 30 min at 65 °C in a water bath. 600 µl of chloroform-isoamyl alcohol (24:1) was added and the samples were mixed gently and centrifuged at 13,000 rpm for 5 min. The aqueous phase was transferred into freshly labeled 1.2 mL tubes. DNA was precipitated with the addition of ⅔ vol ice-cold isopropanol, mixed by inverting, incubated at −20 °C for 1 h, and centrifuged at 13,000 rpm for 5 min. The supernatant was decanted carefully, and the DNA pellet was washed twice with 500 µl of cold 70% ethanol. The ethanol was drained completely and the pellet dried at 37 °C for 30 min. DNA was resuspended in 50 µl of low TE (1 mM Tris, 0.1 mM EDTA) and 3 µl RNase, incubated at 37 °C for 1 h, and stored at 4 °C.

Three Illumina Nextera mate-pair libraries were prepared at the UC Davis Genome and Biomedical Sciences Facility with the Illumina Nextera Mate Pair Library Prep Kit according to the manufacturer's instructions, and incorporating gel-based size selection. Before library preparation DNA was cleaned with high salt Phenol/Chloroform. Fragment peaks were 2583 bp, 5903 bp, and 9033 bp, as measured via Bioanalyzer 2100 (Agilent). 71M, 56M, and 49M 151 bp read pairs, respectively, were sequenced on HiSeq 4000.

**Generation of HiC data**

Isolation of formaldehyde-fixed nuclei**:** *D. alata* (TDa95/00328) tubers were transferred to soil and kept in a greenhouse at the Institute of Experimental Botany, Olomouc, Czech Republic. Young leaves and apical parts of the stem were used to prepare suspensions of intact nuclei according to ref.[12], with some modifications. Briefly, fresh plant material was cut into approximately 0.2–0.5 cm long fragments and incubated in 2% (vol/vol) formaldehyde solution for 20 min at 5 °C, then washed three times in Tris buffer for 5 min at 5 °C. Plant material was excised and chopped with a razor blade in 1 mL IB buffer on a Petri dish. The crude homogenate was filtered through a 50 mm pore size mesh and suspensions of nuclei were centrifuged at 300 *g* for 5 min at 4 °C. The supernatant was discarded and the pellet resuspended in 500 mL IB buffer containing DAPI (0.2 mg/mL final concentration). Flow cytometry experiments were carried out on FACSAria II SORP flow cytometer and sorter (BD Biosciences) equipped with two lasers (488 nm and 355 nm) and optical detectors with appropriate optical filters. Four batches of 1,000,000 nuclei at the G1 phase of the cell cycle were flow sorted into 2 mL Eppendorf tubes containing 100 µL IB buffer. The samples were then centrifuged at 500 *g* for 30 min at 4 °C, the supernatant was removed, and pelleted nuclei were kept on dry ice for further use. These nuclei were sent to Dovetail Genomics.

A HiC library was prepared at Dovetail Genomics in a similar manner as described previously[13]. Briefly, for each library, formaldehyde-crosslinked chromatin was extracted and digested with DpnII, the 5' overhangs filled in with biotinylated nucleotides, and then free blunt ends were ligated. After ligation, crosslinks were reversed and the DNA purified from protein. Purified DNA was treated to remove biotin that was not internal to ligated fragments. The DNA was then sheared to ~350 bp mean fragment size and sequencing libraries were generated using NEBNext Ultra enzymes and Illumina-compatible adapters. Biotin-containing fragments were isolated using streptavidin beads before PCR enrichment of each library. The libraries were sequenced on an Illumina HiSeq 4000 to produce 358.5 million 2×151 bp paired-end reads.

A listing of all TDa95/00328 sequencing data, and corresponding NCBI SRA accession numbers, may be found in **Supplementary Data 1**.

#### De novo genome assembly

In order to obtain a sequence-based estimate of genome size, we performed *k*-mer counting methods on 65.6 Gb of TDa95/00328 Illumina WGS reads. First, contaminant reads were identified by aligning all reads with BWA-MEM[14] to an intermediate PacBio-based contig assembly (detailed below) to which published plastid[15], mitochondrial[2], and Phi X[16] sequences were added. If either read (or both) in a pair mapped to a contig annotated as a contaminant, the pair was subtracted from the dataset. The filtered reads were then *k*-mer counted with Jellyfish[17] v2.2.10 with *k*=51, and the *D. alata* genome size estimated using the kmerHistAnalyzer.pl script included with the Meraculous (v2.2.6) assembly pipeline[18,19].

The longest 110× of PacBio CLR reads (50.228 Gb in reads 19.8 kb or longer) were assembled with Canu[20] v1.7-221-gb5bffcf (genomeSize=600m corOutCoverage=90 minReadLength=5000 minOverlapLength=1000 corMhapSensitivity=normal obtMhapSensitivity=normal batOptions="-dg 3 -db 3 -dr 1 -ca 500 -cp 50"). These contigs represented an incomplete subset of both TDa95/00328 haplotypes and were subsequently filtered down to a single haploid complement of the genome via manual curation in JuiceBox[21,22] v1.8.9, taking into account both median contig depth (**Supplementary Fig. 13**) and sequence homology. SSPACE[23] v3 was then used (-n 1000000000 -k 3) to order and orient the 532 remaining non-redundant contigs into scaffolds with 2 kb, 4 kb, and 7 kb mate-pair sequences. These Illumina Nextera mate-pair datasets were each preprocessed using nxtrim[24] v0.4.2, aligned to the contigs with BWA-MEM, and duplicate read pairs removed using Picard v2.16.0 (http://broadinstitute.github.io/picard). Draft chromosome-scale super-scaffolds were constructed with the 3D-DNA (commit 2796c3b;

params: run-asm-pipeline.sh -r 0 -i 20000) HiC scaffolding pipeline[25]. Lastly, we leveraged genetic linkage information (from preliminary, parent-specific linkage maps of TDa1401, TDa1403, TDa1419, and TDa1427 available at the time) to identify large-scale misassemblies caused by ambiguities in order or orientation that could not be readily resolved using the HiC alone. Following each step in the scaffolding process, misassemblies were corrected manually in JuiceBox. HiC files required by JuiceBox were prepared using the Juicer[26] v1.5.6 pipeline.

The assembly was screened for contaminants using BLAST+[27,28] v2.10.0 nucleotide searches against the NCBI NT database (obtained Jan 22, 2019). Contigs with best-hits (-evalue 1e-10 -num_alignments 1) to known plastid or non-plant sequences were removed. Notably, a complete *Pseudomonas synxantha* genome sequence, a rhizosphere-dwelling commensal bacterium producing nematicidal toxins[29], was identified among the contaminant sequences.

Finally, the assembly was polished twice with the Arrow[30] signal-based long-read polishing tool (v2.2.2 provided in the SMRT Link v6.0.0.47841 package) using all reads at least 5000 bp long (243× depth) aligned with BLASR[31] v5.3. Residual errors were corrected by two subsequent rounds of Illumina-based polishing with the combined 196× of TDa95/00328 Illumina short-insert paired reads aligned with BWA-MEM. Polishing was performed by calling variants on properly-paired[32] reads with FreeBayes[33] v1.1.0-54-g49413aa (--use-mapping-quality --report-genotype-likelihood-max --min-base-quality 20 --min-mapping-quality 30 --strict-vcf --use-best-n-alleles 3 --haplotype-length 2), selecting variants with reliable calls (between 10–140× depth), and then patching the assembly with a custom script (ILEC v0.1.3; https://bitbucket.org/rokhsar-lab/map4cns). In the final assembly, six chromosomes have telomeric repeats capping both ends (chromosomes 1, 2, 8, 9, 19, and 20), 12 have telomere sequences at only one end (chromosomes 3, 5–7, 10–15, 17, and 18), and the remaining two chromosomes have no telomeric repeat (chromosomes 4 and 16).

#### Supplementary Note 2: Genetic linkage mapping

##### DArTseq genotyping

DNA isolation at IITA for DArTseq genotyping was performed following the same protocol as that used for mate-pair libraries (**Supplementary Note 1**). The sequencing data used for DArTseq genotyping are provided with NCBI SRA accessions in **Supplementary Data 1**.

Using a modified CTAB method[34], DNA extraction was performed on 333 progeny maintained at NRCRI in Umudike, Nigeria. Briefly, lyophilized yam leaf samples were ground to powder in a Qiagen TissueLyser LT for 1 min at a rate of 1500 strokes/min and transferred to 2 mL microtubes. The ground tissue was homogenized in 800 µl of CTAB buffer (100 mM Tris-HCl pH 8.0, 20 mM EDTA pH 8.0, 1.4 M NaCl, 1% polyvinyl pyrrolidone, 2% 2-mercaptoethanol, 3% CTAB), then incubated for 30 min at 65 °C. 600 µl of an equal volume of chloroform and isoamyl alcohol (24:1 vol/vol) was added to the tube and centrifuged for 10 min at 13000 rpm. The nucleic acid in the aqueous phase was precipitated out with cold isopropanol, and the pellets washed by centrifuging at 13000 rpm with 70% ethanol. The pellets were further suspended in 50 µl of sterile water and treated with 3 µl of RNAse A (20 mg/mL) for 1 h at 37 °C. Finally, the samples were stored at −20 °C until use. The DNA samples were quantified using a NanoDrop 1000 (Thermo Scientific) and their integrity assessed by agarose gel electrophoresis.

#### DArTseq quality control and variant filtering

For each of the ten genotyped datasets, the initial DArTseq genotype calls, obtained from DArT or IGSS and provided in either single-row or two-row CSV format (**Supplementary Data 2**), were converted to VCF with MapTK[35] (https://bitbucket.org/rokhsar-lab/gbs-analysis) v1.4.1-6-gd1540e4 using the DArT2VCF subcommand, then mapped onto the v2 genome sequence using the MapTags subcommand. MapTK-MapTags uses BWA-MEM[14] v0.7.17-11-g20d0a13 to align the 69-bp sequence tag accompanying each genotyped locus, requiring one of the two alleles at the locus to match the genome sequence, then re-indexes the genotyped alleles accordingly. Only uniquely-mapped sequence tags (*i.e.*, those with mapping quality values equal to 60) were retained for downstream analyses. A first-pass filtering round was then performed to select loci that fit Mendelian patterns of segregation for both allele and genotype frequencies using goodness-of-fit tests ($\chi^2$ test, $p$-value ≥ $1{\times}10^{-2}$) implemented by the MapTK-ChiF1 subcommand. These provisional segregating $F_1$ loci were filtered further for genotyping completeness using VCFtools[36] v0.1.16-16-g954e607, requiring at least 90% of samples to be genotyped per locus. Filtered loci were then used to perform relatedness analyses, as in refs.[8,35]. Based on these analyses, half-sibs and off-types were identified and removed.

For four of the populations (TDa1402, TDa1506, TDa1610, and TDa1621), the intended TDa02/00012 parent samples genotyped were, in fact, not truly TDa02/00012. The TDa02/00012 samples from three other crosses (TDa1403, TDa1419, and TDa1427), however, shared genetic identity with one another and relatedness values consistent with true paternity to the progeny of their respective populations. Consensus genotypes were generated by majority-rule from these validated TDa02/00012 samples and used to replace the invalidated ones. Paternity between this consensus TDa02/00012 sample and the progeny was then verified by repeating relatedness analyses for each of the four populations. Furthermore, we observed that the sample genotyped as TDa99/00240, the intended seed parent to the TDa1610 population, did not actually share relatedness consistent with true parentage. TDa99/00240 genotypes provided with the TDa1419 population were substituted and parentage was verified by repeating the relatedness analysis.

With parentage validated and only full-sib progeny retained, each population was refiltered for loci that 1) have a 90% genotyping call rate or higher, 2) pass all Mendelian segregation goodness-of-fit tests ($\chi^2$ test, $p$-value ≥ $1{\times}10^{-3}$), and 3) were annotated by MapTK-ChiF1 with a boolean 'P0PHASED' key in the VCF INFO field, indicating that both parental genotypes could be inferred and at least one matched to its parent of origin. Genotypes were then phased and imputed using AlphaFamImpute[37] v0.1, and MapTK-ChiF1 was run on the post-imputed VCFs to identify poorly-imputed loci, which were then removed with VCFtools. Each VCF was converted to locus format with MapTK-VCF2Loc for linkage mapping with JoinMap.

#### Genetic linkage map construction

##### Population-specific linkage maps

A parent-averaged linkage map was constructed for each population using JoinMap[38,39] v4.1 with the maximum-likelihood mapping function for cross-pollinated (CP) populations. Loci and individuals identified as identical were removed.

We identified the locations of the markers in the genome assembly and used these at times to guide LOD choice for proper linkage group formation in JoinMap. In general, the highest LOD was chosen that resolved 20 linkage groups corresponding to the 20 *D. alata* chromosomes. The same LOD was selected for all linkage groups in a JoinMap project for a given cross (or cross pair). In some cases, individual chromosome maps were made separately because no LOD separated their markers properly in JoinMap (either the chromosome was merged with another, or at a LOD that properly split other groups it also was split). In these cases, the same individuals were excluded as in the main project for that population, except two individuals were not excluded from chromosome 1 of NRCRI cross TDa1512/1603.

Additional maps were made using all progeny from the following pairs of mapping populations that shared the same parents: TDa1401 and TDa1401B; TDa1419 and TDa1610; TDa1506 and TDa1621; TDa1512 and TDa1603.

These chromosomes were mapped individually: TDa1403 Chr01, 15; TDa1512/1603 Chr01, 15; TDa1506 all chromosomes; TDa1610 all chromosomes. Chromosome maps made from individual projects were grouped using LOD 2. We did not succeed in making a plausible map of chromosome 5 from TDa1610 or TDa1506.

Loci with identical recombination fractions (i.e., redundant genetic positions) as another locus were identified in JoinMap and excluded in order to ease the computational time required to perform the locus clustering and mapping functions. These markers were later added back to each map in the location of their nearest genomic neighbor locus (see **Supplementary Table 2**). For most settings, default parameters were used, except for those noted in **Supplementary Table 8**. For these, a baseline set of parameters was used for each mapping population and then the maps were compared to the genome assembly for collinearity. If it appeared there were order and/or distance-inflation errors in the map for a given chromosome, the mapping was repeated with more stringent parameters. For populations TDa1401 and TDa1401B, individual maps were used for QTL analysis; the combined map was used as a component map for generating the composite map.

##### Composite mapping with LPmerge

We constructed a composite map using LPmerge[40] v1.7, for each chromosome choosing the linkage group with the least root mean-squared error (RMSE) over the ‘max.interval’ parameter range (1–10) tested. Each input component map was weighted by the number of progeny in its population. To balance computational constraints while capturing the most unique markers and recombinations, we combined the fewest maps that provided the greatest coverage of the parents. We first ranked maps in descending order by progeny count, then chose maps progressively from the top of this list, adding maps to include each distinct parent at least once. For validation, we confirmed the composite map was highly collinear with all component maps (**Supplementary Table 2**) and the chromosome-scale assembly. Component and composite linkage maps may be found in **Supplementary Data 3**.

#### Supplementary Note 3: RNA sequencing and genome annotation

##### RNA Sequencing

###### Tissue collection and RNA extraction

RNA samples from 12 tissues were collected, extracted, and pooled from a single TDa95/00328 plant growing on-site at ICRAF in Nairobi, Kenya. The total RNA was extracted from leaf petiole, roots, various stages of leaves (initial sprouting leaf, leaf bud, young leaf, semi-matured leaf, matured leaf, fifth leaf), bark, stem, first internode, and middle vine, using the Purelink RNA mini kit following the kit recommendations (https://assets.thermofisher.com/TFS-Assets/LSG/manuals/MAN0019350_RNA_Mini_plant_tissue_QR.pdf). In short, approximately 100–200 mg of tissue was ground in liquid nitrogen, mixed with 800 µL–1 mL of lysis buffer (containing externally added 2-mercaptoethanol at 10% vol/vol), and centrifuged in a refrigerated microfuge at 12,000 *g* for 15 s at room temperature. The supernatant was separated, 1.5 volumes of absolute ethanol were added, 700 µL of this solution loaded into the binding column and centrifuged as before, the flow-through discarded and the process repeated until the entire supernatant had been run through the column. The column was washed once with 350 µL of Wash Buffer I followed by centrifugation as before, treated with 80 µL of PureLink DNase Mixture, incubated at room temperature for 15 min, washed with 350 µL Wash Buffer I, centrifuged at 12,000 *g* for 15 s at room temperature, and washed twice with 500 µL of Wash Buffer II (with externally added absolute ethanol at 80% vol/vol) by centrifuging at 12,000 *g* for 15 s at room temperature. The binding column was dry spun at 12,000 *g* for 1 min at room temperature. A volume of 100 µL of RNA-free deionized water was added to the center of the column and incubated for 1 min at room temperature. The total RNA was finally eluted by centrifugation at 12,000 *g* for 2 min at room temperature. The RNA was separated on a 0.8% agarose gel to check RNA integrity, optical density ratios were checked using Nanodrop, and RNA was quantified using a fluorometric method with Qubit. In addition, RIN values for each sample were determined using Bioanalyzer. All the samples had a RIN value (RNA Integrity Number) above 7 except the roots (6.8) and stem (6.9). All the RNA samples were pooled and used for Illumina RNAseq library construction as well as nanopore direct RNA sequencing. The Nanopore DRS and Illumina RNAseq reads were deposited with SRA accessions SRR13683864 and SRR13683865, respectively.

##### Illumina strand-specific RNAseq

Illumina RNAseq library preparation and sequencing were performed at the Agricultural Research Council Biotechnology Platform (ARC-BTP) in Pretoria, South Africa. Library preparation was performed using the Illumina TruSeq stranded mRNA sample preparation kit (Illumina cat# 20020594), per the manufacturer's recommendations. Sequencing was performed on an Illumina HiSeq 2500 with v4 chemistry (2×125 bp). A total of 41 million read pairs were generated.

##### Nanopore Direct RNAseq

Oxford Nanopore Technologies (ONT) Direct RNA Sequencing (Nanopore DRS) and data processing were performed at the University of Dundee, Dundee, UK. Total RNA was precipitated in 2.5 M ammonium acetate pH 5.2 and 33.3% ethanol overnight at −20 °C. Next, the sample was centrifuged at 16,100 *g* for 30 min at 4 °C and washed twice with 75% ethanol (centrifugation at 16,100 *g* for 5 min at 4 °C). Nuclease-free water (ThermoFisher Scientific) was used to resuspend the RNA pellet. The total RNA concentration was measured using a Qubit 1.0 Fluorometer and Qubit RNA BR Assay Kit (ThermoFisher Scientific), while RNA quality and integrity were assessed using a NanoDrop™ 2000 spectrophotometer (ThermoFisher Scientific) and Agilent 2200 TapeStation System (Agilent). Nanopore DRS library preparation was performed using the SQK-RNA001 kit (ONT) as previously described[41] using 5 μg of total RNA as an input for library preparation. The library was loaded onto R9.4 SpotON Flow Cells (ONT) and sequenced using a 48 h runtime. We performed one replicate that generated 626,000 reads.

Nanopore DRS reads were base-called using Guppy v2.3.1 (ONT). Illumina RNAseq 2×150 bp reads were merged using FLASH[42] v1.2.11, with an expected average fragment length of 170 bp and expected fragment length standard deviation of 50 bp. Merged Illumina reads were then used to error correct the Nanopore DRS reads, using proovread[43] v2.14.1 without sampling. Corrected reads were aligned to the genome assembly with Minimap2[44] v2.8, using the 'splice' preset. Aligned reads were used to perform transcript assembly using pinfish v0.1.0 (ONT).

#### Genome annotation procedure and comparison

##### Protein-coding gene annotation

Transcript assemblies were made from roughly 107M pairs of paired-end Illumina RNA-seq reads—sourced from the above-described pooled library, ref.[45] (SRA: SRR1518381 and SRR1518382), and ref.[46] (SRA: SRR3938623)—and approximately 44k 454 ESTs[47] (SRA: SAMN00169815, SAMN00169801, SAMN00169798) using PERTRAN (Shengqiang Shu, unpublished). A total of 86,399 transcript assemblies were constructed using PASA[48] from 18 full-length cDNAs collected from NCBI, roughly 53k collapsed assemblies from short-read corrected Nanopore DRS reads, and the RNA-seq transcript assemblies described above.

Gene loci were determined by transcript assembly alignments and/or EXONERATE[49] alignments of proteins from *Arabidopsis thaliana*[50] TAIR10, *Glycine max*[51] Wm82.a4.v1, *Sorghum bicolor*[52] v3.1.1, *Oryza sativa*[53] v7.0, *Setaria viridis*[54] v2.1, *Amborella trichopoda*[55] v1.0, *Zostera marina*[56] v2.2, *Musa acuminata*[57] v1, *Ananas comosus*[58] v3, *Vitis vinifera*[59] v2.1 proteomes obtained from Phytozome v13 (https://phytozome-next.jgi.doe.gov), and Swiss-Prot[60] (2018, release 11) proteome. Alignments were performed against the repeat soft-masked TDa95/00328 genome (see below) with up to 2 kb extension on both ends unless extending into another locus on the same strand. Gene models were predicted using homology-based predictors: FGENESH+[61], FGENESH_EST (similar to FGENESH+, but uses ESTs to compute splice site and intron input instead of protein/translated ORF), EXONERATE, PASA assembly-derived ORFs (an in-house homology-constrained ORF finder), and from AUGUSTUS via BRAKER1[62]. The best-scored predictions for each locus were selected using multiple positive factors, including EST and protein support, and one negative factor (i.e., overlap with repeats).

The selected gene predictions were improved by PASA. Improvements included adding UTRs, splicing correction, and adding alternative transcripts. PASA-improved gene model proteins were subjected to protein homology analysis to the above-mentioned proteomes to obtain C-score and protein coverage. The C-score is a protein's BLASTP[63] score ratio to its mutual best hit (MBH) BLASTP score, and protein coverage is the highest percentage of protein aligned to the best of its homologs. PASA-improved transcripts were selected based on C-score, protein coverage, EST coverage, and their coding sequences (CDS) overlapping with repeats. The transcripts were selected if their C-scores were larger than or equal to 0.5 and protein coverage larger than or equal to 0.5, or they had EST coverage but their CDS overlapped repeats were less than 20% of their lengths. For gene models whose CDS overlapped with repeats more than 20%, their C-scores had to be at least 0.9 and homology coverage at least 70% to be selected. The selected gene models were then subjected to Pfam analysis: gene models whose protein was annotated with more than 30% transposable element domains were removed. Incomplete gene models, those with low homology support without full transcriptome support and short single-exon (< 300 bp CDS) models without protein domains nor good expression were manually filtered out.

Annotation completeness was measured using the BUSCO[64] v3.0.2-11-g1554283 pipeline with the Embryophyta OrthoDB[65] v10 database. Among the ten (0.7%) genes that the pipeline determined were missing, three could be partially aligned to the assembly with EXONERATE[49] v2.4.0, and one was aligned at full-length, but with an internal frameshift. No alignments could be found for the remaining six sequences. Only six of the ten missing/fragmented genes could be characterized with BLASTP[28] searches against the NCBI NR database, and included one of each of 1) 5'-nucleotidase domain-containing protein DDB_G0275467 isoform X1; 2) F-box/kelch-repeat protein OR23; 3) deoxycytidylate deaminase; 4) D-aminoacyl-tRNA deacylase isoform X2; 5) ferredoxin, root R-B2; and 6) phosphoglucan phosphatase LSF1, chloroplastic isoform X1.

**Genomic repeat annotation**

A first-pass repeat annotation was performed to aid the protein-coding gene annotation. The RepeatMasker[66] repeat library used consists of *de novo* repeats inferred by RepeatModeler[67] (v1.0.11) on the TDa95/00328 genome (v1, sourced from Phytozome v13 via https://phytozome-next.jgi.doe.gov) combined with *Dioscorea* repeats deposited in RepBase[68].

A second annotation of low-complexity sequences and transposable elements was later performed with RepeatModeler v2.0.1 on the substantially more complete v2 genome sequence with the LTR structural analysis (-LTRStruct) option enabled. The resulting *de novo* repeat collection (see **Supplementary Table 3**) was used as the input library to RepeatMasker v4.1.1 (with options: -gff -rmblast), which was employed to annotate their genomic locations in GFF format and identified an additional 56.2 Mb of repetitive sequence.

**Comparisons with other monocot genomes**

OrthoFinder[69] v2.4.1 was used to perform orthologous gene clustering on the available assembled Dioscoreaceae species: *D. alata*, *D. rotundata*, *D. dumetorum*, *D. zingiberensis*, and *T. zeylanicus*. From these clusters, genes in strict 1:1:1:1 correspondence among the *Dioscorea* species were selected. There were 5,454 such clusters, of which 99.9% (n=5451), 90.5% (n=4937), and 99.1% (n=5404) of these were localized to chromosome-scale scaffolds in *D. alata*, *D. rotundata*, and *D. zingiberensis*, respectively.

OrthoFinder was also leveraged to infer orthologous gene clusters among a broader set of monocots: *D. alata, D.rotundata*, *D.dumetorum*, *D.zingiberensis*, *T. zeylanicus*, *Xerophyta viscosa*, *Apostasia shenzhenica*, *Dendrobium catenatum*, *Asparagus officinalis*, *Elaeis guineensis*, *Phoenix dactylifera*, *Musa acuminata*, *Oriza sativa*, *Zea mays*, *Ananas comosus*, *Spirodela polyrhiza*, *Zostera marina*, *Arabidopsis thaliana*, and *Amborella trichopoda*. These orthologous clusters, and those from the Dioscoreaceae-specific run, were visualized with the ClusterVenn online tool[70] and can be found in **Supplementary Fig. 6**.

Benchmarking Universal Single-Copy Orthologs (BUSCO) statistics were calculated for each of the assembled Dioscoreaceae species using the OrthoDB[65] v10 Embryophyta benchmark dataset (n=1,375) and the v3.0.2-11-g1554283 BUSCO[71] pipeline. Results of these analyses are available in **Supplementary Table 4**.

#### Supplementary Note 4: Chromosome landscape, Rabl chromatin structure, and centromere estimation

##### Chromosome landscape

The genome was divided into 500 kb non-overlapping windows with the BEDtools[72] v2.28.0 ‘makewindows’ function and the abundances of the genomic features were calculated for each with the ‘intersect’ subcommand. These 500 kb windows formed genomic vectors for each type of genomic feature that were used to calculate Pearson’s correlation coefficients (*r*) between each with the R[73] v3.5.3 ‘cor’ function. The genomic features examined included gene count, low-complexity and transposable element repeat densities, recombination rate, and A/B compartment domain status.

Gene counts and repeat densities were calculated from the protein-coding gene and repeat annotations described above. Recombination rate was calculated for each window by interpolating the genetic positions of the beginning and end of each window using the MapTK Predict subcommand[35] with two nearest neighbor map markers, and then computing the slope between those two points.

The A/B compartment structure for each chromosome (**Supplementary Fig. 7**) was inferred as follows: HiC reads were aligned to the TDa95/00328 v2 genome assembly with Juicer, as described previously, and KR-balanced[74] intra-chromosomal HiC contact matrices of observed counts were extracted at 100 kb matrix resolution with Juicer Tools (v1.8.9) at a minimum mapping quality of 30. An R script ('call-compartments' in https://bitbucket.org/bredeson/artisanal), implementing a principal component analysis (PCA)-based algorithm on intra-chromosomal Pearson's correlation matrices, was used to extract and plot the loadings of the first principal component against chromosome position. For chromosome 14, localizing the PCA along the diagonal of the correlation matrix with sliding windows (with window-size half the length of a chromosome) mitigated much of the confounding signal introduced by intra-chromosomal p-q arm contacts, thereby amplifying compartment signal. When computing correlations between A/B compartments and other genomic features, the loadings of five 100 kb windows intersecting each 500 kb non-overlapping window were averaged using the arithmetic mean.

#### Rabl chromosome structure and centromere estimates

The three-dimensional 'Rabl' conformation of chromosomes within the nucleus, first described by Carl Rabl[75], is reviewed in more detail by ref.[76]. HiC contact patterns sampled from Rabl-structured chromosomes have been observed in yeast and barley[77,78], and similar patterns were readily observed upon visualizing TDa95/00328 leaf tissue HiC data in JuiceBox[21] v1.9.0. Unlike in barley, however, 'A/B' compartment structure was the dominant layer (i.e., strongest variance component) of contact information represented in the intra-chromosomal HiC contact matrices, so contacts between chromosomes were used to model the Rabl conformations (**Fig. 1** and **Supplementary Fig. 4**).

KR-balanced[74] inter-chromosomal matrices of observed HiC contact counts were extracted at 100 kb matrix resolution for all pairwise combinations of chromosomes using Juicer Tools (v1.8.9) with a minimum mapping quality of 30. Metacentric chromosome 2 was chosen as a comparator chromosome. For any row or column with zero variance in a matrix, a jitter of $1\times10^{-4}$ was introduced. PCA was then performed on the matrix using the 'prcomp' function in R[73] v3.5.3 with scaling and centering both enabled.

The first two principal components (PCs), each corresponding to vector of 100 kb non-overlapping windows along a chromosome, were plotted against their genomic coordinates and smoothed using the built-in 'ksmooth' R function with a Gaussian smoothing kernel and the bandwidth set to 10% of the chromosome length. These smoothed points were then colored according to their distance from the estimated centromere (black), where chromosome p-arms were colored blue, and q-arms colored yellow (chr-structure.R' script in https://github.com/bredeson/Dioscorea-alata-genomics). Centromeric positions were estimated by eye in JuiceBox using the principles described in ref.[77].

#### Supplementary Note 5: Phenotyping

##### Planting scheme at IITA for phenotyping

Phenotyping of five mapping populations was performed at IITA from 2016–2019. In 2016, mapping populations were planted in single pots and grown in the screenhouse for seed tuber multiplication and screening of anthracnose disease in a controlled environment. In 2017, individual mini-tubers of each mapping population were pre-planted in pots to ensure germination, and one-month-old seedlings were transplanted in the field using a ridge-and-furrow system. Three plants per genotype were planted in ridges with a 1 m distance between the plants within a ridge and between two ridges. The mapping populations were planted following a randomized complete block design (RCBD) with three replicates for each mapping population. Land preparation, weeding, staking and harvesting were carried out following standard field operating protocol for yam as reported by Asfaw[79]. In 2018 and 2019, harvested tubers were cut into mini-setts of 100 g each and treated with pesticide (marcozeb: 70 g, chlorpyrifos: 75 mL mixed with 10 L of tap water) for 10 minutes to avoid rotting of mini-setts. Planting was carried out in the field following the same procedures as described above.

##### Phenotyping of mapping populations for anthracnose disease

###### Assessment of anthracnose disease at IITA

*Field assessment*: Each plant in the five IITA populations (TDa1401, TDa1402, 1403, 1419 and 1427) was visually scored for yam anthracnose disease (YAD) severity at 3 MAP (months after planting) and 6 MAP using a 1–5 scale as follows: scores 1, no symptoms; 2, 1–25%; 3, 25–50%; 4, 50–75%; and 5, >75%. Simple means of these scores and the incidence were calculated using the formula, 100 × number of infected plants / total number of plants. The area under the disease progression curve (AUDPC) was also calculated using the formula:

$$A_k = \sum_{i=1}^{N-1} \frac{(y_i + y_{i+1})}{2} (t_{i+1} - t_i)$$

where $A_k$ is AUDPC, $y_i$ is the disease index as a proportion of the *i*th observation, $t$ is time equivalent to days after planting, and $N$ is the total number of observations.

*Detached leaf assay* (DLA): DLA analysis was performed at IITA in 2016 on plants grown in the screenhouse, and in 2017 and 2018 on plants grown in the field. Two disease-free young leaves were detached from between the sixth and eleventh fully expanded leaf of each of the parental lines and progeny of the mapping populations. DLA screening was performed using a slightly modified protocol of ref.[80] and Nwadili *et al.*[81] as follows: The most virulent *Colletotrichum gloeosporioides* isolate identified by Nwadili *et al.*[81], Kog01R1, was grown on potato dextrose agar (PDA) amended with lactic acid for 8 days at 28 °C. The 8-day-old culture plates were washed with sterile distilled water and sieved with cheesecloth; the suspension was thereafter adjusted to a concentration of $1\times10^6$ spores per mL using a hemocytometer.

The leaves were surface sterilized in 0.5% sodium hypochlorite (vol/vol) for 2 min, followed by 3 rinses of sterile distilled water for 1 min each, and thereafter plated in petri plates lined with 2–3 layers of filter paper moistened with 1–2 mL distilled water with the adaxial surface down. The abaxial surfaces were thereafter inoculated with 10 µl each of $10^6$ spores per mL of freshly harvested *C. gloeosporioides* inoculum amended with Tween 20, at 4 points. Controls were inoculated with only water amended with Tween 20. They were incubated under 12 h of fluorescent light and 12 h darkness in an incubator with a relative humidity of about 90% and temperature of 28 °C. The experimental layout was a completely randomized design with two replicates. Data on the disease severity was recorded using Leaf Doctor[82] at 7 Days After Inoculation (DAI), 14 DAI, and 21 DAI. We adopted a 1–5 scoring system (as used for field phenotyping) to classify the parental lines and progeny of the mapping population as highly resistant, resistant, moderately resistant, susceptible, and highly susceptible respectively. AUDPC was also calculated for data obtained from DLA.

**Anthracnose disease assessment at NRCRI**

At the National Root Crops Research Institute (NRCRI, Umudike, Nigeria), site-specific *C. gloeosporioides* isolates were first collected and evaluated, and the most virulent isolates were selected for conducting DLA on NRCRI mapping populations.

*Isolation and identification of C. gloeosporioides isolates*: We sampled anthracnose-infected yam leaves obtained from open fields at 24 yam farms at different locations in Umudike, Nigeria in 2017. The yam leaf samples showed a variety of anthracnose symptoms and samples were collected over a period of five months covering the whole duration of the yam growing season in Nigeria, to trap the different strains of the *C. gloeosporioides*. These leaf samples were surface sterilized in 20% sodium hypochlorite for 1 min and rinsed in four changes of sterile distilled water separately in beakers. The surface-sterilized yam leaf samples were cut into pieces of approximately 4 × 4 mm using surgical blades and forceps sterilized to red hot over flame under aseptic condition under laminar airflow hood. The cut pieces were cultured for fungal growth by plating four pieces on PDA in Petri dishes with spaces of approximately 5 mm away from each other. After growing the organism on PDA for 4–7 days, the organisms were subcultured for purification into pure isolates and identified under the microscope based on the morphology and growth pattern.

*C. gloeosporioides pathogenicity test*: Forty-six different *C. gloeosporioides* isolates were obtained and tested on detached leaves of four varieties of yams (*D. alata*) with known varying anthracnose resistance. Many of the isolates showed different levels of virulence but two were selected based on the highest vertical and horizontal virulence on the yams tested. Isolate Um-7 had the highest vertical virulence which did not differ significantly from isolate Um-11 which had significantly higher horizontal virulence than Um-7. The isolate Um-11 was used for DLA of the three sets of greater yam. Copies of the isolates were kept on slants in a 4 °C refrigerator for use at any time required time of evaluation.

Detached Leaf Assay (DLA): Anthracnose severity evaluation was carried out at NRCRI on *D.alata* mapcross progeny using the yam DLA protocol[81]. Young healthy yam leaves approximately three months old were cut from a growing yam plant. The detached leaves were surface sterilized using 20% sodium hypochlorite for 1 min and passed through four rinses of distilled water for 1 min each in a beaker. Excess water on the leaf sample was wiped off with a sterile paper towel. The detached leaves were placed on filter paper moistened with distilled water in a transparent plastic take-away plate and covered with the lid. These leaves were inoculated with a 30 µl drop of inoculum at a concentration of $1\times10^6$ spores per mL on the abaxial surface. Plates containing three replicates of the inoculated leaf samples were incubated at ambient temperature (+28 °C) and evaluated on days 4, 8, 12, and 16 after inoculation.

Anthracnose severity evaluation used the YAD scoring[81] scale of 1–5 as follows: score 1, 0.0% anthracnose symptom observed (rated as highly resistant [HR]); score 2, 0.1–25% anthracnose observed (moderately resistant [MR]); score 3, 25.1–50% anthracnose observed (resistant [R]); score 4, 50.1–75% anthracnose observed (susceptible, S); and score 5, 75.1–100% (highly susceptible, HS) anthracnose symptoms observed on the leaf.

All phenotyping data can be found in **Supplementary Data 5**.

#### Supplementary Note 6: Whole-genome ancestry reconstruction and population genetic analysis

##### WGS Illumina sequencing and variant calling

DNA isolations from the breeding lines listed in **Supplementary Table 1** were performed at IITA as for DNA used in mate-pair libraries.

Early DNA isolations at IITA from *D. alata* TDa01/00039 (male) and TDa00/00005 (female) samples were performed essentially in this manner, with some slight differences. Samples were collected using liquid nitrogen. For polysaccharide removal, tissue was first treated with a HEPES buffer (for 20 mL: 0.1 M HEPES buffer pH 8.0, 204 mg PVP, 180 mg L-ascorbic acid, 400 µl 2-mercaptoethanol). Just before use, the ingredients were mixed and the buffer was vortexed thoroughly. 1 mL of buffer was added to 100 mg (wet mass) ground plant tissue, mixed thoroughly, centrifuged for 5 min at about 8000 rpm, and the supernatant discarded. The washing step was repeated if necessary. DNA extraction was performed using the CTAB method.

TruSeq Illumina libraries were constructed and sequenced at the Vincent J. Coates Genomics Sequencing Laboratory at UC Berkeley. Inferred insert sizes ranged from 247–876 bp. These were sequenced on HiSeq 2500 or HiSeq 4000 with read lengths ranging from 150–251 bp, yielding combined sample depths of 19 to 230×. **Supplementary Data 1** lists all Illumina sequence data from our breeding lines, including external data, and accompanying summary statistics.

Dense single-nucleotide variants (SNVs) used to characterize the fine-scale genetic backgrounds of the founders of the mapping populations were extracted from Illumina whole-genome shotgun resequencing reads. For each sample, and for each sequencing run, Illumina TruSeq adapter fragments were removed with fastq-mcf (from the ea-utils[83] tool suite) v1.04.807-18-gbd148d4, and the trimmed sequences were aligned with BWA-MEM[14] v0.7.17-11-g20d0a13 to the TDa95/00328 v2 genome sequence containing complete *D. alata* plastid and mitochondrial sequences, as well as the *Pseudomonas synxantha* sequence (described above) included as bait for contaminating sequences, as all samples were grown at the same field site and *P. synxantha* sequences could similarly be present. Duplicate read pairs were removed using the 'fixmate' (-c -m) and 'markdup' subcommands included in SAMtools[32] v1.9-93-g0ca96a4. The BAM files for each sample were subsequently merged with SAMtools into a single sample-level BAM file, then filtered further to include only properly-paired (-f3 -F3852) reads for variant calling.

The filtered resequencing alignments were reduced to base-pair resolution genome variant call format (gVCF) files with the Genome Analysis ToolKit (GATK; v3.8-1-0-gf15c1c3ef) HaplotypeCaller (-ERC BP_RESOLUTION --min_base_quality_score 20 --min_mapping_quality_score 30 --heterozygosity 0.01 --indel_heterozygosity 0.001), and sample-level genotypes called with GATK's GenotypeGVCFs tool[84] (params: -G StandardAnnotation -G StandardHCAnnotation --includeNonVariantSites). Next, filters were implemented to ameliorate the effects of poor mappability and mismapping genomic repeat sequences, which result in false-positive variant calls and noise in downstream analyses. Genotypes were hard-filtered (*i.e.*, converted to "./.") if any of the following criteria were satisfied: 1) locus depth less than ~0.75 times the genome-wide median depth, or greater than 2.5 standard deviations above; 2) allele balance less than 0.30 or greater than 0.70 at heterozygous loci, or greater than 0.10 (or less than 0.90) for loci homozygous for the reference (alternate) allele; and 3) allele-balance binomial test $p$-value < 0.001 ($p_0$ = 0.5 for heterozygous loci, $p_0$ = 0.01 for homozygous loci) (**Supplementary Fig. 14**). The above filters were applied using BCFtools 1.9-213-g4411f1e and custom Python (v2.7.11 and v3.7.6) scripts[85,86].

Additionally, several interval masks were developed based on examining annotation and alignment tracks in the Integrative Genomics Viewer (IGV)[87]. The first mask imposed a secondary maximum-depth criterion calculated from all (unfiltered) read alignments and targeted regions of the genome where repetitive sequences were incompletely filtered by the first-round maximum-depth criterion. Windows of 100 bp were used to calculate the average depth across the genome; windows with a depth greater than 1.75 standard deviations above the mean were merged into a single, larger window if within 500 bp of another. A masking interval was discarded if its length was less than 150 bp (the typical length of a sequencing read). The resulting masks were then extended in each direction by 100 bp. The second and third masks leveraged the RepeatMasker annotations, run both with and without LTR_harvest/LTR_retriever, to mask annotated repeat sequences. Any loci intersecting either of these three masks were omitted using BEDtools[72] v2.28.0. Finally, the sample-level VCF files were merged into a multi-sample VCF with GATK's CombineVariants tool. Only biallelic SNVs were used in downstream analyses.

#### WGS population analyses

From the restrictive set of 1.89 million SNVs, initial pairwise estimates of genome-wide relatedness were obtained using the KING-robust[8] relatedness measure triggered by the '--relatedness2' option implemented by VCFtools[36] v0.1.16-16-g954e607. The first two numbers that follow the TDa in a breeding line's identifier indicate that it was selected from a population generated in that year, e.g., TDa95/00328 was selected in 1995. We used this year information to prepare the pedigree where needed to infer the directionality of relatedness, for example in a parent-child relationship we inferred the older line must have been the parent. Once IITA pedigrees and the relationships inferred from the sequencing data were established to be consistent, custom scripts ('IBD' script in https://bitbucket.org/rokhsar-lab/wgs-analysis) were used to infer and visualize their pairwise segmental identity-by-descent (IBD) (**Fig. 4**). Briefly, sliding windows of 5000 SNVs wide (with 1000 SNV step) were used to quantify the percent of genotypes sharing alleles identical by state (IBS). If two individuals shared ≥95% of their genotypes IBS, that genomic segment was called IBD for the same state; e.g., a window of 5000 loci with 98% of genotypes sharing both alleles (IBS2) implies two shared haplotypes (IBD2) across this segment. The IBD fractions, $f_{IBD0}$, $f_{IBD1}$, and $f_{IBD2}$ were calculated for each window using the following equations:

$$f_{IBD0} = \max \{ 0, 2 \cdot n_{IBS0} / (n_{het} + n_{hom} - n_{IBS1}) \}$$

$$f_{IBD1} = \max \{ 0, 1 - n_{IBS0} \}$$

$$f_{IBD2} = \max \{ 0, 2 \cdot (n_{IBS2} + n_{IBS2^*}) / (n_{het} + n_{hom} - n_{IBS1}) \}$$

Where $n_{IBS0}$, $n_{IBS1}$, and $n_{IBS2}$ represent the numbers of loci with genotypes sharing zero, one, and two alleles, respectively, and $n_{IBS2^*}$ is the number loci where both individuals compared are heterozygous. The variables $n_{het}$ and $n_{hom}$ represent the numbers of loci with heterozygous and homozygous genotypes in either individual.

The intrinsic heterozygosity (i.e., SNV rate) and homozygosity were estimated for each individual as the number of heterozygous (or homozygous) biallelic SNVs per kilobase of callable loci. These estimates were calculated using variable-sized sliding windows to sample 100 kb of loci (with a 10 kb step) that passed all of the locus- and genotype-level filters described above ('snvrate' script at https://bitbucket.org/rokhsar-lab/wgs-analysis). A window was deemed homozygous (i.e., autozygous) if the rate of heterozygous genotypes in the window was less than $2 \times 10^{-4}$. This threshold was determined empirically (**Supplementary Fig. 3**).

#### Supplementary Note 7: Mitochondrial and plastid sequence assemblies

*D. rotundata* mitochondrial fragments and plastid sequences for previously published *Dioscorea* species and outgroups (**Supplementary Table 7**) were downloaded from NCBI, concatenated to an early version of the TDa95/00328 genome assembly, and used as target sequences to bait PacBio reads via Minimap2[44] v2.5-284-g1739a26 alignment. Reads aligning to plastid sequences were collected and assembled into contigs with Canu v1.7-221-gb5bffcf. This process was iterated until the number of assembled contigs converged to a single value, meaning that the assembled sequences could be extended no further. The assembled *D. alata* mitochondrial sequence fragments were then scaffolded manually in JuiceBox. The mitochondrial scaffold and complete plastid were polished using short paired-read data, as was performed for the nuclear genome.

The *D. dumetorum* plastid sequence deposited in NCBI (**Supplementary Table 7**) was used to search the IboSweet3 *D. dumetorum* assembly using Minimap2. The IboSweet3 plastid was found to be represented as a single contig (contig888) at full length, flanked on either side by an additional incomplete copy. The full-length sequence was extracted and polished using short paired-read data (SRA: ERR3611139 and ERR3611140), as described above.

For other breeding lines in this study, for which only short-read data were available, plastid sequences were aligned to the previously published *D. alata* plastid sequence (MG267382.1) with BWA-MEM[14] v0.7.17-11-g20d0a13 and the allelic state of each locus was called at BP_RESOLUTION as gVCF output with GATK HaplotypeCaller in haploid mode. gVCF files were genotyped jointly using the GenotypeGVCFs[84], which were then converted to FASTA format with GATK's FastaAlternateReferenceMaker tool.

All assembled plastid sequences were manually trimmed, recentered, and their replication origins re-oriented to conform to standard convention. Nucmer, from the MUMmer[88] suite (v3.23), was used to perform alignment and visualization, and BEDtools[72] v2.28.0 and custom scripts were used for sequence manipulations.
